# Biophysical cartography of the native and human-engineered antibody landscapes quantifies the plasticity of antibody developability

**DOI:** 10.1101/2023.10.26.563958

**Authors:** Habib Bashour, Eva Smorodina, Matteo Pariset, Jahn Zhong, Rahmad Akbar, Maria Chernigovskaya, Khang Lê Quý, Igor Snapkov, Puneet Rawat, Konrad Krawczyk, Geir Kjetil Sandve, Jose Gutierrez-Marcos, Daniel Nakhaee-Zadeh Gutierrez, Jan Terje Andersen, Victor Greiff

## Abstract

Designing effective monoclonal antibody (mAb) therapeutics faces a multi-parameter optimization challenge known as “developability”, which reflects an antibody’s ability to progress through development stages based on its physicochemical properties. While natural antibodies may provide valuable guidance for mAb selection, we lack a comprehensive understanding of natural developability parameter (DP) plasticity (redundancy, predictability, sensitivity) and how the DP landscapes of human-engineered and natural antibodies relate to one another. These gaps hinder fundamental developability profile cartography. To chart natural and engineered DP landscapes, we computed 40 sequence- and 46 structure-based DPs of over two million native and human-engineered single-chain antibody sequences. We found lower redundancy among structure-based compared to sequence-based DPs. Sequence DP sensitivity to single amino acid substitutions varied by antibody region and DP, and structure DP values varied across the conformational ensemble of antibody structures. Sequence DPs were more predictable than structure-based ones across different machine-learning tasks and embeddings, indicating a constrained sequence-based design space. Human-engineered antibodies were localized within the developability and sequence landscapes of natural antibodies, suggesting that human-engineered antibodies explore mere subspaces of the natural one. Our work quantifies the plasticity of antibody developability, providing a fundamental resource for multi-parameter therapeutic mAb design.

## Introduction

Monoclonal antibodies (mAbs) are widely used therapeutics against cancer, autoimmune, and infectious diseases (*1–5*). The global mAb market is forecasted to grow to more than $300 billion in 2025 (*6*). Despite their commercial success, mAb discovery remains a resource- and time-consuming process resulting in a costly and lengthy clinical approval, hindering their accessibility and affordability (*7*, *8*). A successful mAb molecule should not only show sufficient affinity in its target binding profile but, ideally, also exhibit a desirable “developability” profile (*9*). The term “developability” refers to a combination of intrinsic physicochemical parameters defined as developability parameters (DPs) that relate to biophysical aspects of antibodies and their formulations – including aggregation, solubility, and stability (*10–14*). The feasibility of an antibody candidate to successfully progress from discovery to development is underpinned by specific DPs, which reflect its manufacturability and druggability (*10*, *15*). Thus, suboptimal developability is one of the main factors of mAbs failure in preclinical and clinical development stages (*16–18*). Therefore, the ability to predict and prospectively design developability properties, in line with clinical and manufacturing requirements, would help by reducing the time and resources invested in the development of therapeutic mAbs, thus, boosting their success rate (*19–21*).

Traditionally, developability screening is performed through a series of laborious *in vitro* assays (*17*, *22*, *23*). Therefore, major efforts have been invested into developing real-world-relevant *in silico* tools and machine learning (ML) algorithms that can computationally quantify or predict DP values using antibody sequence and/or structure information (*7*, *24–35*). Given the current high throughput of antibody structure prediction at repertoire scale (*36–38*), tools for computational developability determination have become available, which can be used to identify potential design shortfalls during development and selection of lead mAb candidates (*39–42*).

In contrast to design and selection of pharmaceutical mAbs, the natural (or native, used interchangeably) immune system has the capacity to “design” antigen-specific antibodies with physiologically compatible and optimized biochemical properties within days to weeks (*43–46*). Indeed, it was previously reported that antibodies obtained via, for example, humanized mice exhibit far fewer developability risks compared to mAb candidates obtained from *in vitro* display campaigns (*4*, *17*, *20*, *47*). In line with these findings, clinical mAbs have been reported to exhibit high sequence identity matches (>70%) with natural antibodies for both heavy and light chains, implying that artificially developed mAbs are not entirely dissimilar from their native counterparts, thus, highlighting the relevance of mining the natural antibody repertoires for therapeutic mAb discovery (*48*, *49*). These findings motivate the embedding of therapeutic mAb sequences in the developability landscape (here defined as multidimensional distribution of DP parameter values across DPs and antibodies) of the natural (human and mouse) antibody repertoires to assess the nativeness of their developability profiles (DPLs, a DPL is a set of DP values for a given antibody sequence/structure) (*50*). As such, a mAb candidate with a DPL falling outside the range of its variation in natural antibodies may be assumed unnatural and, therefore, more likely to exhibit undesirable *in vivo* characteristics (*49*). Recent studies have also pointed to the futility of attempting complete separation between natural antibodies and therapeutic mAbs based solely on DP values (*51*). Similar findings continue to emphasize the valuable knowledge that can be harnessed from interrogating the growing sequence space of naturally sourced antibody sequences to accelerate the engineering and optimization of mAb candidates (*2*, *48*).

So far, the relationship between the developability landscape of the natural antibody repertoire and that of therapeutic (or, more broadly, human-engineered) antibodies remains unclear. While not all natural antibodies may be suitable as therapeutic candidates from a developability perspective (*7*, *51*), we lack a large-scale overview of sequence and structure-based natural antibody developability landscapes stratified by antibody isotype and species. Furthermore, given previous low-sample size investigations, we are unaware to what extent sequence changes affect a given DP and which sequence or structure-based design restrictions may limit all-vs-all DP optimization with a given antibody sequence. Furthermore, many studies have focused on extracting developability guidelines from a limited number of successful mAbs, considering them as a “ground truth” of desired developability (*7*, *10*, *17*, *29*, *52*). In addition, most studies have focused on a small number of DPs (*17*, *53*), and apply their hypotheses to limited antibody datasets comprising of a few 100s to 1000s antibody sequences (*7*, *10*, *34*, *52*) or datasets not including patent-submitted antibodies or antibodies that failed during early clinical trials (*7*, *10*, *54*). So far, the lack of datasets with sufficient sample size has hindered an in-depth understanding of the plasticity of the antibody developability space.

Understanding the natural antibody landscape could enable the integration of both current and prospective antibody therapeutic candidates, which will improve our interpretation of the disparities in developability between human-engineered mAbs and natural antibodies (Figure 1). To this aim, we have built an atlas of over two million unique native antibody sequences from human and murine heavy and light chains (≈200,000 per isotype and chain) annotated with DPs. We predicted the 3D structure of all antibodies and calculated 40 sequence-based and 46 structure-based DPs for each antibody. Using correlation and graph theory, we identified a subset of DPs that are maximally different from one another, thus delineating a non-redundant multidimensional antibody developability space. Across all antibody isotypes, we found lower interdependency among structure-based DPs in contrast to sequence-based DPs. Notably, distinct developability landscapes emerged across species (mouse, human) and antibody chains (heavy, light). In addition, we quantified DP sensitivity by analyzing the DP value distribution of mutants with single amino acid substitutions. We also found that the values of DPs measured on the conformational ensembles of antibodies evolve throughout their molecular dynamics (MD). Regarding predictability, our analysis revealed that ML is more successful in predicting the values of sequence-based DPs, indicating a less confined design landscape for structure-based DPs. Our analysis also suggested that the observed developability spaces of human-engineered antibodies are essentially subsets of the broader natural developability space (in terms of the major principal components of variation). While our study relies on computationally predicted developability, in which experimental correspondence may vary (*15*), it serves as a practical and real-world relevant use case of integrating and charting repertoire-wide developability to guide mAb selection and development.

**Figure 1.**
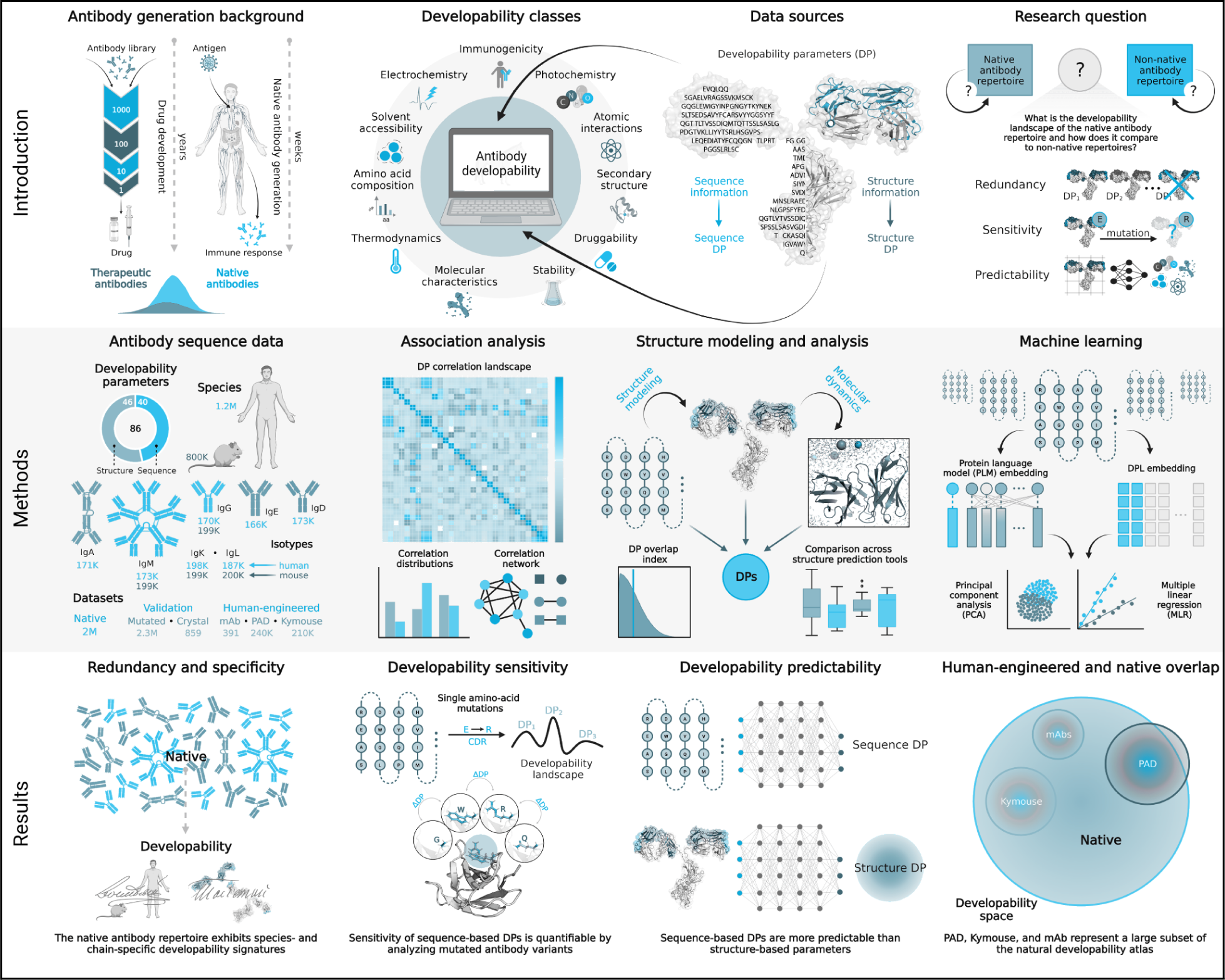
Graphical abstract: redundancy, sensitivity, and predictability of developability parameters in native and human-engineered antibodies. <u>Introduction</u>: The development of therapeutic mAbs takes years, and DPs dictate the selection and design of candidates for (pre-)clinical testing. Here, we analyzed the plasticity of the developability landscapes of natural antibodies in terms of DP redundancy (extent of DP intercorrelation), sensitivity (extent of DP change as a function of antibody sequence change), and predictability (predictability of a given DP based on one or several DPs). <u>Methods</u>: To analyze the constraints on natural antibody developability and to relate these to current human-engineered antibody datasets, we assembled a dataset of over 2M native antibody sequences (heavy and light chain isotypes, human and murine) and computed 40 sequence- and 46 structure-based DPs. To reduce redundancy, we determined the minimum-weight dominating sets (MWDS) of DP correlation networks. To quantify sensitivity, we analyzed single-amino-acid substituted variants followed by characterization of the impact of sequence variation on DP distribution. To compute predictability and assess the interdependence of DPs, we trained multiple linear regression (MLR) using developability profile (DPL) and protein language model (PLM) embeddings. These embeddings were used to relate native antibodies to human-engineered ones via principal component analysis (PCA). Moreover, we performed classical molecular dynamics simulations to analyze the distributions of antibody DP values and define how the rigid models fit into these distributions. <u>Results</u>: Our results address all three research areas (redundancy, sensitivity, and predictability). Redundancy: We found a lower degree of interdependence among structure DPs compared to the sequence-based ones for all isotypes of the native dataset, and higher pairwise antibody sequence similarity was not always associated with higher pairwise antibody developability similarity. Native antibody datasets contained species- and chain-specific developability signatures. Sensitivity: We propose methods to quantify the sensitivity of antibody DPs to minimal sequence changes. Predictability: We found that structure-based DPs are less predictable than sequence-based DPs using protein language model (PLM) and multiple linear regression (MLR) embeddings. The comparison between native and human-engineered datasets revealed that human-engineered (therapeutic, patented, and Kymouse) datasets were localized within the native developability landscape.

## Results

### Sequence-based DPs show higher association and redundancies compared to structure-based DPs

The diversity and redundancy of the natural DP landscape of human and murine antibody repertoires has not been investigated. To do so, we assembled a dataset of ∼2M non-paired V_H_ and V_L_ native antibody sequences (∼170K–200K sequences for each isotype, human; IgD, IgM, IgG, IgA, IgK, and IgL, murine; IgM, IgG, IgK - Supp. Figure 1A). Next, we computed 40 sequence-based and 46 structure-based developability parameters (Supp. Table 1) for each antibody after having predicted their 3D structures using ABodyBuilder (ABB) (*36*). The choice of these DPs was intended to cover a comprehensive array of the physicochemical properties of antibodies based on our understanding of the developability literature and antibody structure. This includes categorical amino acid composition, electro- and photo-chemical parameters, as well as structural interactions and conformational descriptors (among others detailed in Supp. Table 1, Supp. Table 2).

Prior to the DP analysis, we controlled for the quality of the computational data generated along two axes (Supplemental File). Briefly, (1) given that the majority of the work was performed on unpaired chain data, due to a lack of isotype-stratified paired-chain data, we verified that structure-based DPs measured on a subset of paired-chain antibodies strongly correlate with the corresponding DP values measured on each of the unpaired (heavy and light) single chains separately (median Pearson correlation 0.84–0.9, Supp. Figure 16). (2) We also analyzed, using molecular dynamics (MD), the dependence of structure-based DPs on computational antibody structure prediction methods (such as IgFold (*55*) and AbodyBuilder2 (*37*). We found that the predominant structure prediction method used in this study (AbodyBuilder (*36*)) faithfully replicated conformations within the antibody structure conformational ensemble of a given antibody sequence as determined by MD (Supp. Figure 18), thus validating our strategy to compare antibody developability landscapes.

First, we inquired about the degree of correlation among different DPs within the native antibody dataset. To address this, we examined the pairwise correlation among the values of DPs by isotype and species for the full native dataset (Figure 2A). Overall, we found that sequence-based parameters were significantly more correlated with one another compared to structure DPs across all isotypes of the native dataset, suggesting greater general association among sequence DPs (median absolute Pearson correlation coefficient 0.14–0.21 for sequence-based DPs, 0 for stricture-based DPs, Figure 2A). Although the degree of correlation for both sequence and structure DPs was relatively low (median absolute Pearson correlation ≤0.21), sequence-based parameters formed larger correlation clusters with high correlation values (>0.6) in the native human IgG dataset (Figure 2B; boxed in black). Similar clustering patterns with high correlation values were found across non-IgG isotypes for the sequence-based DPs (Supp. Figure 2, Supp. Figure 3). As a control measure, we repeated this analysis on permuted DPs and reported a drastic loss of correlation and a significant change in the distribution of Pearson correlation coefficient values (Supp. Figure 5A,B).

**Figure 2.**
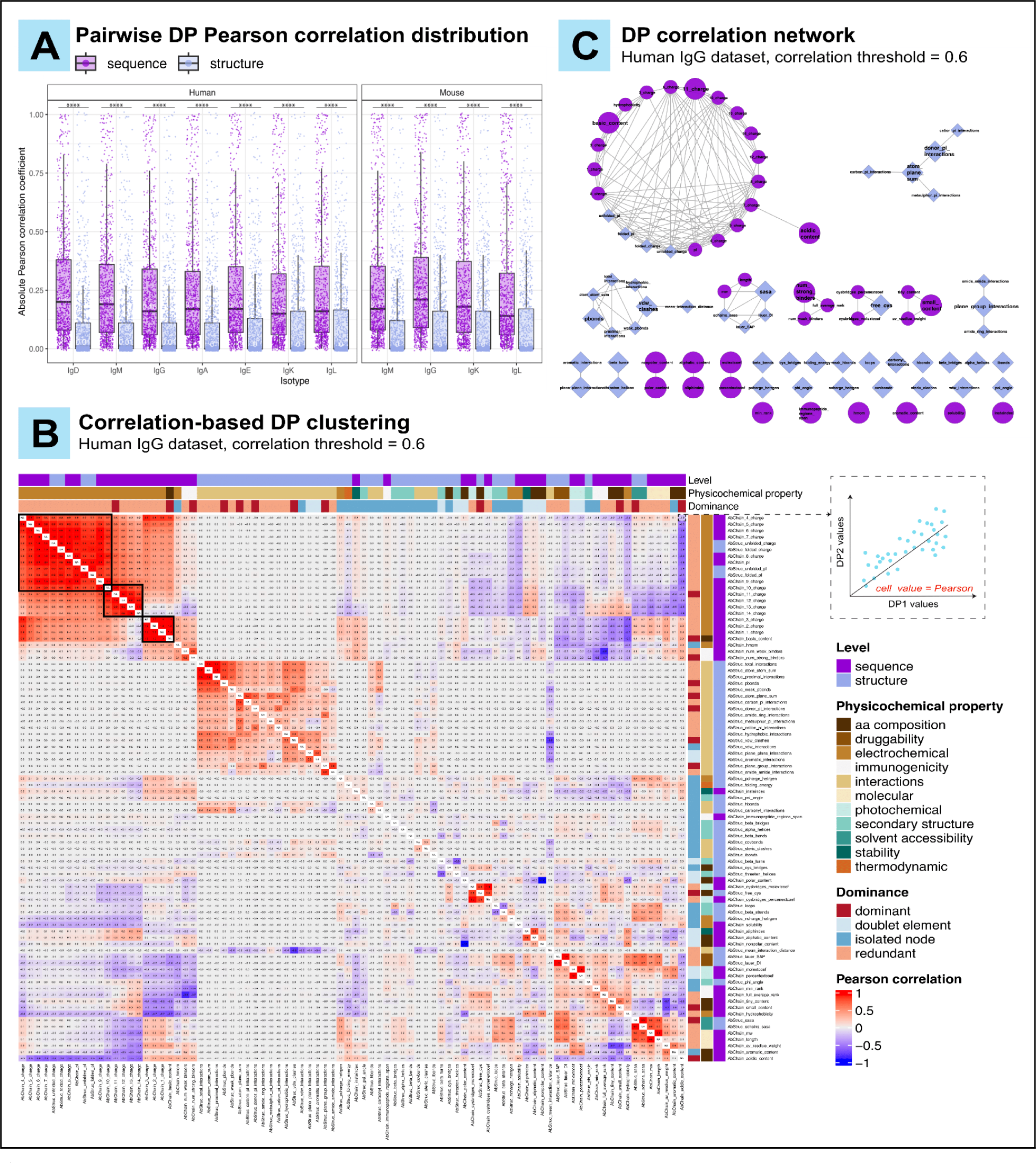
Sequence-based developability parameters show higher redundancies compared to structure-based parameters. Absolute pairwise Pearson correlation of sequence and structure developability parameters within the native antibody dataset. Numerical values on the figure represent the median of Pearson correlation for the corresponding subset. Differences were assessed using pairwise Mann-Whitney test with p-value adjustment (Benjamini-Hochberg method). **** p < 0.0001 (Human; IgD: 2.09e^-112^, IgM: 6.54e^-104^, IgG: 5.21e^-100^, IgA: 8.94e^-101^, IgE: 1.04e^-94^, IgK: 4.61e^-74^, IgL: 7.46e^-80^, Mouse; IgM: 1.276e^-98^, IgG: 4.76e^-101^, IgK: 6.9e^-88^, IgL: 6.68e^-71^). **(B)** Hierarchical clustering of 40 sequence and 46 structure developability parameters based on pairwise Pearson correlation for 170,473 IgG human antibodies (median of absolute Pearson correlation: 0.02 +/-0.003 SEM). As explained in the inset (top right), each cell within the heatmap reflects the value of Pearson correlation for a pair of DPs. Developability parameters are color-annotated with their corresponding level (sequence or structure), physicochemical property (as detailed in Supp. Table 1), and dominance status from the ABC-EDA algorithm output at Pearson correlation coefficient threshold of 0.6 (see Methods). Black boxes highlight correlation clusters that contain more than three DPs and exhibit pairwise Pearson correlation coefficient > 0.6 **(C)** An undirected network graph of the pairwise parameter Pearson correlation data from with nodes representing DPs. An edge was drawn between a pair of nodes when the absolute pairwise Pearson correlation coefficient was > 0.6 (see Methods). Circular (dark orchid) nodes represent sequence DPs and square (cloudy blue) nodes represent structure ones. The size of the node reflects its dominance state (large: dominant, small: redundant) as determined by the ABC-EDA output. Out of the 86 total parameters, 10 were classified as doublets, 23 as isolated nodes and 12 as dominant parameters for the human IgG repertoire. Structure-based developability parameters are majorly present as isolated nodes due to the lack of correlation with sequence parameters and among themselves (pairwise Pearson correlation mostly lower than 0.6), whereas sequence-based parameters are mainly present in larger subnetworks as they are more correlated. **Supplementary Figures:** Supp. Figure 2, Supp. Figure 3, Supp. Figure 4A.

Subsequently, we asked which DPs are redundant, as quantified by the previous analysis. To answer this question, we utilized the pairwise Pearson correlation coefficient values from Figure 2A,B to construct undirected weighted network graphs (Figure 2C). Additionally, we employed the hybrid artificial bee colony – estimation of distribution hybrid algorithm (ABC-EDA, (*56*)) to identify the minimum weighted dominating set (MWDS) among DPs (see Methods). In this context, the MWDS comprises the most uncorrelated DPs that sufficiently reflect the overall developability for a set of antibodies at a given Pearson correlation threshold (*56*). When we conducted this analysis on the native IgG repertoire at an absolute correlation threshold of 0.6, we found that the pairwise relationships among DPs can form subnetworks, doublets, and isolated nodes (Figure 2B,C). At this threshold, subnetworks were largely formed by DPs that reflect similar physicochemical properties regardless of their level (sequence or structure – Supp. Table 1). For instance, sequence- and structure-based charge and isoelectric point (electrochemical) DPs (n=20) clustered with the acidic and basic amino acid composition DPs (Figure 2C). The largest subnetwork was predominantly occupied by sequence DPs. Meanwhile, structure-based DPs mainly formed isolated nodes (n=17) and smaller subnetworks (Figure 2C).

When we repeated this analysis on all the isotypes of the native dataset (Supp. Figure 1A), we found that the proportion of isolated nodes to the initial DP count was consistently higher among structure-based DPs, starting from low correlation thresholds (0.1–0.2) across all isotypes, in comparison to sequence-based DPs (Supp. Figure 4A). For example, only 12.5%–22.5% of sequence-based DPs (5–9 out of 40) compared to 26.1%–41.3% of structure-based DPs (12–19 out of 46) were classified as isolated nodes at a Pearson correlation threshold of 0.6 (Supp. Figure 4A). Meanwhile, we found that the proportion of subnetwork dominant DPs was higher among sequence DPs across all isotypes for the higher correlation thresholds (Pearson correlation > 0.6). For example, 7.5%–12.5% of sequence-based DPs were categorized as dominant nodes at the strictest correlation threshold (0.9) in comparison to structure-based DPs (2.2% – 6.5%) (Supp. Figure 4A). Collectively, these results emphasize the lower interdependence among structure-based DPs when compared to the sequence-based counterparts.

### The native antibody dataset exhibits chain-type and species-specific developability signatures

Next, we asked to what extent the isotypes of the native datasets are similar to one another in regards to DP redundancies (MWDS parameters) and associations (DP pairwise correlations). In relation to these driving questions, we also asked to what extent natural antibodies harbor chain (V_H_,V_L_) and species-specific (human, mouse) DP differences. To address these questions, we first investigated the similarities in parameter redundancies among the native dataset for a given Pearson correlation value (0.6). Specifically, we explored the pairwise intersection size of the MWDS parameters on both sequence and structure levels for the human and murine antibody datasets, featuring the common isotypes (IgM and IgG) between the two species in our dataset (Figure 3A), and all the isotypes of the human V_H_ antibodies (Supp. Figure 4B).

**Figure 3.**
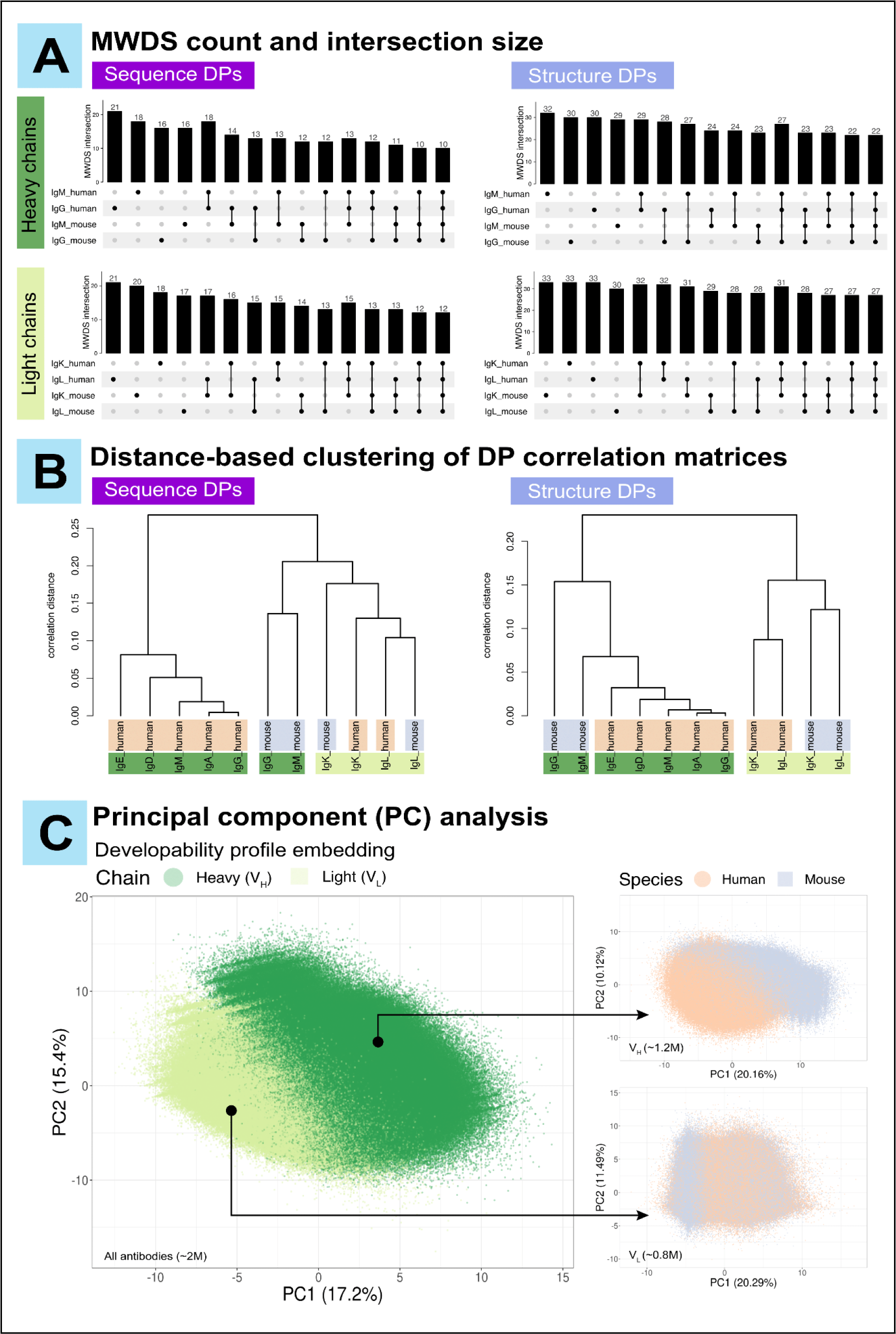
The native (human and murine) antibody datasets exhibit chain-specific and species-specific developability signatures. (A) MWDS intersection size for the human and mouse native datasets. Numerical values on the figure reflect the MWDS count (for an individual subset) and intersection size (for more than one subset). The MWDS for the respective isotypes was identified using the ABC-EDA algorithm (see Methods) at a threshold of absolute Pearson correlation of 0.6. For MWDS intersection size among all human heavy chain subsets, please refer to Supp. Figure 4B. **(B)** Distance-based hierarchical clustering of isotype-specific pairwise DP correlation matrices (sequence and structure levels). The height of the dendrograms (shown to the left of the dendrograms) represents the correlation distance among the dendrogram tips. **(C)** Repertoire-wide principal component analysis (PCA) of the native antibody developability profiles. We performed this analysis for the complete native dataset (left pane; ∼2M data points) and for the chain-specific datasets (right panels; ∼1.2M data points in the top panel, ∼0.8M data points in the bottom panel). The dimensionality of complex developability profiles was reduced to 2D PCA projections. The full value distribution of the corresponding PCs associated with each projection is shown in Supp. Figure 4C. **Supplementary Figures:** Supp. Figure 4B,C.

This analysis revealed that both heavy (IgG and IgM) and light chain human antibody datasets displayed a larger MWDS intersection size on both sequence and structure levels in comparison to the murine counterparts (Figure 3A). For instance, human IgM and IgG datasets shared 18 sequence DPs (86% overlap) and 29 structure DPs (90% overlap) in their MWDS sets, whereas the same heavy chain isotypes of the murine dataset shared only 12 sequence DPs (75% overlap) and 23 structure DPs (77% overlap) (Figure 3A). Moreover, the human heavy chain dataset displayed greater or comparative MWDS intersection size even when considering all five antibody isotypes (71% overlap on sequence level, 84% overlap on structure level) in comparison to the murine heavy chain dataset (IgM and IgG only) (Figure 3A and Supp. Figure 4B). Similarly, the MWDS overlap for the human light chains (IgK and IgL) was greater on both levels (15 DPs – 71% overlap on sequence level and 32 DPs – 97% overlap on structure level) in comparison to the mouse light chain dataset (14 sequence DPs – 67% overlap and 29 structure DPs – 88% overlap) (Figure 3A). Thus, our findings suggest greater consistency among the isotypes in the human antibody dataset when it comes to DP redundancies (MWDS overlap), as opposed to the mouse antibody dataset.

Second, we sought to investigate the similarities in DP associations among the isotypes of the native antibody dataset. To this end, we clustered the isotype-specific datasets based on the distance of their pairwise DP correlation matrices (see Methods). This analysis revealed chain-specific segregation (heavy and light) and, within a given chain, species-specific segregation (human and mouse) of antibody subsets on the structural level (Figure 3B – right panel). Additionally, the human dataset showed a closer distance among its isotypes within the heavy and light chain clusters (0.03 for heavy chain isotypes, 0.08 for light chain isotypes) in comparison to the mouse dataset (0.15 and 0.12 for heavy and light chain clusters, respectively). An equivalent sequence-based analysis (Figure 3B – left panel) drew a similar conclusion regarding the uniqueness of chain-type developability. However, interspecies isotype-specific clustering occurred among the light chain subsets (Figure 3B). Similarly to the structure-based DP association clustering, the human heavy chain isotypes showed a smaller distance among themselves (0.08) in comparison to the murine IgM and IgG subsets (0.14) (Figure 3B). These findings suggest that native (human and murine) datasets harbor chain-type-specific developability signatures. Species-specific developability differences were less pronounced, especially for the light-chain antibody subsets.

The aforementioned clustering of antibody datasets based on DP associations led us to investigate whether the antibody species and chain type are key sources of variance in DP values. To this end, we performed a dimensionality reduction analysis on the developability profiles of the native antibodies using a principal component (PC) analysis (PCA) (Figure 3C). Examining the 2D PCA projections of developability profiles of all native antibodies (∼2M) further emphasized differences in antibody developability by antibody chain type (Figure 3C). In fact, the axes of maximal variance (PC1 and PC2) separated antibody sequences by V_H_ and V_L_ chain (absolute differences of medians = 5.6 and 1.8, respectively – Supp. Figure 4C). A similar projection of each chain type subset (heavy; ∼1.2M, light; ∼0.8M) allowed for (partial) species-based distinction of antibodies (Figure 3C). The influence of the antibody species on developability was more prominent among the heavy chain antibodies (absolute difference of PC1 medians = 4.1) in comparison to the light chain ones (absolute difference of PC1 medians = 3.2) (Supp. Figure 4C).

In summary, the isotypes of the human native dataset exhibit greater pairwise relatedness in regard to their DP associations and redundancies when compared to the murine dataset. Moreover, the antibody chain type and species of origin are significant factors influencing its overall developability.

### The sensitivity of sequence-based developability parameters is quantifiable by single-amino acid substitution analysis

Future antibody design will be performed in a multi-objective manner (*3*, *57*), which means that the design approach is to optimize a multitude of parameters at once. In certain cases, introducing minor changes might be sufficient to improve the value of a certain developability parameter. However, improving one parameter may result in compromising another (*3*, *58*). Therefore, it is of interest to understand to what extent minor sequence changes impact DP values. To address this question, we performed a single-substitution sensitivity analysis of DPs by quantifying the changes in DP value induced by one amino acid alterations in the antibody sequence at a time (see Methods, Figure 4A). Since there are no established methods and metrics to quantify sensitivity of antibody developability parameters (*59*), we employed two proxy measures to estimate DP sensitivity. First, we define the *average sensitivity* by the kurtosis, and secondly, the *potential sensitivity* as the range of a DP distribution of an antibody and all its possible single amino acid substituted variants (see Methods). Given the inability of current antibody structure tools, both template-based or de novo deep learning-based, to resolve differences between structures of antibodies with single amino acid variations (Supp. Figure 19 in Supplemental File), we focused our analysis on sequence-based DPs only.

**Figure 4.**
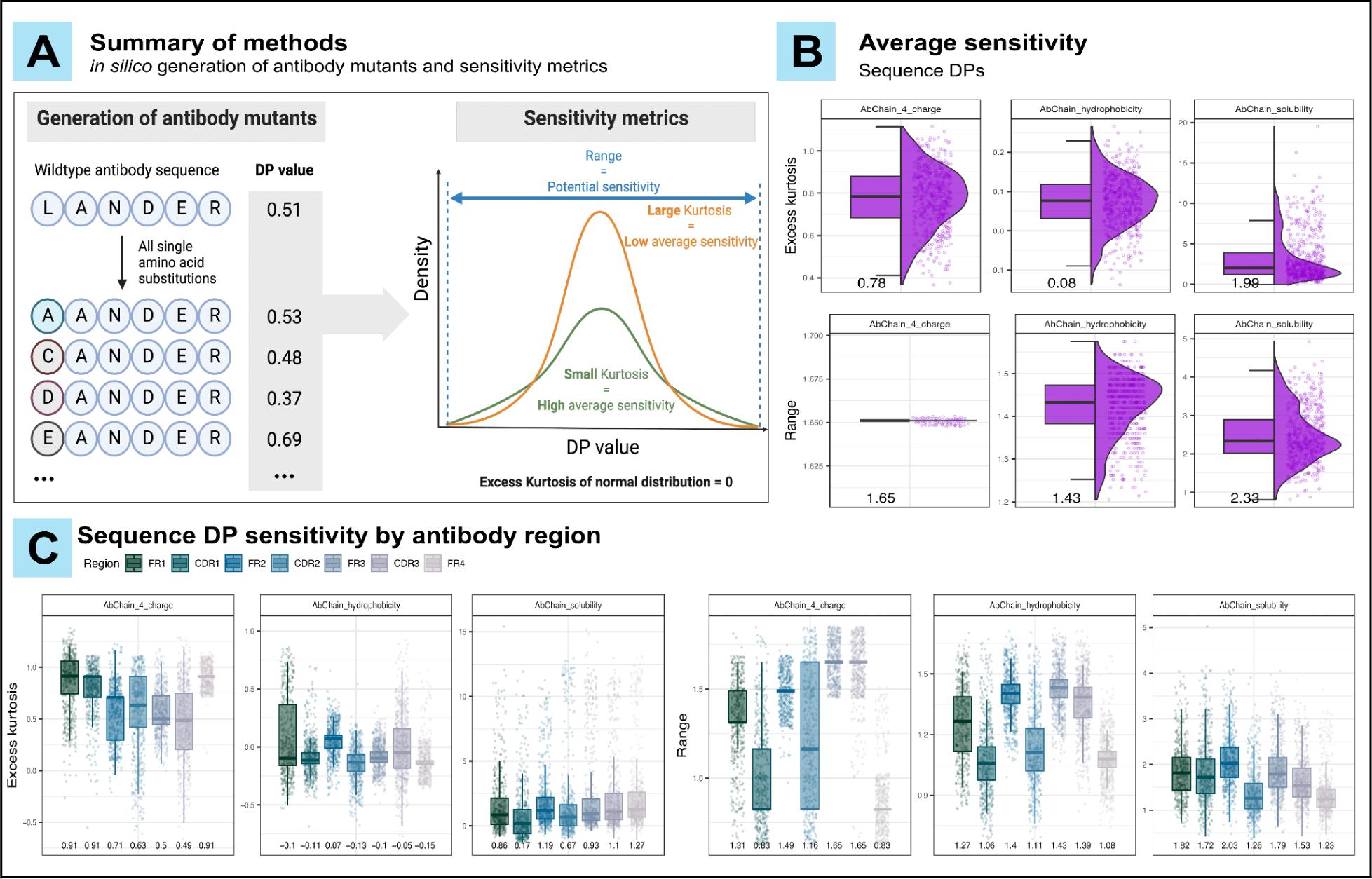
Developability parameter sensitivity can be quantified by analyzing mutated variants of wildtype antibodies. (A) DP values were computed for all possible single amino acid substituted mutants of 500 sampled wildtype human V_H_ antibody sequences (100 sequences sampled per isotype; 301,777 mutants in total). Each DP was scaled and mean-centered. The sensitivity was quantified for each DP by analyzing the DP dispersion of the mutants from their corresponding wildtype. Average sensitivity was measured by excess kurtosis (low kurtosis = high average sensitivity), while potential sensitivity was measured by the range (see Methods). **(B)** Average and potential sensitivity of selected sequence-based DPs. **(C)** Average and potential sensitivity of DPs from (B) grouped by antibody region in which the mutation occurred. In both (B) and (C), numerical values on the x-axis represent the median of the corresponding sensitivity metric. **Supplementary Figures:** Supp. Figure 12.

We linked the dispersion of a given DP value distribution (from mutated variants and their corresponding sampled wildtype) to its sensitivity and employed two metrics to quantify this dispersion (see Methods). The first metric, excess kurtosis (*60*), was implemented as a proxy measure for the *average sensitivity.* It describes how far the “tailedness” of a given distribution deviates from that of a Gaussian distribution (i.e., excess kurtosis = 0, Figure 4A). In this context, a positive excess kurtosis indicates a low average sensitivity, and a negative excess kurtosis implies a higher sensitivity with an increased proportion of mutant DP values diverging from the wildtype. Strictly, this is only valid under the assumption of a bell-shaped distribution and should be considered when estimating sensitivity by excess kurtosis.

The second metric is the range, which is the absolute difference between the smallest and largest values of a distribution after normalization. It was used as a proxy measure for the *potential sensitivity* of DPs as it reflects the potential extreme changes that can be introduced on DP values as a factor of amino acid sequence change (Figure 4A). Since only single amino acid substitutions were analyzed, we removed all DPs describing categorical amino acid composition (Supp. Table 1) where it is trivial to predict the changes in DP values, as well as the length of the sequence (AbChain_length) since it is not affected by substitutions. Because DPs denoting a sequence’s charge at a given pH were clustered in three highly correlated clusters (Figure 2B), we chose to retain only three of them which represent acidic, neutral and basic pH (AbChain_4_charge, AbChain_7_charge and AbChain_12_charge respectively). Additionally, all DPs with ‘_percentextcoef’-suffix were analyzed instead of their highly correlated counterparts with ‘_molextcoef’-suffix (Supp. Table 1).

We found the median excess kurtosis of most sequence-based DPs was > 0 and that most DPs exhibit an *average sensitivity* close to that of a normal distribution (excess kurtosis of a normal distribution = 0, Figure 4A, Supp. Figure 12A). In fact, some parameters, such as the molar extinction coefficient (of cysteine bridges) and the hydrophobic moment, were insensitive on average to substitutions as indicated by their high kurtosis (median excess kurtosis: 6.7, 49.02, and 29.49, respectively; Supp. Figure 12). Notably, none of the tested DPs displayed high average sensitivity (median excess kurtosis << 0, Figure 4A, Supp. Figure 12), suggesting that developability is relatively stable to the average single amino acid mutation with few outliers. Nevertheless, since DP values were normalized, small relative shifts induced by a mutation may still have a large effect in practice.

Next, we show that the median range in sequence-based DPs varied between 0.38 for the molecular weight DP (AbChain_mw) and 3.82 for the hydrophobic moment DP (AbChain_hmom – Supp. Figure 12B). Notably, some DPs (such as AbChain_4_charge – Figure 4B first column second row) have a constant or close to constant range, likely due to the fact that the set of all possible single amino acid substitutions covers the entire range of possible DP values. For example, when considering the charge of an antibody, the lowest possible charge results from substituting the most positively charged amino acid with the most negatively charged amino acid and vice versa. Since all amino acids are present in close to all sampled wildtype sequences, their ranges are almost identical as well. In DPs that depend on non-linear amino acid interactions, such as solubility (AbChain_solubility), the range was more diverse (2.33, Figure 4B).

To investigate the impact of substitutions across antibody regions, we grouped DP values of mutants by the region in which the substitution occurred and calculated their sensitivity metrics separately. We observed that electrochemical DPs such as charge and hydrophobicity (AbChain_4_charge and AbChain_hydrophobicity) exhibited higher potential sensitivity (range) in CDR3 and framework regions (median ranges AbChain_4_charge and AbChain_hydrophobicity respectively; CDR3: 1.65, 1.39, FR1: 1.31, 1.27, FR2: 1.49, 1.4 and FR3: 1.65, 1.43) compared to CDR1 (0.83, 1.06) and CDR2 (1.16, 1.11) and FR4 (0.83, 1.06; Figure 4C). Although we found the average sensitivity (excess kurtosis) to differ by antibody region as well (Figure 4C), there was no apparent general rule separating framework regions and CDRs.

In summary, our sensitivity analysis suggests that most DPs are comparably ‘normally’ sensitive (close to 0 excess kurtosis: mutant DP distribution as ‘tailed’ as a Gaussian distribution), and some parameters are especially *insensitive* to the average substitution. Additionally, although average and potential sensitivity differ by antibody region in which a mutation occurs, the differences are not generalizable across DPs.

### Antibody sequence similarity does not imply antibody developability similarity

Given that the values of DPs were prone to change with minor sequence changes, we asked to what extent pairwise sequence similarity is related to pairwise developability profile similarity, where the developability profile (DPL) was defined as a numerical vector that carries (sequence and/or structure) DP values in a fixed order for a given antibody sequence (see Methods).

To this end, we first examined the pairwise correlation of antibody developability profiles (<u>d</u>evelopability <u>p</u>rofile <u>c</u>orrelation: DPC) alongside the pairwise sequence similarity score (see Methods) for a random sample of 100 natural antibodies from the human IgM dataset that share the IGHV gene family annotation (Figure 5A). This is to eliminate the role of the V-gene as a factor of variance in our analysis, as up to 80% of sequence similarity can be expected among antibodies that belong to the same IGHV gene family (*61*, *62*). Sequence-level DPC clusters were often, but not always, accompanied by sequence similarity clusters, while structure-level DPC clusters were independent in regards to sequence similarity clusters (Figure 5A, Supp. Figure 6, Supp. Figure 7 and Supp. Figure 8). To quantify the association between the two metrics (DPC and sequence similarity), we computed the Pearson correlation coefficient between the pairwise DPC matrices and the pairwise sequence similarity matrices (Figure 5B, Supp. Figure 9A). We repeated this analysis for 100 randomly sampled sets of 100 sequences each (within the same IGHV gene family). Samples were taken from all isotypes of the native dataset to account for variation in associations among batches. We restricted the single set (batch) size to 100 antibodies to ensure correlation matrix regularization (*63*). We examined the resulting Pearson correlation values alongside the average sequence similarity score (100 values for each metric for the 100 sampled sets per isotype – Figure 5B). We found that Pearson correlation coefficients (between DPC and average sequence similarity) tended to be higher on the sequence level (0.2–0.7) than on the structure level (0.1–0.4) across all antibody isotypes (Figure 5B). For instance, the mean Pearson correlation coefficient for the human IgD dataset was 0.5 on the sequence level and 0.2 on the structure level (Figure 5B). This finding suggested that similar sequences exhibit higher sequence-based developability similarity compared to structure-based developability similarity. However, higher sequence similarity was not always accompanied by higher Pearson correlation values of DP profiles. For example, although the average sequence similarity of the murine IgL dataset was as high as 0.9, the mean value of Pearson correlation coefficient was only 0.5 (Figure 5B). We reported a similar Pearson correlation average (0.5) for the human IgE dataset, even though its mean sequence similarity was less than the IgL murine dataset (0.7, Figure 5B).

**Figure 5.**
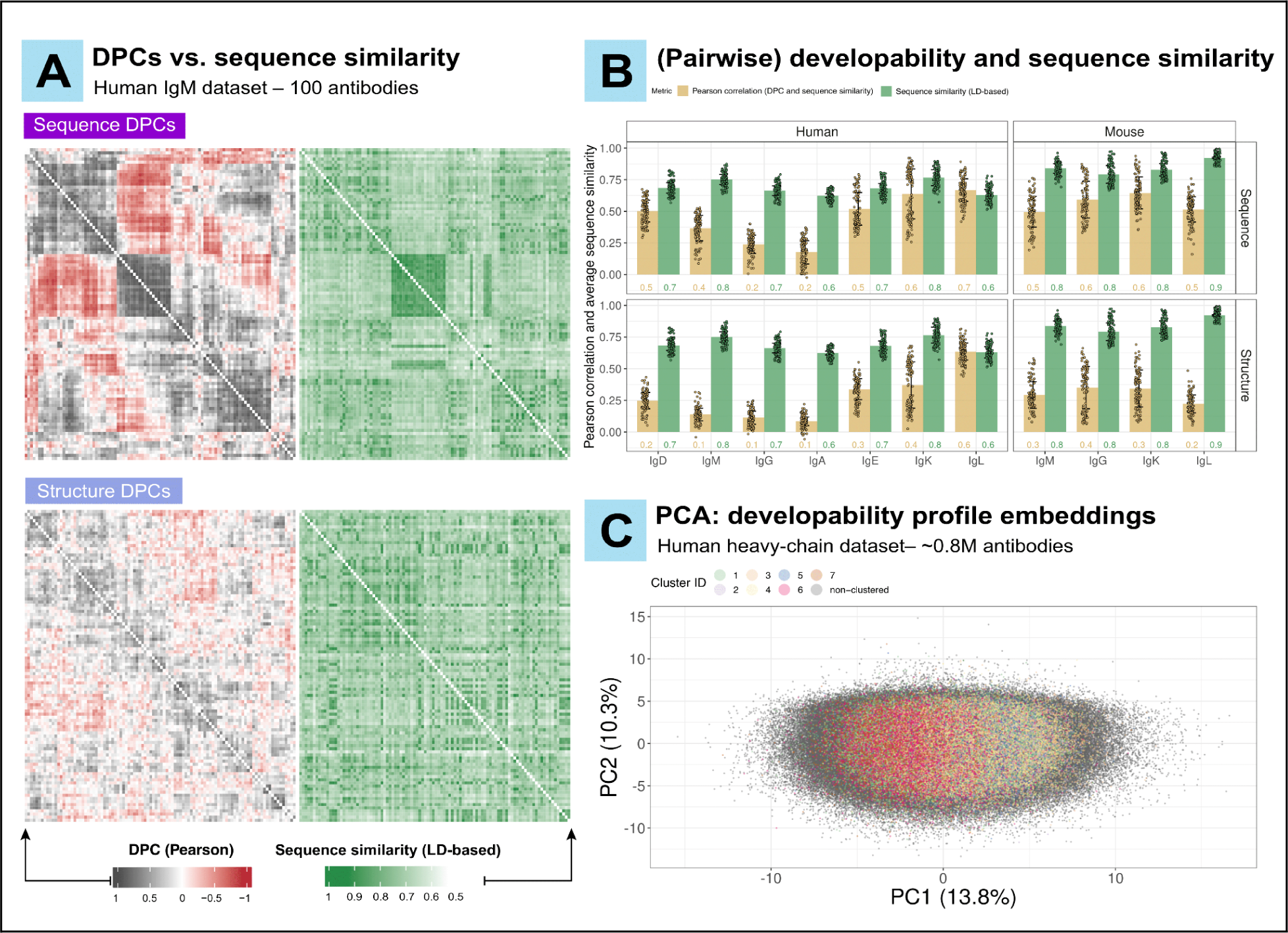
Developability profile similarity is not necessarily associated with sequence similarity. (A) Pairwise <u>d</u>evelopability <u>p</u>rofile Pearson <u>c</u>orrelation (**DPC** - left panels) alongside the pairwise Levenshtein distance (LD) based-sequence similarity score (right panels – see Methods) for a random sample of 100 antibodies from the human IgM dataset (100x100 matrices) that share the same IGHV gene family (IGHV1) annotation (shown both for sequence and structure DPLs). Each row and each column represent a single antibody sequence. Rows and columns in the left panels were hierarchically clustered. In the right panels (sequence similarity), rows and columns were ordered in the same order as the corresponding left panel (DPC) for ease of comparison. The distribution of DPC and sequence similarity is shown in Supp. Figure 9A. **(B)** Pearson correlation between DPC and sequence similarity matrices for 100 sets of randomly sampled non-overlapping 100 antibody sequences (within the same IGHV gene family per batch) from all isotypes of the native dataset. Pearson correlation coefficient values (shown in beige) are presented alongside the corresponding mean sequence similarity values (shown in green) for the same 100 sets. The height of the bars and the numerical values on the figure reflect the mean of the corresponding metric (mean Pearson correlation and the mean sequence similarity). The error bars represent the standard deviation. **(C)** Principal component analysis (PCA) of the developability profiles of the native human heavy-chain dataset (∼0.8M antibodies). The developability profiles (DPLs) were utilized as embeddings for this analysis (see Methods). Antibody clusters (1–7) were created for the groups of antibodies that are at least 75% similar in sequence (as determined by USEARCH) and contain at least 10K antibodies. Antibodies that did not satisfy the clustering conditions were labeled as “non-clustered” (727861 sequences) and sent to the back layer of the figure. For antibody counts per cluster, please refer to Supp. Figure 9B. **Supplementary Figures:** Supp. Figure 6, Supp. Figure 7, Supp. Figure 8 and Supp. Figure 9.

Next, we sought to investigate the relationship between antibody developability and sequence similarity using a geometric approach to test the finding that developability profile similarity and sequence similarity are not necessarily associated (Figure 5A,B). We leveraged the fact that antibody developability profiles are numerical vectors in the developability space (*R^N^*), where *N* represents the number of DPs that compose a single developability profile (see Methods). This space (*R^N^*) offers a natural notion of antibody developability diversity. However, it is challenging to inspect due to its high dimensionality. Thus, we applied a dimensionality reduction technique (principal component analysis: PCA) on the developability profile space of the human heavy-chain antibody dataset (Figure 5C) as it is the largest data subset that belongs to a single species and chain type (∼0.8M antibodies, Supp. Figure 1A). Within this subset, we identified seven sequence similarity groups (1–7) where each group encapsulates at least 10K antibodies and group members exhibit at least 75% pairwise sequence similarity (Supp. Figure 9B – see Methods). We found that antibody members that belong to the same sequence similarity group did not occupy a restricted space on the PCA projection plane, suggesting that antibody developability and sequence similarity are not correlated (Figure 5C). To quantify this observation, we studied the correlation between the pairwise Euclidean distances in the developability space (*R^N^*) and the pairwise (normalized) Levenshtein distances for 5000 antibody sequences from the human IgM dataset that share the same IGHV gene family annotation (IGHV3) and belong to the same sequence similarity group (group 1 – Supp. Figure 9B,C). We found that the two distance measures showed only minimal correlation (Pearson correlation coefficient = 0.18, Supp. Figure 9C), which indicated the independence of developability profile and CDRH3 sequence similarity among antibodies of the native dataset.

In conclusion, our analysis demonstrated that antibody developability and sequence similarity were largely independent, suggesting that improving the developability profile for a certain therapeutic mAb candidate with a desirable target binding profile may be possible by introducing small changes in its amino acid sequence.

### DP predictability implies interdependence and antibody design space restriction

Building on the prior finding that no significant association exists between antibody sequence similarity and developability similarity, we inquired whether missing values of given DPs can be predicted based on the knowledge of the values of other DPs. This is to investigate to what extent the developability space is amenable to orthogonal DP design (Figure 6A). In this context, high predictability of a given DP would indicate a restriction of the antibody design space, while low predictability could signify a more plastic space with a higher degree of freedom for the values of this DP. Answering the above question would also provide insights into which DPs can be better predicted (with the provided knowledge of the remaining DPs), which may accelerate antibody developability screening. Also, we evaluated the predictability of missing DP values depending on the sole knowledge of the amino acid sequences of antibodies (through their protein language model representations, Figure 6A). This aims to investigate the feasibility of DP predictability in the absence of other DP data.

**Figure 6.**
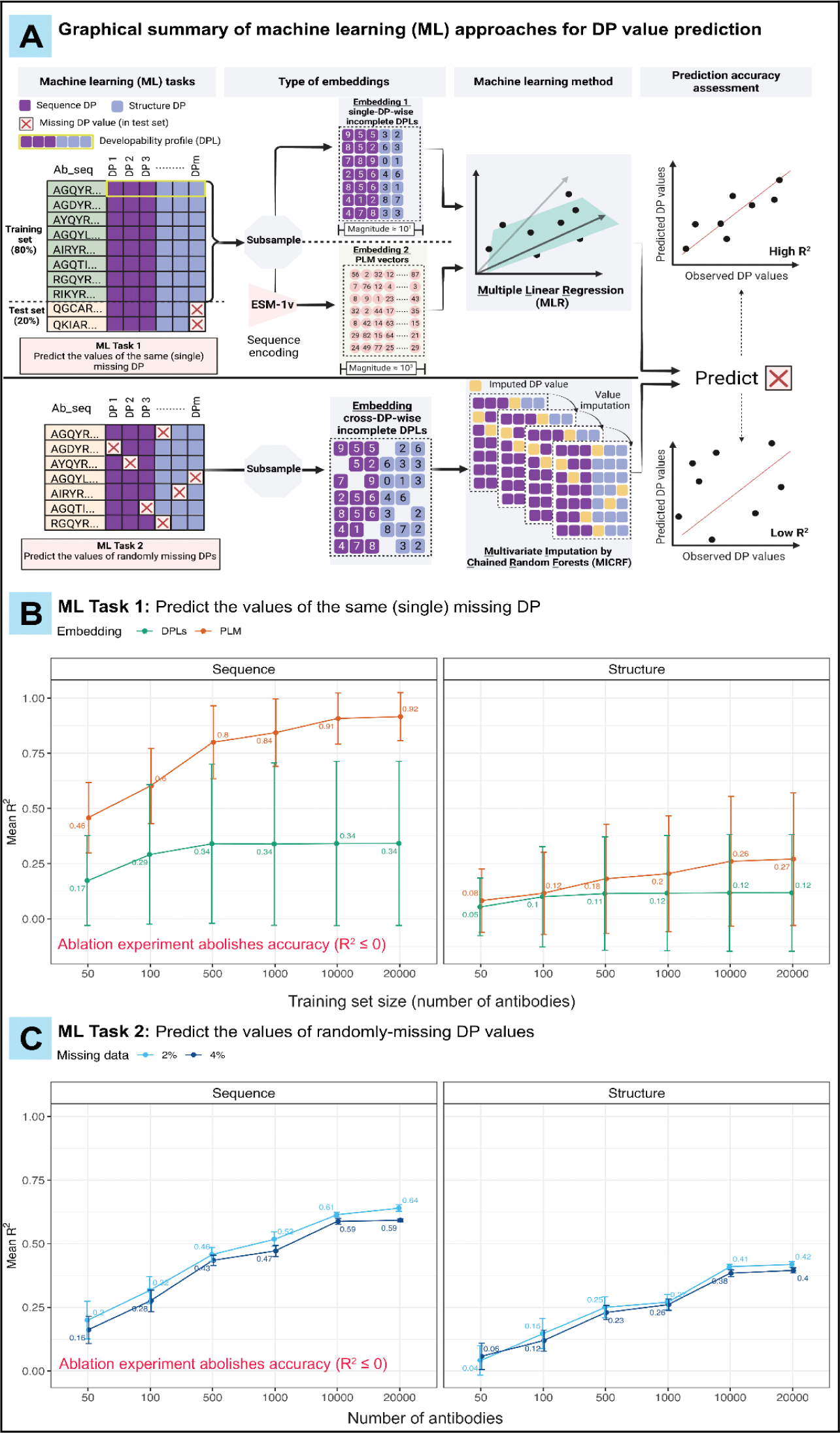
**Sequence-based developability parameters are more predictable than structure-based parameters**. Graphical representation of machine learning (ML) approaches used to assess the predictability of DPs. We investigated two scenarios where the missing (deleted) DP values were either all from one (single) DP (ML Task 1) or were randomly missing from several DPs (ML Task 2). For <u>ML Task 1</u>, we compared the predictive accuracy of two different embeddings; single-DP-wise incomplete developability profiles (DPLs) (embedding 1; order of magnitude 10^1^) and PLM vectors (embedding 2; order of magnitude 10^3^). We used these embeddings to train multiple linear regression (MLR) models (separately) to predict the missing DP values in the test set. To enable the comparison between these two embeddings, we used identical training subsamples (in regards to size and antibody identity, see Methods). For <u>ML Task 2</u>, we used cross-DP-wise incomplete developability profiles as input for the multivariate imputation by chained random forests (MICRF) algorithm to predict missing DP values. For both ML tasks, we estimated the prediction accuracy by computing the coefficient of determination (R^2^) using observed and predicted DP values (*78*). **(B)** Comparison of the predictive accuracy of incomplete developability profiles (single-DP-wise incomplete DPLs) and PLM vectors as embeddings for MLR models to predict the values of missing DPs in the test set (ML Task 1). The x-axis reflects the number of antibody sequences (sample size) used for the embedding. For each sample size, we repeated the prediction of missing DPs 20 times (20 independent subsamples). The y-axis represents the mean R^2^ for sequence DPs (left facet) and structure DPs (right facet). Error bars represent the standard deviation of R^2^. Missing DPs tested in this analysis belonged to the MWDS exclusively, as determined at a Pearson correlation coefficient threshold of 0.6, for the human IgG dataset, summing to 13 sequence DPs and 28 structure DPs (after removing a single element from each doublet and immunogenicity DPs, Supp. Table 3). **(C)** Evaluating the predictability of randomly missing DP values using the MICRF algorithm where cross-DP-wise incomplete developability profiles are used as embeddings. The x-axis reflects the number of antibodies (sample size) used for the embedding. For each sample size, we repeated the prediction of missing DPs 20 times (20 independent subsamples). The y-axis represents the mean R^2^ for sequence DPs (left facet) and structure DPs (right facet) when the proportion of the missing data is either 2% (light blue line) or 4% (dark blue line). Error bars represent the standard deviation of R^2^. Missing DPs tested in this analysis belonged to the MWDS, analogously to (B). **Supplementary Figures:** Supp. Figure 14 (DP predictability comparison by redundancy).

Importantly, the primary objective of this analysis is to evaluate how various factors (type of ML input, type of predicted DP, the redundancy state of the predicted DPs) affect the predictability of DPs rather than achieving absolute predictive precision given that we did not perform a comprehensive benchmarking of encoding or ML approaches (*64*, *65*). Ultimately, when comprehensive benchmarking and fine-tuning of multitask ML models is performed on developability data, it might be possible to depend on these models for DP value predictability, rather than relying on several *in silico* tools to achieve the same task.

We initially used MWDS DPs to eliminate collinear dependence that may increase ML prediction accuracy in a non-interesting or trivial manner. To this end, we investigated two scenarios where the missing (deleted) DP values were either all from a single DP (ML Task 1), or randomly missing from several DPs (ML Task 2) (Figure 6A). In practical terms, ML Task 1 replicates a use case where the values of a single DP – that might be challenging to compute or measure – are missing from all antibodies in the dataset. Meanwhile, as “spotted” (i.e., missing-at-random) data is a real-world problem in biomedical research (*66–68*), ML Task 2 replicates the use case of obtaining a developability dataset where the values of several DPs are sporadically missing from the antibodies (Figure 6A).

For ML Task 1, we compared the predictive accuracy of two types of (input) embeddings to predict the missing DP values via multiple linear regression (MLR) models after defining training and test (sub)sets from the native human V_H_ antibody dataset (Figure 6A, see Methods). (i) The first embedding is the single-DP-wise incomplete developability profiles (DPLs) which depend on the knowledge of all other DP values included in the training set (low-dimensional embedding – order of magnitude ≈ 10^1^ – Figure 6A). (ii) The second embedding is an amino acid sequence encoding output produced by the protein language model (PLM) ESM-1v, which generates large semantically-rich digital representations of antibodies (high-dimensional embedding – order of magnitude ≈ 10^3^) (*69–71*). Unlike DPL-based embedding, PLM-based embedding is entirely unaware of antibody DP values (Figure 6A) and we aim to test whether these biochemical properties are implicitly contained in it.

We found that the predictive accuracy of MLR models increased with increasing training set size for both embeddings. However, DPL-based models reached their saturation point earlier than PLM-based ones for both sequence and structure DPs (1000 antibodies for DPLs, 20000 for PLM – Figure 6B). This is also (partly) attributable to the fact that higher dimensional inputs (PLM embeddings) correspond to additional degrees of freedom when training the model. Overall, both embeddings achieved higher prediction accuracy for sequence DPs compared to structure DPs at their saturation points (Figure 6B). However, the disparity in prediction accuracy between the two embeddings was more pronounced for sequence DPs, with PLM-based embeddings achieving a mean prediction accuracy of 0.92 compared to 0.34 for DPL-based ones (Figure 6B). Thus, the high predictability enabled by PLM-based embedding on sequence level highlighted its capacity to capture the biophysical properties of antibodies based on amino acid sequence (*72*).

When conducting an analogous analysis including non-MWDS (redundant) DPs, we found notable improvement in DPL-based MLR models to predict missing DP values (mean R^2^ of 0.88 for sequence DPs and 0.36 for structure DPs) (Supp. Figure 14A). This is due to the inherent collinearity among non-MWDs DPs (Figure 2B) that simplifies DPL-based prediction, resulting in higher prediction accuracy (*73*). In contrast, PLM-based embeddings showed far lesser to no distinct improvements when predicting non-MWDS DPs (mean R^2^ of 0.96 for sequence DPs and 0.38 for structure DPs) (Supp. Figure 14A).

For ML Task 2, we implemented the multivariate imputation by chained random forests (MICRF) algorithm (*67*, *74*) to evaluate its prediction accuracy to recover randomly missing (deleted) DP values from the developability dataset (Figure 6A – see Methods). We investigated two cases where either 2% or 4% of DP values are missing from either sequence or structure MWDS parameters (Figure 6C), as implementing the MICRF algorithm with a greater fraction of missing data could compromise the accuracy and reliability of the imputed (predicted) DP values (*74*). Overall, and similarly to the observations reported from ML Task 1 (Figure 6B), structure DPs were more challenging to predict, and a larger number of data inputs (antibodies) aided the achievement of higher prediction accuracies (Figure 6C). However, we noticed (only) subtle differences in the prediction accuracy (mean R^2^) when comparing the algorithm capacity to restore 2% or 4% of missing DP values (Figure 6C), outlining its robustness within the advised data loss limits for its application (*75*, *76*). For example, we reported mean R^2^ of 0.64 (sequence DPs) and 0.42 (structure DPs) with 2% data loss compared to 0.59 (sequence DPs) and 0.4 (structure DPs) with 4% data loss, at a saturation point of 20000 antibodies (Figure 6C).

Similarly to ML Task 1, non-MWDS DPs were shown to be easier to predict (Supp. Figure 14B). However, the disparity in prediction accuracy between the two classes of DPs (MWDS and non-MWDS) was less pronounced in comparison to DPL-based predictions in ML Task 1. For instance, at 2% data loss, the improvement in prediction accuracy of sequence DPs was (only) ∼0.2 higher (0.85 for non-MWDS, 0.64 for MWDS – Supp. Figure 14B) compared to ∼0.6 in DPL-based predictions in ML Task 1 (0.88 for non-MWDS, 0.34 for MWDS Supp. Figure 14A).

Of note, we performed ablation studies (*77*) on both ML tasks, by randomly permuting the values of DPs at the columns in the input data (feature shuffling), and confirmed that the prediction accuracy was diminished (R^2^ ≤ 0, Figure 6B,C, see Methods).

In summary, for the ML methods and embeddings applied in this analysis, we found that the structural developability space is less restricted. Additional analyses suggested that variations in structure prediction alone are insufficient to explain this (Supplementary File). Additionally, this analysis highlighted the potential of PLMs to reflect the sequence-based aspects of antibody developability, if provided with sufficient training data. We want to emphasize that our main goal was not obtaining perfect prediction accuracy of DPs but rather measuring the amount of readily available biological information contained in the two representations examined, i.e. DPL- and PLM-based representations. Finally, we also addressed the presence of potential gaps in developability information in practical settings, by showing how two different ML algorithms can be utilized in two different scenarios of missing data.

### Patent-submitted, humanized mouse (Kymouse) and therapeutic monoclonal antibodies represent a subset of the natural developability atlas

Previously, clinically approved therapeutic antibodies were used to build classifiers for optimal developability (*7*, *33*). However, the higher abundance of native antibody data incentivizes the investigation of how human-engineered antibodies relate to their native counterparts. Thus, we investigated to what extent we can detect differences between native and human-engineered antibodies (therapeutic mAbs, Kymouse and PAD). To this end, we examined the proportional abundance of liability sequence motifs and the representation of developable germlines across the native and human-engineered datasets (Supp. Figure 13). Liability motifs are short amino acid sequences, which may negatively impact various aspects of antibody developability when present in their CDRs (*79*). Developable germlines represent a group of human immunoglobulin genes (V_H_ and V_L_), which have been suggested to harbor favorable biophysical properties (*80*, *81*).

The native dataset was comparable to the human-engineered ones in regards to the proportion of antibodies that exhibit liability motifs, and no consistent increased or decreased representation (trend) of these motifs was reported (Supp. Figure 13A). For instance, while the native human V_L_ dataset exhibited a higher abundance of the asparagine deamidation motif “NS” (17.8%) compared to PAD and mAbs (15.7%, 16.6%), it exhibited a lower abundance of the proteolysis motif “DP” (0.5%, 2%, 3.8% - Supp. Figure 13A). Similarly, the native dataset harbored a comparable representation of the developable germline genes with the exception of IGHV1 members (5.2% in native, 3.1% in Kymouse, 16.6% in PAD, and 21.6% in mAbs) and IGK1 members (13.3% in native, 20.8 in PAD, and 25.9 in mAbs – Supp. Figure 13B). These findings suggest a notable resemblance between the native and human-engineered antibody datasets with regard to potential developability-related liability sequence motifs and germline annotations.

Thus, we asked how the native antibody dataset relate to the human-engineered ones in terms of DP values. To this end, we first conducted a distance-based clustering (similar to Figure 3B) starting from the pairwise DP correlation matrices of the native and human-engineered antibody datasets (Figure 7A). On the sequence level, we found that the general species-specific (human and mouse) and chain-specific (light and heavy) clustering, as previously reported in our investigation of native dataset (Figure 3B; left panel), remained consistent when integrating native with human-engineered antibody datasets (Figure 7A; top panel). Specifically, the human antibodies (light and heavy chains) from the PAD and mAbs datasets localized within the native human clusters of the same chain type (correlation distance ≤0.1 – Figure 7A; top panel). Similarly, the light-chain human-engineered murine antibodies (PAD and mAbs) clustered with the IgK native mouse dataset (correlation distance ≈ 0.1 – Figure 7A; top panel). Kymouse antibodies were the most distant subset, which may be explained by their unique intermediate diversity between mice and humans (*82*).

**Figure 7.**
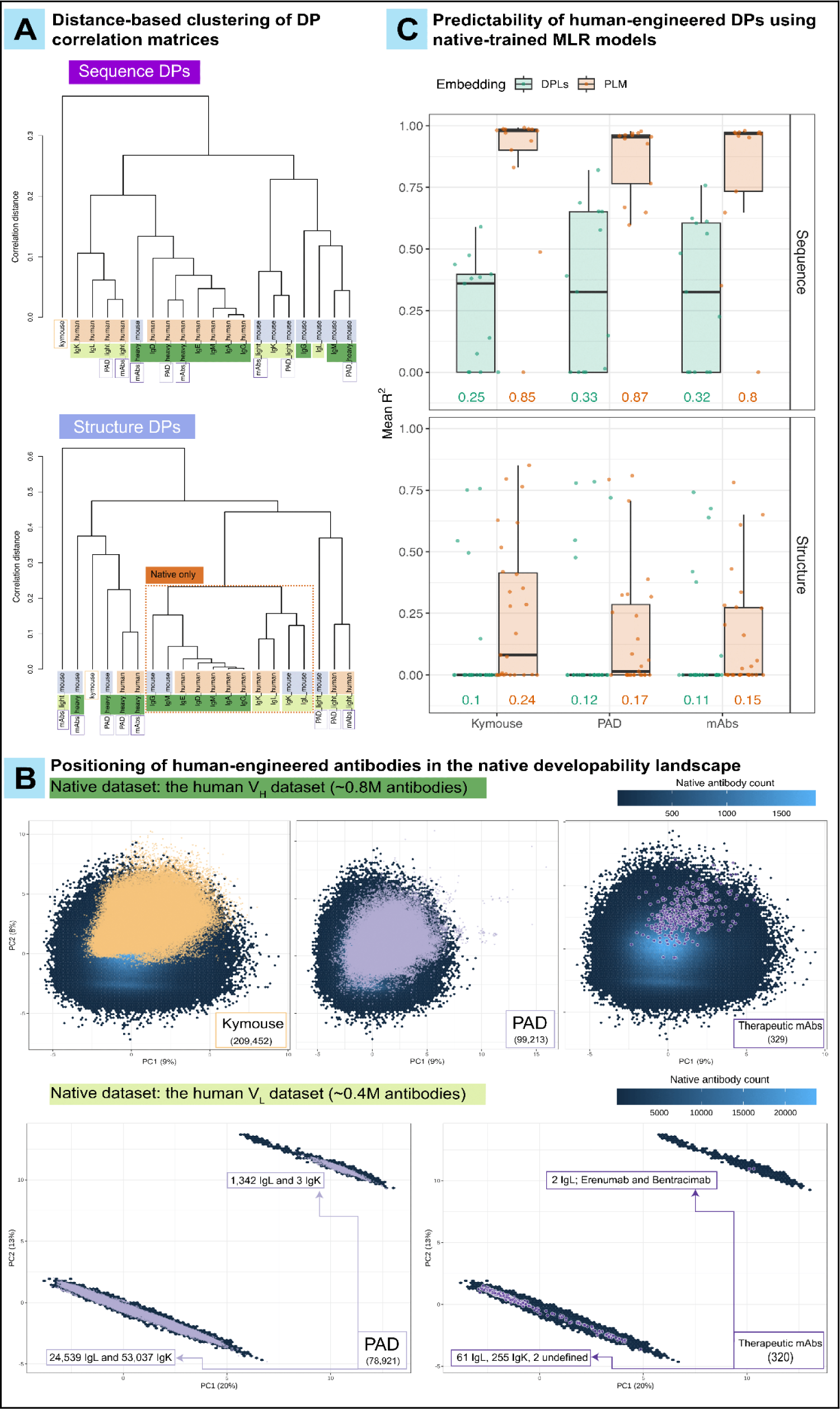
Human-engineered antibodies are contained in the developability landscape of natural antibodies. Distance-based hierarchical clustering of isotype-specific pairwise DP correlation matrices (sequence and structure levels - similar to the analysis shown in Figure 3A). The height of the dendrograms (shown to the left of the figure) represents the correlation distance among the dendrogram tips. The dashed square in the right (structure-based) panel highlights the native-only dataset. **(B)** Top three panels: The positioning of the human-aligned human-engineered V_H_ antibodies (Kymouse; 209,452 , PAD; 99,213 and therapeutic mAbs; 329) in the developability profile space of the native human V_H_ dataset (854,418 antibodies) based on a principal component analysis (PCA, see Methods). Bottom two panels: The positioning of the human-aligned human-engineered V_L_ antibodies (PAD; 78,921 and therapeutic mAbs; 320) in the developability space of the native human V_L_ dataset (385,633 antibodies). The hexagonal bins (shown in the back layer) represent the count of native antibodies (scale shown on the top right of the panels), and the human-engineered antibodies are represented as data points. **(C)** Evaluating the predictability of sequence (left panel) and structure DPs of the human-aligned human-engineered V_H_ antibodies (Kymouse; 209,452 , PAD; 99,213 and therapeutic mAbs; 329), using multiple linear regression (MLR) models trained on native human V_H_ antibodies. As explained in Figure 6A (ML Task 1), the predictive accuracy of two types of embeddings was tested, including single-DP-wise incomplete developability profiles (DPLs) and ESM-1v protein language model encoding vectors (PLM). MLR models were trained using 1000 antibodies for DPL-based predictions and 20000 antibodies for PLM-based predictions (respective saturation points). Missing DPs tested in this analysis belonged to the MWDS exclusively as determined at a Pearson correlation coefficient threshold of 0.6, for the native human IgG dataset, summing to 13 sequence DPs and 28 structure DPs (Supp. Table 3). The y-axis represents the mean coefficient of determination (R^2^) across 20 repetitions. Numerical values shown represent the mean R^2^ across (sequence or structure) DPs. **Supplementary Figures:** Supp. Figure 10, Supp. Figure 11, Supp. Figure 13, Supp. Figure 15

On the structure level, we found that the original clustering pattern among the native antibody isotypes is preserved from the analysis conducted on the native-only dataset (Figure 3B, Figure 7A; “Native only” zone). Human-engineered subsets clustered apart from the “Native only” zone, maintaining either species-specific or chain-specific clustering. Although suggestive, the results of this analysis should be interpreted with caution due to the imbalance of dataset sizes (number of antibodies) (Supp. Figure 1A,B). Indeed, a correlation matrix stabilization study suggested that a dataset size of at least 50K antibodies is required to stabilize association values among DPs (Supp. Figure 15A,C) and the distribution of their pairwise Pearson correlation values (Supp. Figure 15A,B,D). This threshold (50K antibodies) is higher than the count of antibodies included in the murine subsets of the PAD dataset (≈24K for heavy chains, ≈21K for light chains) and all the species-specific and chain-specific subsets within the mAbs dataset (329 for human heavy chains, 320 for human light chains, 62 for murine heavy chains, 71 for murine light chains) (Supp. Figure 1B).

Next, we leveraged the principal component analysis (PCA) that we conducted on the developability profiles of native antibodies (Figure 5C – see Methods) to examine how human-engineered antibodies relate to the native ones in the developability spaces (V_H_; Figure 7B: top panels, V_L_; Figure 7B: bottom panels). In addition to comparing native and human-engineered antibodies in the *developability profile* space (dimensionality: 10^1^), we also used *sequence* embeddings provided by the ESM-1v (*70*) protein language model (PLM, dimensionality: 10^3^) (Supp. Figure 10B, Supp. Figure 11B). PLM embedding is an alternative to biologically motivated features (such as DPL), learned without supervision from a large pool of proteins (*83*). Protein language modeling enables capturing longer-distance relationships within protein sequences (*84–86*). We focused our analysis on human antibodies as they represent the largest species-specific subset among our datasets (≈0.8M for native V_H_ antibodies, ≈0.4M for native V_L_ antibodies, 99213 for PAD V_H_ antibodies, 78921 for PAD V_L_ antibodies, 329 for V_H_ mAbs and 320 V_L_ mAbs – Figure 7B, Supp. Figure 1A,B).

Overall, we found that human-engineered antibodies (both V_H_ and V_L_) are majorly contained within the developability and PLM landscapes of the native antibodies (Figure 7B, Supp. Figure 10B, Supp. Figure 11B), suggesting that – for the DPs included in our analysis (see Methods) – the developability and sequence landscapes of human-engineered antibodies are mere subspaces of the natural ones (in terms of the two main axes of variation studied).

From the native repertoire perspective, V_H_ antibodies coalesced into a single cluster in the V_H_ developability space, and the positioning of the antibodies in this space was independent of both their isotype and IGHV gene family annotation (Supp. Figure 10A). In contrast to V_H_ antibodies, native V_L_ antibodies clustered in two distinct clusters in the V_L_ developability space where the majority of IgK antibodies (99.999%) and IgL antibodies (95.6%) occupied the bottom cluster, and a small proportion of IgL antibodies (4.4%) predominantly occupied the top cluster (Supp. Figure 11A). This finding suggests that, except for a minor subset of native IgL antibodies, V_L_ sequences (both IgK and IgL) are homogeneous with respect to their developability. This aligns with a recent report where native IgL antibodies exhibited comparable structural developability characteristics to those of native IgK antibodies (*33*). Among the human-engineered antibodies, only two (out of 320 V_L_) therapeutic mAbs (Erenumab and Bentracimab) of the isotype IgL, and only 1,345 (1,342 IgL and 3 IgK, out of 78,291 V_L_) patent-submitted antibodies were found to be contained in the top cluster (Figure 7B), displaying a similar distribution trend between the two clusters as the native V_L_ dataset. It is worth noting that Erenumab has been previously reported by other studies to exhibit developability risk factors (*7*, *10*), which may explain its localization in the top cluster of the V_L_ developability space (Figure 7B). Bentracimab was not included in these studies as it is not yet clinically approved (Phase III of clinical development as of September 2023) (*87*).

As the human-engineered antibodies were shown to harbor comparable developability and sequence properties to those of the native ones (Figure 7B, Supp. Figure 10B, Supp. Figure 11B), we investigated the generalisability of the native-trained multiple linear regression (MLR) models to predict the values of single missing DPs of the human-engineered datasets (Figure 7C). Specifically, we implemented the MLR models from ML Task 1 (Figure 6A,B) at the training sample size, which achieved the highest prediction accuracy before plateauing (1000 antibodies for DPL-based predictions, 20K antibodies for PLM-based predictions) to predict the values of MWDS DPs (sequence and structure) on the human-engineered antibody datasets (Figure 7C).

We found that the predictability of DP values for the human-engineered antibodies was similar to that of the native antibodies (Figure 6B, Figure 7C). For instance, the mean prediction accuracy (mean R^2^) of sequence DPs was between 0.25–0.33 for DPL-based predictions, and between 0.80–0.87 for PLM-based predictions among human-engineered datasets (Figure 7C) in comparison to 0.34 and 0.92 (DPL and PLM respectively) for native antibodies (Figure 6B). Similarly, when considering human-engineered datasets, the mean prediction accuracy of structure DPs ranged from 0.10 to 0.12 for DPL-based predictions and from 0.15 to 0.24 for PLM-based predictions (Figure 7C). In comparison, the native antibodies exhibited prediction accuracies of 0.12 and 0.27 (respectively – Figure 7C).

Collectively, our results suggest that the knowledge learned from the native antibodies in regards to their developability and sequence properties is in part generalizable to the human-engineered antibody datasets. Nevertheless, it is worth reiterating that the predictability analysis performed on the native and human-engineered datasets aims to suggest an experimental design and investigate generalizability rather than improving predictability.

## Discussion

### Systems immunology approach to profiling the plasticity of the developability space ensures real-world relevance

Previous studies have shown that a number of experimental DPs may be computationally inferred (*7*, *13*, *34*, *52*, *88*), which supports the real-world relevance of computer-based developability screening of large antibody sequence libraries. However, discrepancies between computational and experimental DP profiling remain (*18*, *31*). Furthermore, previous studies using computational profiling varied in the number of DPs used, ranging from less than 10 to more than 500 (*10*, *51*, *53*). In this study, we used 86 sequence and structure-based DPs, many of which were pairwise lowly or uncorrelated to one another (Figure 2A,B, Supp. Figure 2, Supp. Figure 3), thus capturing at least a subspace of the multidimensional developability space. So far, the relevant dimensionality of the antibody developability space remains unclear. That said, our analysis can be replicated with any other set of computationally or experimentally determined DPs. While the majority of previous studies focus on small antibody datasets and mostly on the comparison between experimental and computational DPs, this study explores the plasticity of the *in silico* DP space on large-scale antibody datasets. While our findings may partly depend on the DPs studied, conclusions based on DP profiling agreed with PLM-based profiling (Figure 7B, Supp. Figure 10B, Supp. Figure 11B). In other words, our study should be understood as forecasting the types of analyses possible once large-scale experimental or highly validated computational DPs exist, which would allow for routine repertoire-scale DP profiling and prediction. In summary, the real-world relevance of our systems approach is grounded in (i) the profiling of a large number of pairwise-independent DP parameters, (ii) sequence and rigid and dynamic structure-based DP analysis (Supplementary File), (iii) diverse experimental datasets ranging from native to human-engineered, and (iv) parameter-independent computational and machine learning analysis. In the future, it would be of interest to add display libraries for comparison (*89–91*) as well as a larger number of paired datasets with broad and deep isotype information (*92*, *93*).

### Developability parameter redundancy may be reduced through analysis of intercorrelations

We found that structure-based DPs show higher independence (lower pairwise correlation) than sequence-based DPs (Figure 2). These findings emphasize the significance of structural consideration for therapeutic antibody design (*3*, *8*, *25*, *94*, *95*). Due to the scarcity of antibody structural information, structure-based developability estimation has previously presented a substantial challenge in developability screening (*3*, *20*). However, *in silico* high-throughput antibody structure prediction tools have been evolving in speed and accuracy with machine learning algorithms, permitting the screening of structural parameters in large datasets. (*9*, *37*, *55*, *96*).

The scale of this work’s analysis facilitated the discovery of parameter redundancies that could potentially accelerate future antibody developability screening processes. For instance, while Chen and colleagues included both the molar extinction coefficient and the extinction coefficient of the variable region sequence (AbChain_molextcoef, AbChain_percentextcoef) and the cysteine bridges (AbChain_cysbridges_molextcoef, AbChain_cysbridges_percenextcoef) as important developability predictors (*53*), we found that one of these coefficients could likely be sufficient to replace the other with near-perfect pairwise correlations (>0.9) for all chain types (V_H_, V_L_) included in our analysis (Figure 2B, Supp. Figure 2, Supp. Figure 3). Similarly, although Ahmed and colleagues highlighted the importance of structure-based isoelectric point (pI) as an essential developability parameter on a limited-size therapeutic antibody dataset (77 clinical-stage antibodies) (*10*), our analysis suggested that sequence-based pI (AbChain_pI) could potentially replace structure-based pI with high pairwise correlation with structure-based pI measure on folded and unfolded variable region structure for native human IgG antibodies (Figure 2B) and across all other isotypes and chains in the native human and mouse datasets (Pearson correlation 0.9–1, Supp. Figure 2, Supp. Figure 3). We further highlighted the importance of large-scale antibody developability data to stabilize the associations among DPs (Supp. Figure 15), and, thus, to reveal such redundancies. Additionally, most developability studies emphasize the relevance of antibody charge in the physiological pH (7.4) (*7*). However, mAbs are usually exposed to a wider range of solution pH (4.8-9) during production and formulation (*10*, *97*). Also, antibody variable region charge in acidic pH has proven to be a critical factor in IgG mAb pharmacokinetics (*98–100*) and product formulation (*29*). Therefore, we included the sequence-based variable region charge measures in 14 pH points (1–14) in our analysis. We found that the charge measures of native IgG sequences formed three correlation clusters with intermediate (0.4–0.7) pairwise correlations, emphasizing the importance of antibody charge considerations on a wider spectrum of pH values (Figure 2B). A similar three-cluster pattern was also identified across all other isotypes of the native dataset (Supp. Figure 2, Supp. Figure 3). Other pairs of potential parameter redundancies are the sequence content of polar and non-polar amino acids (AbChain_polar_content Vs AbChain_nonpolar_content), the sequence content of aliphatic amino acids and the sequence aliphatic index (AbChain_aliphatic_content Vs AbChain_aliphatic_index), the average atomic interaction distance of the antibody structure and the number of Van der Waals clashes (AbStruc_mean_interaction_distance Vs AbStruc_vdw_clashes) and the sequence length and the sequence molecular weight (AbChain_mw Vs AbChain_length) (Figure 2A). Redundancy-based parameter reduction, analogous to feature selection for ML models, would accelerate future antibody developability investigations by screening for a more comprehensive and sufficiently representative set of parameters.

### Relevance of species- and chain-specific developability spaces for antibody discovery efforts

Our analyses revealed chain-specific developability signatures, in relation to DP values and pairwise associations (Figure 3B,C), emphasizing possible differences in developability design considerations for therapeutic antibody development. Within each chain type, we found that murine and human antibodies occupy distinguishable developability spaces (Figure 3C), which highlights the importance of transgenic mice for antibody screening and the challenges of antibody humanization efforts (*101–104*). Interestingly, our findings suggest that the Kymouse (humanized mice) dataset under investigation undersampled the human dataset, both with respect to developability (Figure 7B) and sequence spaces (Supp. Figure 10B), even though lower-level features such as VDJ gene usage and CDR3 length were previously found to overlap (*82*).

In addition to chain type and species, we investigated whether antibody isotype is associated with similarities in developability parameters. We found that human heavy chain (V_H_) isotypes harbor high similarities in regard to their pairwise DP associations (Figure 3B) and redundancies (Supp. Figure 4B). (Figure 3A). We found that they aggregated homogeneously in the developability space regardless of their isotype (Supp. Figure 10A). Thus, although all currently approved therapeutic mAbs belong to the IgG isotype (*105*), our findings provide the incentive to explore the available native antibody Fv sequence space beyond the isotype annotation for novel mAb discovery. In regards to V_L_ isotypes, human IgK and the majority of IgL antibodies clustered together in the developability space (Supp. Figure 11A), questioning the previous association of IgL antibodies with poor developability, in line with recent findings by Raybould and colleagues (*33*), and providing an incentive to re-include IgL sequences in future antibody discovery libraries. Nevertheless, we are aware that our findings involve only the Fv regions of antibody sequences as current antibody-specific structure prediction models do not take into account the Fc region and do not account for the impact of the Fc region on developability (*100*, *106*).

### Developability similarity is only loosely related to antibody sequence similarity

Our findings suggest that antibodies that are highly similar in sequence can possess dissimilar developability profiles (Figure 5), which is in line with previous findings (*107*). This may suggest that there are degrees of freedom available for therapeutic antibody candidate engineering to optimize developability with minimal changes in antibody sequence.

For instance, if an antibody candidate exhibited optimal antigen binding properties with a suboptimal developability profile, there could be minor sequence changes that result in a substantial improvement (or deterioration) of this profile without major changes to its antigen binding properties. In this context, Petersen and colleagues showed that the native human antibody repertoire can aid the identification of advantageous (or disadvantageous) “universal” (native) framework mutations that could facilitate therapeutic mAb development (*50*). Among these mutations, S54A was found in four FDA-approved mAbs (Pertuzumab, Atezolizumab, Omalizumab, and Trastuzumab), and was associated with increased structural stability of these antibodies (*50*). Furthermore, small changes in antibody sequence with large effects on function have been observed for both developability and antibody binding properties (*3*, *108*), underscoring the importance of simultaneous optimization of multiple design parameters. Since optimization of a specific property often comes with undesirable trade-offs for other properties (*30*, *58*), studying the effect of substitutions can help with designing antibodies that exhibit improved developability, with comparable efficacy.

To directly probe the relationship of sequence to developability, we performed a single amino acid substitution analysis. Although changing one amino acid at a time only explores a small fraction of the input factor space (*59*), it allows for the definitive attribution of the change in output to a specific amino acid substitution. Briefly, we found that some parameters were especially insensitive to the average mutation. More generally, including all possible variants with increasing numbers of substituted amino acids would geometrically inflate the space of possible substitutions. Because exhaustively mapping the DPs of such large sequence spaces is (currently) computationally infeasible, to explore more than single amino acid substitutions, one must find ways to efficiently and uniformly sample the input space, possibly through latin hypercube sampling or low-discrepancy sequences (*109*, *110*). Furthermore, to quantify the sensitivity of a parameter, we measured the dispersion of parameter values of all possible single amino acid substituted variants of an antibody. We used “tailedness” (measured by excess kurtosis) and range of the distributions as proxy measures for average and maximum potential sensitivity, respectively. Since excess kurtosis is, strictly speaking, defined for normal distributions, its interpretive power depends on the nature of the observed parameter. It may be beneficial in future work to consider additional properties of distributions, such as skewness or diversity (as measured by Shannon entropy). Although we only studied general trends in sensitivity from the perspective of individual developability parameters, it would be of interest to explore how a mutation impacts values across all DP of an individual antibody.

In this work, we did not perform structure-based substitution analysis. Specifically, given that there is a lack of experimentally determined structures of antibody variants, it remains unclear how accurately antibody structure prediction tools can resolve single amino acid structural differences and reflect them in models (Supplementary File; Supp. Figure 19) (*111*). More generally, in the context of classical MD simulations of antibodies, the conformational landscape typically exhibits minor variations compared to the initial rigid structure (*112*). In MD simulations of five experimentally determined antibodies without antigen (Supplementary File), we found that the structural fluctuations in the CDRH3 loop regions were generally minimal (on the order of 0.1Å) over the course of 100 ns simulations (as determined by RMSD) (*113*). However, the structural variance between the initial (rigid) antibody conformation and the relaxed version was between 1.2Å and 1.7Å on average (Supplementary File; Supp. Figure 18A). Given the relatively small magnitude of CDRH3 fluctuations and the current limitations of structure prediction methods, it is challenging to detect and accurately represent these small conformational changes in rigid antibody models. The accuracy of current structure prediction tools typically is below the range of these small fluctuations (*114*). However, when it comes to studies involving antibody variants, we cannot overlook these fluctuations, as they reflect the effects of mutations. These computational structure analyses should be combined with experimentally determined antibody structures, such as X-ray or cryo-EM data. While experimental structures contain variations due to differences in crystallization conditions, resolution levels, and inherent protein dynamics, making the same structure slightly different in various studies, computationally predicted structures lack the experimental-related noise found in real structures, causing all models to appear identical outside the mutation site. Given these limitations, we anticipate that the differences in distances between variants, specifically at the mutation site, will be lower than expected in experimental variants and greater than expected in computational models. Our findings suggest that the reality of variant effects lies between the generated models and structures observed experimentally. Furthermore, due to the dynamic nature of proteins, simulated or experimentally determined structural differences of the same protein can overshadow differences caused by mutations, especially in highly flexible antibody loops like CDRH3.

### Predictability of antibody developability is a function of ML method and DP type

ML models have been shown to be able to predict the biophysical characteristics of proteins (*30*, *32*, *115*, *116*). Advances in both experimental data generation, computational structure prediction and ML methods have enabled and improved upon the *in silico* prediction of developability parameters (*23*). While their use may not be widespread in the pharmaceutical industry, the emergence of AI/ ML may become routine as part of initial *in silico* efforts to screen and assess molecular properties and interactions prior to any experimental efforts (*23*). To efficiently determine biological (and, possibly, clinical) properties of antibodies using machine learning-based methods, it is important first to identify a suitable representation. In our study, we compare two alternative embedding types, one (DPL), which collects (a subset of) DPs into a numerical vector and the other (PLM) obtained by encoding the antibody sequence using a pre-trained neural network (NNs). The former representation mirrors feature-selection approaches to DP prediction, while the latter embraces the latest advances in deep NNs and, in particular, transformer-based models. We compute DP predictions by passing DPL and PLM embedding through *linear regressor heads*. As shown in Figure 6B, the predictive power of our models is limited and this holds especially for DPL predictors, which achieve low R^2^ scores even on sequence DPs, which are instead well predicted using the PLM representation. The stark difference in performance is not surprising and can be understood as the combination of two facts. First, we observe that our use of linear heads is a restrictive architectural choice, which is likely over-simplistic: it is natural to expect that most – even though not all – DPs cannot be written as a linear combination of the remaining ones. This limitation is exacerbated by multiple collinear features (as in the full developability profile, Figure 2) and by a reduced number of dimensions (the number of coordinates of DPLs vs PLMs span two orders of magnitude). Second, as visible in Supp. Figure 10C (right), PLM representations retain considerable information about sequence identity and they can be thought of as a learned map from sequences to a latent space. They are, therefore, clearly more suitable to derive sequence DPs and, as our experiments find, also structure DPs, although to a lesser extent. Overall, PLM-based DP predictions generalize well across human-engineered datasets, hinting at a modest amount of overfitting. This, paired with the observation that regressor quality improves with bigger train set sizes (no clear plateaus found for the cardinalities tested in Figure 6B), indicates a healthy learning trend and suggests that predictions may benefit from more advanced head architectures.

DP predictors also shed light on the relationship between different groups of DPs, e.g., between sequence and structure DPs, and MWDS vs. non-MWDS ones. Both DPL- and PLM-based regressors fail in inferring most structure-based parameters and, with few exceptions (notably AbStruc_psi_angle and AbStruc_unfolded_pI), succeed on the same DPs (e.g., AbStruc_loops, AbStruc_beta_strands, AbStruc_sasa, AbStruc_unfolded_pI, AbStruc_pcharge_hetrgen, AbStruc_psi_angle). This seems to suggest the failure of both representations to capture the complex relations between sequences in linear subspaces of the latent space. We also remark that although the lower correlation between structure DPs seen in Figure 2A may suggest that noise introduced by structure prediction tools also contributes to the weak prediction outcomes, we were unable to obtain better scores when predicting structure DPs computed on the (few) available solved antibody structures.

*Comparing the predictability of MWDS and non-MWDS DPs*. MWDS parameters are chosen to avoid high correlations between one another (Figure 2C). Thus, these parameters may be useful hints to know something about the non-MWDS parameters. Although they cannot be used to deduce the exact values of non-MWDS DPs, they are strongly correlated to the latter. Predicting MWDS DPs with DPL models that are trained only on other MWDS DPs therefore tends to be difficult (Figure 6B). This raises the question of whether this difficulty is due to special characteristics of selected MWDS DPs or simply caused by the exclusion of highly correlated DPs. Supporting the latter possibility, with our current approach, the selection of MWDS DPs is non-deterministic and depends on hyperparameters – including the correlation threshold and isotype – which can alter resulting MWDS DP sets. Furthermore, when using PLM embedding, the MWDS prediction performance does not differ from that of non-MWDS parameters, indicating that the difference in DPL performances is a mere consequence of the choice of MWDS DPs. Based only on sequence features, PLM predictions are not biased (neither positively nor negatively) by correlations among DPs. Therefore, we conclude that although a given MWDS represents only one of the (many) non-redundant subsets of all assayed DPs, there is no special relevance to the particular parameters included in the MWDS for PLM-based embedding ML predictions.

Finally, the ML methods employed in this work are by no means exhaustive and only represent a first step toward ML-based DP predictability. It is of interest that already relatively simple ML approaches achieve good prediction accuracy. More generally, our ML approach is to be understood as an example of potential studies that may be performed once it is clearer which DPs are causally linked to downstream antibody candidate success.

### Challenges in the computation of structure-based developability parameters

Our findings suggest that structure-based DPs calculated on different antibody structure prediction models correlate overall poorly (Supp. Figure 17A). These observations align with recent reports by others (*33*, *34*, *114*). Importantly, we found that although ABB2 (*37*) and IgFold (*55*) have been found to be superior to ABB (*36*, *37*, *117*) (which was used in this work for the majority of the results), the correlation of ABB with all other tools (experimental) in terms of structure-based DPs was low and did not differ to the aforementioned tools. Strikingly, we found that structures obtained by computational structure prediction methods belonged to a common underlying ensemble structural distribution as revealed by MD (Supplementary File; Supp. Figure 18). Therefore, the usage of ABB did not significantly bias the results in this paper.

However, it remains challenging to predict the antibody CDRH3 loop regions, which are of significant interest due to their involvement in antibody-antigen binding (*94*) as well as developability (*20*). These loop regions are generally more flexible and less structurally conserved compared to stable secondary structure elements like beta sheets (*114*). Consequently, predicting the specific conformations of these loops accurately can be difficult, and it becomes even more challenging to determine when a prediction is correct without combining experimental validation and MD simulations (*34*, *114*, *118*).

MD is important for a fuller understanding of antibody systems as it provides insight into the flexibility and fluctuations of antibody structure and developability parameters (*29*, *33*, *34*, *117*, *119*, *120*). Specifically, Park and Izadi found that antibody developability surface descriptor parameters (e.g., positive electrostatic potential on the surface of the CDR region (*121*)) vary extensively as a function of the structure prediction method used, which is in line with the findings in this manuscript (Supplementary File). However, after Gaussian-accelerated MD (GaMD) simulations and averaging the values of the descriptors through MD frames, the consistency between the descriptors improved (*34*). Similarly, Raybould and colleagues (*33*) conducted an evaluation of variations in four structure-based Therapeutic Antibody Profiler (TAP) (*7*) scores using molecular dynamics (MD). The mean values of the TAP properties observed during the simulations closely matched an ensemble generated from three TAP predictions based on static Fv models directly generated by ABodyBuilder2. Furthermore, certain structure prediction tools, such as IgFold (*55*), refine the final model using short relaxation using OpenMM (*122*) or PyRosetta (*123*).

In the future, generating multiple models with various refinements could yield a conformational ensemble similar to what is obtained from a full-fledged long MD simulation (*114*, *118*). Training on dynamic data, such as MD simulation-based conformations, enables the model to capture loop flexibility and variability, resulting in more robust and realistic predictions that can improve currently challenging CDRH3 structure prediction. An ideal ML model trained on dynamic data could streamline antibody structure prediction by eliminating the need for additional MD simulations, saving computational resources, and providing a higher level of confidence in predicted structures and insights into hidden antibody characteristics and developability properties.

### Human-engineered antibodies mostly fall within the natural developability and sequence landscapes

Since the development of therapeutic antibodies involves human-directed sequence engineering to attain desirable developability properties, there have been efforts to separate natural from therapeutic antibodies (*7*, *51*). However, we showed that human-engineered (including therapeutic) antibodies fall *within* the developability space of natural antibodies (Figure 7B). Although this could simply be due to the particular selection of DPs and the (major) principal components of variations, we showed that human-engineered antibodies also fall within the *sequence* space of natural antibodies (Supp. Figure 11B). Here, we discuss several possible explanations: (1) Our findings are consistent with (*48*), who have shown considerable sequence overlap between therapeutic antibodies and NGS-derived natural repertoires, indicating that therapeutic antibody sequences are largely derived from natural sequences with few modifications. This could also explain why MLR models trained on natural DPs also predict DPs of human-engineered antibodies with similar accuracy (Figure 7C). (2) Although the development and formulation of therapeutic antibodies involve distinct challenges from natural antibody generation, both might be subject to converging primary restrictions. As shown by (*50*), certain framework mutations regularly occurring in clinical-stage therapeutic antibodies are also frequently observed in natural repertoires, indicating converging selection towards common characteristics such as stability. It is conceivable that the sequence space of stable (and non-immunogenic) antibodies is restricted so that natural and human-engineered sequences occupy overlapping subspaces within. (3) There might be considerable engineering of the Fc region (*124*), which is not included in our study, that could separate human-engineered from natural antibodies in sequence space. (4) Lastly, the number of patent-submitted (*54*, *125*) and therapeutic antibody sequences available is relatively small, which means that there may be potential therapeutics in yet unexplored regions of the sequence space. However, given that sample sizes differ across the human-engineered datasets, more data is needed to study how cross-transferable conclusions are.

Despite being globally similar to the native dataset, DPLs of human-engineered antibodies show distinctive traits. Most notably, they differ in the patterns of correlation found among DPs. This is best seen in Figure 7B, which is obtained by projecting native DPLs along two axes which are determined in order to be uncorrelated. In this subspace, the cluster originated by native antibodies has a circular shape, i.e., it is isotropic, meaning that the knowledge of one of the two coordinates gives limited information about the other. This property, however, does not hold for the set of engineered antibodies, which form an elongated shape, stretching along a diagonal. A similar cluster shape implies that growing x-values tend to correspond to bigger y-values or, in other terms, that there is a (weak) positive correlation between the two axes. Among human-engineered antibodies, then, alterations to the two parts of DPL, which are projected on each axis, do not happen independently anymore, likely signaling the artificial effect on DPs of the antibody optimization process. In the future, it would be of interest to study how this potential signal of the antibody optimization process is represented in different PLM embeddings, for example, in antibody-specific PLM embeddings, for which evidence is conflicting as to whether general protein or antibody-specific PLM more faithfully represent inter-sequence functional similarity (*72*, *126*, *127*)

Furthermore, it is interesting to note that human-engineered DPLs do not cover entirely the native DPL cluster (Figure 7B). Among human-engineered antibodies, it is hence less probable to find some types of DPLs, which appear instead frequently among native ones: this suggests the existence of an unexplored region of DP space. Our conclusion is strengthened by the presence of a similar low-density zone on the right of the cluster of PLM embedding (Supp. Figure 10B), which may constitute evidence of under-investigated classes of antibody sequences.

### Future directions for computational developability profiling

Computational developability profiling, similar to computational antigen binding profiling (*3*, *95*), will be increasingly useful once the real-world relevance of computational DPs has been further established (*31*, *32*, *34*, *97*). To this end, multiple avenues require further development: (i) insight into differences of paired vs single-chain developability (*92*, *93*), (ii) faster computation of structural and MD-based DPs (*128*), (iii) negative controls for clinical stage antibodies that have not progressed due to developability issues, (iv) large-scale data where multiple properties for a given antibody are captured to start exploring multiparameter generative design (*30*, *58*, *104*, *129*), and (v) open and unbiased competitions with agreed-upon quality experimental (and simulated (*64*, *130*)) data to quantify the current state-of-the-art in DP predictability, causal relationships and importance ranking vis-a-vis DP impact on antibody developability.

## Methods

### Antibody datasets

#### The native antibody dataset

We collected a total of 2,036,789 native human and murine antibody sequences (Supp. Figure 1A). Briefly, we assembled non-redundant sequences of human (heavy chains: IgD 173,342, IgM 173,437, IgG 170,473, IgA 171,174, IgE 165,992, and light chains: IgK 198,255, IgL 187,378) and murine (heavy chains: IgM 198,967, IgG 199,326, and light chains: IgK 198,795, IgL 199,650) native antibody variable region sequences (Supp. Figure 1A). Antibody sequences were majorly sourced from Observed Antibody Space (1,738,091 sequences) (*131*) in addition to including our own experimentally-generated sequences (total of 298,698 sequences with the IgD, IgK and IgL human datasets) to provide balanced antibody count among all isotypes. Experimental sequences were generated following a protocol inspired by (*132*), starting from human blood (see Supp. Methods), and exported as VDJ clones from raw sequencing files using MiXCR (version 3.0.1) (*133*). OAS sequences were aligned and IMGT-numbered using the IMGT/High V-QUEST (*134*). The kappa and lambda light chain types (IgK and IgL) were often referred to as isotypes in this manuscript to provide a cohesive reading experience.

### The human-engineered antibody datasets

#### (1) The therapeutic antibody dataset (mAb)

A total of 782 therapeutic antibody sequences belonging to the “whole mAb” format were obtained from TheraSAbDAb as per July 2021 (*135*). First, all mAb sequences were aligned to both murine and human germlines (species of interest) with the antigen receptor alignment and annotation tool ANARCI (*136*). We used the same IMGT-numbering scheme as implemented in the native dataset sequences numbering to ensure consistent methodology. For each chain, we kept the alignment with the highest germline certainty (human or mouse) by selecting for the highest V gene identity (the sequence identity over the V-region to the most sequence-identical germline). In case V gene identity was equal for murine and human alignments, we kept the alignment with the highest J gene identity (the sequence identity over the J-region to the most sequence-identical germline). To ensure the accuracy of annotations, we excluded sequences with V gene identities less than 0.7 (Supp. Figure 1B).

#### (2) The patented antibody database (PAD)

Under a non-commercial agreement with NaturalAntibody, we obtained a total of ≈240K unpaired variable region antibody sequences (Supp. Figure 1C). Sequences were originally extracted from patent documents from intellectual property organizations and bioinformatic databases (WIPO: world intellectual property organization, USPTO: United States Patent and Trademark Office, EBI: European Bioinformatics Institute, DDBJ: DNA Data Bank of Japan), and mined as explained by (*54*). Sequences were provided with species and V-gene annotation metadata and IMGT-numbered. We selected for human and murine sequences and classified them into heavy (IgH) and light (IgK and IgL) chains based on the corresponding V-gene annotation resulting in a final count of 223,613 PAD sequences (Supp. Figure 1B).

#### (3) Humanized mouse dataset (Kymouse)

We obtained 209,452 IgM V_H_ sequences from humanized transgenic mice (Kymouse) as described in (*82*). Sequences were downloaded from OAS (*131*), and their structures were predicted with ABB as explained above. Finally, we also measured their sequence and structure DPs.

### The *in silico* mutated antibody dataset

We firstly sampled 500 wildtype (WT) antibodies from the human native V_H_ dataset (100 samples per isotype for IgM, IgD, IgG, IgA and IgE). We generated all possible single amino acid substitutions for each antibody, resulting in a total of 301,777 sequences for which we predicted/computed their sequence-based developability parameters as previously described. We used this data to perform the sequence DP sensitivity analysis described in Figure 4.

### Antibody structure prediction

We used ABodyBuilder to predict the structures of antibody variable regions (*36*). The high-throughput version of ABodyBuilder is provided as an image for Vagrant VirtualBox (version 2.2.16), known as SabBox. We ran this version with default parameters using unpaired single chain input (heavy or light) to predict all structures in all datasets, unless mentioned otherwise.

### *In silico* calculation of developability parameters

Calculation of sequence-based developability parameters

***Molecular parameters:*** the molecular weight of an antibody variable region sequence was calculated using the Peptides R package version 2.4.4 (*137*, *138*). We calculated the antibody length and the average residue weight using custom R scripts as described by (*139*). ***Amino acid categorical composition:*** Using a custom R script, we calculated the proportional (%) content of amino acid categories as described by (*137*) (Supp. Table 2) by dividing the occurrences of the amino acids in one group by the length of the antibody variable region sequence. ***pI and charge***: To compute the pI and charge of a given sequence, we used the Peptides R package (*137*) using the “Lehninger” scale between pH=1 and pH=14 with step size

=1 (14 data points). *Extinction coefficient and molar extinction coefficient (for all and for cysteine bridges):* Calculations were performed as mentioned in (*140*, *141*) using custom R scripts. *Hydrophobicity and hydrophobic moment*: To compute hydrophobicity and the hydrophobic moment, we used the Peptides R package (*137*), the scale of choice was Eisenberg. For hydrophobic moment calculation, we selected ten amino acids for the length of the sliding window size based on our understanding of the antibody secondary structure and as explained by (*142*). We also specified the angle value as 160 as recommended by (*143*). Instability index and aliphatic index: The aliphatic index is defined as the relative volume occupied by aliphatic side chains (Alanine, Valine, Isoleucine, and Leucine – Supp. Table 2). It may be regarded as a positive factor for the increase of thermostability of globular proteins (*144*). The instability index was first developed by (*145*) to reflect the stability of proteins based on their content of certain dipeptides that were found to be associated with degradation tendency. We calculated the instability and aliphatic indices using the Peptides R package (*137*). *Protein solubility prediction:* We used SoluProt (version 1.0) to predict the solubility index for antibody variable region sequences (*146*). We chose this tool as it allows the consideration of protein expressibility into its solubility score prediction, while proving comparable performance compared to other state-of-the-art solubility prediction tools (*146*). Sequences with solubility scores above 0.5 were predicted to be soluble when expressed in *Escherichia Coli*. *Immunogenicity prediction:* We used netMHCIIpan version 4.0 (*39*) to estimate the immunogenicity of the antibody variable region sequences. Briefly, we examined the global immunogenicity of the antibody sequences by predicting their affinities for HLAII supertypes that are found in 98% of the global population (*108*). We obtained the following numerical values from these calculations including (i) minimum rank, (ii) number of weak binders (percentage rank >2 & <10), (iii) number of strong binders (percentage rank <2), (iv) full average percentage rank and (v) the number of antibody regions where the maximal immunogenic peptide stretches (maximal_immunogenicity_region_span). As these calculations are computationally intensive, we computed the immunogenicity DPs on 10% only of the native, Kymouse and PAD datasets.

### Calculation of structure-based developability parameters

First, we added hydrogen atoms to each antibody variable region structure that we previously built with ABodyBuilder using the Reduce software (version 3.24.130724) (*147*). Hydrogenated structures were used as an input (pdb format) to calculate structure-based developability parameters as they were reported to provide closer chemical representation of their potential in vivo structures (*148*). As detailed in Supp. Table 1, we used BioPython (version 1.79) to compute secondary structure parameters (*149*), FreeSASA (version 2.1.0) for solvent accessibility predictions (*150*), PROPKA (version 3.4.0) for the calculation of thermodynamic and electrochemical parameters (*151*) and ProDy (version 2.0) to count free and bridged cysteines (*152*). We used custom python scripts for the calculation of developability index (DI) and spatial aggregation propensity (SAP) as described by Lauer and colleagues (*52*, *153*). We used the original Black-Mould hydrophobicity scale as suggested by (*120*). Finally, we used Arpeggio (version 1.4.1) to calculate interatomic interactions after converting antibody hydrogenated structures to cif format (*154*).

### Parameter correlation calculation and visualization

For the analysis in Figure 2B, Supp. Figure 2 and Supp. Figure 3, we computed pairwise developability parameter correlations (Pearson) matrices using the ‘cor()’ function from the ‘base’ package in R (version 4.0.3). for each isotype/species combination to investigate parameter associations and redundancies. Missing data was accounted for by choosing the argument *use.pairwise.obs=complete* option in the function. We visualized the correlation matrices using ComplexHeatmap R package version 2.9.4 (*155*) and annotated the heatmaps with the threshold-specific ABC-EDA/MWDS algorithm output (*156*).

### Determination of the minimum weight dominating set of developability parameters

To investigate the redundancy of developability parameters (DPs), we first constructed undirected weighted network graphs for each species and isotype where the nodes represent DPs and the edge weights represent the pairwise Pearson correlation values (as in Figure 2C). For this purpose, we constructed and visualized networks with Cytoscape 3.9.1 (*157*, *158*). The constructed networks could include up to three distinct classes of pairwise relationships for a given correlation threshold (from 0.1 and 0.9 in intervals of 0.1): (1) isolated nodes representing DPs that have correlation values below the given threshold with all other DPs, (2) doublets (exclusive pairs) where two DPs are solely correlated with each other (above the given threshold), and (3) correlation subnetworks where more than two DPs form correlation clusters.

Secondly, the minimum weight dominating set (MWDS) for a specific network at a given correlation threshold was defined as the set of parameters for which the sum of the associated edge weights is minimal (*56*). Thus, the MWDS is a subset of nodes where each node in the network is either in the MWDS or connected to a member of it by an edge (*56*). Based on this definition, we included all DPs from classes (1) and (2) in the MWDS, where no meaningful selection based only on correlation can be made. However, finding the dominating parameters among class (3) DPs is an NP-hard problem. Thus, we implemented an algorithm described by (*56*), which leverages the local optimization of an artificial bee colony algorithm guided by iterative global estimations of distribution (ABC-EDA algorithm) to approximate optimal solutions (*56*). Ultimately, this algorithm classifies class 3 DPs as either “dominant” or “redundant”. The dominant DPs were added to the final MWDS.

### Correlation distance dendrograms

This analysis aims to examine the similarities in the pairwise associations of developability parameters among isotypes and species of the native and human-engineered antibody datasets (Figure 3B and Figure 7A). For this analysis, we transformed the isotype- and species-specific parameter correlation matrices (sequence and structure parameters separately) into numerical vectors. Subsequently, we quantified the pairwise Pearson correlation distance for these numerical vectors using the ‘get_dist’ function from the R package factoextra version 1.0.7 (*159*). We finally clustered the resulting distance matrices using the hierarchical clustering ‘hclust’ command following the complete linkage method from the stats package, version 4.0.3 (*160*), where the height of the dendrogram represents the Pearson distance (0–1).

### Analysis of parameter sensitivity to single amino acid substitution: excess kurtosis and range Excess kurtosis

To investigate the impact of changes in the amino acid sequence on DP values (sensitivity - analysis in Figure 4 and Supp. Figure 12), we normalized (mean-centered) the DP values of the mutated antibody dataset by subtracting their mean and then dividing by their standard deviation. The DP distributions of all mutants of each wt-antibody were analyzed by calculating and comparing two µ metrics First, the excess kurtosis, defined as 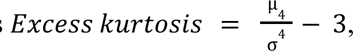, where µ is the fourth central moment and σ the standard deviation of the normalized DP values. An excess kurtosis of 0 represents that of a normal distribution. It increases with peakedness and decreases with uniformity of the distribution, under the assumption that it is bell-shaped. Thus, while a high excess kurtosis indicates a small change induced by single amino acid substitutions on average or low average sensitivity, low excess kurtosis suggests high average sensitivity. The second metric is the *range* of a distribution, defined as the distance between the highest and lowest DP values and represents the maximum normalized shift that is inducible by a single amino acid substitution.

### Pairwise developability profile correlations and pairwise sequence similarity studies

We define the ***antibody developability profile (DPL)*** as a numerical vector that carries the values of developability parameters in a fixed order for a given antibody sequence. Developability parameter values were mean-centered and scaled to unit variance (normalized) as previously described (*161*). Of note, scaling for a given group of antibody sequences was performed after accounting for chain type and species.

For the analysis in Figure 5A,B, we defined the ***pairwise developability profile correlation (DPC)*** as the Pearson correlation value between a pair of sequence developability profiles. We computed the amino acid-based sequence similarity between two antibody variable region sequences (A and B) as follows:

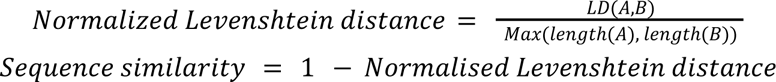

Where LD(A,B) represents the Levenshtein distance between A and B. We used the stringdist R package version 0.9.8 (*162*) and the Levenshtein python package version 0.20.8 to calculate LD (*163*).

In Figure 5C we inspected the relationship between sequence similarity and normalized developability profile (DPL) similarity by firstly examining the proximity of human heavy chain antibodies that belong to the same sequence similarity cluster on the 2D PCA projection plane of the developability space (*R^N^*), where *N* is the number of DPs that makes a developability profile. In this analysis, we used the MWDS DPs (which have full values for all antibodies) as identified by the ABC-EDA algorithm for the human IgG dataset at absolute Pearson correlation coefficient threshold of 0.6, accounting for 46 DPs (*N* = 43, Supp. Table 3). The PCA was computed using the python packages scikit-learn version 1.1 (*164*) and dask-ml version 2022.5.27 (*165*). Sequence similarity groups were identified using USEARCH version 11.0 (*166*) as groups of 10000 (or more) antibodies that share at least 0.75 sequence similarity as defined by Levenshtein distance (Supp. Figure 9B).

Then, to quantify the relationship between sequence similarity and developability profile similarity, we studied the correlation between the pairwise Euclidean distance (ED) in the developability space (*R^N^*) and the pairwise normalized LD for a sample of 5000 human IgM antibodies as in Supp. Figure 9C (computationally intensive process; 12,500,000 data points for each ED and LD calculation). We ensured that these sampled antibodies belong to the same IGHV gene family to exclude the effect of IGHV family variance on the LD computations (*61*, *62*). Antibody pairs with ED ≥ 15 were considered as outliers and were excluded from this analysis (forming 0.8% of total data points).

Of note, the PCAs from this analysis were used to examine the positioning of the human human-engineered antibody datasets within the developability space of the human native antibodies (Figure 7B) and the role of native antibody isotype and germline gene annotation in the positioning within the developability space Supp. Figure (Supp. Figure 10A, Supp. Figure 11A).

### Predictability of developability parameters

We used the human V_H_ antibody sequences (854,418 antibodies) and their computed DP values (normalized; mean-centered) to assess the predictability of developability parameters in two machine-learning tasks (ML task 1 and ML task 2). Of note, we excluded mouse antibodies and V_L_ sequences to avoid species- and chain-specific biases, ensuring greater data homogeneity. In both tasks, the predictability of DPs was assessed by computing the coefficient of determination (R^2^) between observed and predicted DP values (*78*).

### ML task 1; predicting the values of the same (single) missing DP

For this task (Figure 6A,B, and Supp. Figure 14A), we randomly split the human V_H_ antibody sequences into two subsets: (i) ***training set***; containing ∼80% of sequences (683,534), which was further subsampled to derive training sets of variable sizes (50, 100, 500, 1000, 10000, 20000). Specifically, for each size, we defined 20 independent training subsets (ii) ***test set;*** containing the remaining (∼20%) sequences (170,884). Then, we used two types of embeddings to train multiple linear regression (MLR) models: (1) single-DP-wise incomplete developability profiles (low-dimensionality embeddings – order of 10s) and (2) antibody sequence encodings obtained from the protein language model (PLM) ESM-1v (high dimensionality embeddings – order of 1000s (*69–71*)). Beginning-of-sentence (BOS) processing was used to compress the vectors produced by the PLM to avoid biases that might occur by averaging the entire vectors (as detailed below in “Antibody sequence encoding with PLM”). Finally, we compared the predictive power of both embeddings to predict the values of DPs in the test set after deleting single-DP column values at a time. For each type of embedding, we predicted the missing DP using linear regressors, trained with the MSE loss. We did not opt for more complex types of regressors in order to contrast the occurrence of overfitting. The DP value prediction was repeated 20 times (using the 20 independently trained MLR models per training set size) and the mean R^2^ was reported.

The native-trained MLR models from this task were then utilized to predict the values of MWDS DPs (Supp. Table 3) in the human-engineered datasets (shown in Figure 7C). Models were trained using the training set size which achieved the highest prediction accuracy before plateauing (1000 antibodies for DPL-based predictions 20K antibodies for PLM-based predictions).

### ML task 2; predicting the values of randomly missing DPs

For this task (Figure 6A,C and Supp. Figure 14B), we randomly deleted (either 2% or 4% of) DP values from subsamples of the human V_H_ antibody sequences (683,534 sequences defined as training set in ML Task 1). We then predicted the deleted (missing) data using the multivariate imputation by chained random forests (MICRF) algorithm (*74*) via the missRanger R package (*167*). We repeated this step 20 times for each subsample size (50, 100, 500, 1000, 10000, 20000 antibodies) and reported the mean R^2^.

For both ML tasks – and both embedding types implemented in ML Task 1 – we performed ablation studies by randomly permuting the column values in the input datasets for the ML models, and confirmed that the prediction accuracy was abolished (R^2^ ≤ 0).

### Antibody sequence encoding with PLM

Antibody sequences were encoded using a Protein Language Model (PLM). PLMs are deep neural networks designed to transform protein sequences into contextual embedding vectors depending on the entirety of the protein sequence. In our experiments, we use the PLM ESM-1v since this model is optimized to predict the effects of mutations on the function of proteins (*70*).

To extract a global, fixed-size representation from PLM embeddings, we use a compression scheme based on the Beginning-Of-Sequence (BOS) token, as is customary for LLMs, e.g., for the [CLS] token in (*168*). BOS tokens are trained to provide a *summarized* representation of the entire protein sequence and are therefore a natural choice to represent antibodies.

### Graphical illustrations

We used BioRender.com to create the illustrations in Figure 1, Figure 4A, and Figure 6A. Antibody structural images were produced in PyMOL v2.5.5 (*169*). We generated the remaining figures in RStudio (*138*) using the ‘ggplot2’ package (*170*) and figure panels were aggregated with Adobe Illustrator (*171*).

## Acknowledgements

We acknowledge the help of Haidara Nadwa (University of Siena) for providing valuable insights in regard to the MD analysis.

## Author contributions

V.G. conceived the study. H.B. and V.G. conceptualized the study and designed the experiments. H.B., E.S., M.P. and J.Z. performed all data analysis and visualization, and developed the project online repositories. H.B., E.S., M.P., J.Z. and V.G. wrote the first draft of the manuscript. H.B. and E.S. prepared sequence and structural developability data (respectively) for all sections of the manuscript. E.S. and R.A. performed antibody structural modeling and molecular dynamic simulations. J.Z. coded and ran the ABC-EDA hybrid algorithm. M.P. and D.N-Z.G. generated sequence encodings with the protein language model ESM-1v. R.A. and M.S. provided advice on experimental design and statistical analysis. K.L.Q. and I.S. generated the experimental BCR sequences included in this study. K.K. provided the patented antibody dataset (PAD) under a non-disclosure agreement with V.G. H.B. and V.G. made significant contributions to the final version of the text and figures. All authors revised the manuscript and approved its content.

## Data and materials availability

Our code and low-size data (scripts and figures) are available on GitHub: https://github.com/csi-greifflab/developability_profiling. Additional structural data, such as predicted models and MD trajectories, are stored on Zenodo: 10.5281/zenodo.10013525.

## Funding

The Leona M. and Harry B. Helmsley Charitable Trust (#2019PG-T1D011, to VG), UiO World-Leading Research Community (to VG), UiO: LifeScience Convergence Environment Immunolingo (to VG and GKS), EU Horizon 2020 iReceptorplus (#825821) (to VG), a Norwegian Cancer Society Grant (#215817, to VG), Research Council of Norway projects (#300740, #311341, #331890 to VG), a Research Council of Norway IKTPLUSS project (#311341, to VG and GKS), and Stiftelsen Kristian Gerhard Jebsen (K.G. Jebsen Coeliac Disease Research Centre, SKGJ-MED-017) (to GKS), and BBSRC (BB/V011065/1, to JG-M). This project has received funding from the Innovative Medicines Initiative 2 Joint Undertaking under grant agreement No 101007799 (Inno4Vac). This Joint Undertaking receives support from the European Union’s Horizon 2020 research and innovation programme and EFPIA (to VG). This communication reflects the author’s view and neither IMI nor the European Union, EFPIA, or any Associated Partners are responsible for any use that may be made of the information contained therein.

## Competing interest statement

V.G. declares advisory board positions in aiNET GmbH, Enpicom B.V, Absci, Omniscope, and Diagonal Therapeutics. V.G. is a consultant for Adaptyv Biosystems, Specifica Inc, Roche/Genentech, immunai, Proteinea and LabGenius. K.K. is the founder of NaturalAntibody. M.P. and D.N-Z.G. are employed by Adaptyv Biosystems.

## Abbreviations

2D: two dimensional
3D: three dimensional
aa(s): amino acid(s)
Ab(s): antibody(ies)
ABB(2): ABodyBuilder(2)
ABC-EDA: artificial bee colony and estimation of distribution algorithm
AF: AlphaFold
ANARCI: antigen receptor numbering and receptor classification
BOS: beginning of sequence
(c)DNA: (complementary) deoxyribonucleic acid
CDR: complementarity-determining region
DI: developability index
DP(s): developability parameter(s)
DPC: developability profile correlation
DPL: developability profile
ED: Euclidean distance
FR: antibody framework region
Fv: antibody variable fragment
Ig: immunoglobulin
IMGT: immunogenetics information system
LD: Levenshtein distance
mAb(s): monoclonal antibody(ies)
MD: molecular dynamics
MICRF: multivariate imputation by chained random forests
ML: machine learning
MLR: multiple linear regression
MWDS: minimum weighted dominating set
ns: nanosecond
NPT: constant particle number (N), constant pressure (P), constant temperature (T)
NVT: constant particle number (N), constant volume (V), constant temperature (T)
OAS: observed antibody space
PAD: patented antibody dataset
PC(A): principal component (analysis)
PDB: protein data bank
PLM: protein language model
PME: particle mesh Ewald
ps: picosecond
R^2^: coefficient of determination
RMSD: root mean square deviation
SAP: spatial aggregation propensity
SEM: standard error of the mean
TAP: therapeutic antibody profiler
Thera-SAbDAb: Therapeutic Structural Antibody Database
TIP3P: transferable intermolecular potential with 3 points
V-rescale: velocity-rescaling
V_H_: antibody heavy chain variable domain
V_L_: antibody light chain variable domain

## Notes

https://github.com/csi-greifflab/developability_profiling

### Summary of Updates

Updated the title and the abstract of the manuscript.

