## supplementary information for "Biophysical cartography of the native and human-engineered antibody landscapes quantifies the plasticity of antibody developability"

#### Supplementary Tables

**Supp. Table 1 | Sequence and Structure developability parameters.** In this table, we show the developability parameters we included in our study, alongside their physicochemical classification and method of calculation.

| **Physicochemical property** | **Parameter (unit)** | **Nomenclature** | **Method of calculation** | **Description** |
|---|---|---|---|---|
| | | **Sequence parameters** | | |
| Molecular | Molecular weight (Daltons) | AbChain_mw | Peptides R package | - |
| | Sequence length (aa) | AbChain_length | R script nchar(seq) | - |
| | Average residue weight (Daltons) | AbChain_av_resdue_weight | R script mw/(nchar(seq) - 1) | - |
| Amino acid composition | Content (%) of aromatic amino acids | AbChain_aromatic_content | R script according to classification scheme in the Peptides R package manual | F+H+W+Y |
| | Content (%) of tiny amino acids | AbChain_tiny_content | | A+C+G+S+T |
| | Content (%) of small amino acids | AbChain_small_content | | A+C+D+G+N+P+S+T+V |
| | Content (%) of aliphatic amino acids | AbChain_aliphatic_content | | A+I+L+V |
| | Content (%) of non-polar amino acids | AbChain_nonpolar_content | | A+C+F+G+I+L+M+P+V+W+Y |
| | Content (%) of polar amino acids | AbChain_polar_content | | D+E+H+K+N+Q+R+S+T |
| | Content (%) of basic amino acids | AbChain_basic_content | | H+K+R |
| | Content (%) of acidic amino acids | AbChain_acidic_content | | D+E |
| Electrochemi | Charge | AbChain_digit_char | Peptides R package | - |

| | | | | |
|---|---|---|---|---|
| cal | | ge<br>digit = [1-14] | | |
| | Isoelectric point | AbChain_pI | Peptides R package | The pH where positive (+) charges equal to negative (-) charges |
| | Hydrophobicity | AbChain_hydrophobicity | Peptides R package | Eisenberg hydrophobicity |
| | Hydrophobic moment | AbChain_hmom | Peptides R package | Maximal hydrophobicity of a sliding window of 10 aa |
| | Solubility (0-1 score: 0.5 threshold) | AbChain_solubility | SoluProt software | Solubility and expressibility prediction. A 0-1 scale. 0.5 is the threshold |
| Photochemical | Molar extinction coefficient ( $cm^{-1} M^{-1}$) | AbChain_molextcoef | Custom R script as per (*140*, *141*) | The MEC of the full sequence |
| | Molar extinction coefficient of cystine bridges ( $cm^{-1} M^{-1}$) | AbChain_cysbridges_molextcoef | Custom R script as per (*140*, *141*) | The MEC of the cystine bridges found in the full sequence |
| | Molecular extinction coefficient ($gr^{-1} cm^{-1}$ ml) | AbChain_percentextcoef | [molar extinction coefficient X 10 ] / molecular weight of the peptide | Same as above, normalized by molecular weight |
| | Molecular extinction coefficient of cystine bridges ($gr^{-1} cm^{-1}$ ml) | AbChain_cysbridges_percenextcoef | [molecular extinction coefficient of cystine bridges X 10 ] / molecular weight of the peptide | Same as above, normalized by molecular weight |
| Stability | Instability index | AbChain_instaindex | Peptides R package | This index can reflect in vivo half-life |
| | Aliphatic index | AbChain_aliphindex | Peptides R package | Indicator of thermal stability |
| Immunogenicity | Maximal immunogenicity read | AbChain_min_rank | netMHCIIpan | The lowest the rank, the more immunogenic the peptide is |
| | Number of peptides with strong binding affinity to HLAII alleles (rank <2%) | AbChain_num_strong_binders | netMHCIIpan | - |
| | Number of peptides with weaker binding affinity to HLAII alleles (rank [2-10]%) | AbChain_num_weak_binders | netMHCIIpan | - |

| | | | | |
|---|---|---|---|---|
| | The immunogenicity average rank | AbChain_full_average_rank | netMHCIIpan | The average immunopeptidome affinity to all the HLAII molecules included in the analysis |
| | The span of maximal immunogenic peptide | AbChain_immunopeptide_regions_span | netMHCIIpan | The count of regions where the maximal immunogenic peptide stretches in each sequence |
| **Structure parameters** | | | | |
| Interaction | Steric clashes | AbStruc_steric_clashes | Arpeggio | Number of atoms which are involved in a steric clash. |
| | Covalent bonds | AbStruc_covbonds | Arpeggio | Number of atoms that are supposed to be covalently bonded |
| | Van der Waals clashes | AbStruc_vdw_clashes | Arpeggio | Number of atoms which van der Waals radius is clashing with one or more other atoms. |
| | Van der Waals interactions | AbStruc_vdw_interactions | Arpeggio | Number of atoms whose van der Waals radius is interacting with one or more other atoms |
| | Proximal interactions | AbStruc_proximal_interactions | Arpeggio | Number of atoms involved in proximal interactions with one or more atoms |
| | Hydrogen bonds | AbStruc_hbonds | Arpeggio | Number of atoms that form a hydrogen bond. |
| | Weak hydrogen bonds | AbStruc_weak_hbonds | Arpeggio | Number of atoms that form a weak hydrogen bond |
| | Ionic bonds | AbStruc_ibonds | Arpeggio | The number of atoms which may interact via charges. |
| | Aromatic interactions | AbStruc_aromatic_interactions | Arpeggio | The number of atoms which an aromatic ring atom **interacts** with another aromatic ring atom |
| | Hydrophobic interactions | AbStruc_hydrophobic_interactions | Arpeggio | The number of hydrophobic interactions within the molecule of interest |
| | Carbonyl interactions | AbStruc_carbonyl_interactions | Arpeggio | Number of carbonyl-carbon:carbonyl-carbon **interactions**. |
| | Polar bonds | AbStruc_pbonds | Arpeggio | The number of atoms that are less strict hydrogen bonding |
| | Weak polar bonds | AbStruc_weak_pbonds | Arpeggio | The number of atoms involved in less strict **weak** hydrogen **bonding** |
| | Atom-atom summary | AbStruc_atom_atom_sum | Arpeggio | This parameter denotes the **summary** number of clash, covalent, vdw, vdw_clash, proximal, hbond, weak_hbond, ionic, aromatic, polar, hydrophobic, carbonyl, and weak_polar interactions. |
| | Carbon-π interactions | AbStruc_carbon_pi_interactions | Arpeggio | This parameter represents the number of weakly electropositive carbon atom-π interactions. |
| | Cation-π | AbStruc_cation_pi_interactions | Arpeggio | This parameter represents the number of cation-π interactions. |

| | | | | |
|---|---|---|---|---|
| | Donor-π | AbStruc_donor_pi_interactions | Arpeggio | This parameter represents the number of **hydrogen bond donor-π interactions** |
| | Sulphur-π interactions | AbStruc_metsulphur_pi_interactions | Arpeggio | This parameter denotes the number of methionine sulphur-π ring **interactions** |
| | Atom-plane summary | AbStruc_atom_plane_sum | Arpeggio | This parameter denotes the summary number of carbon_pi, cation_pi, donor_pi and metsulphur_pi |
| | Amide-amide interactions | AbStruc_amide_amide_interactions | Arpeggio | This parameter denotes the number of side chains **formamide groups - π aromatic ring interactions** |
| | Amide-ring interactions | AbStruc_amide_ring_interactions | Arpeggio | This parameter denotes the number of **side-chain amino** (only the positively charged NH interactions, C=O not included) groups that interact **with the aromatic side-chains ring electron cloud**. |
| | Plane-group summary | AbStruc_plane_group_interactions | Arpeggio | AbStruc_amide_amide_interactions + AbStruc_plane_group_interactions |
| | Planar interactions | AbStruc_plane_plane_interactions | Arpeggio | This parameter denotes the number of **interactions** between atoms in planar side chains (i.e. aromatic residues, and sulfide planes) |
| | Total interactions summary | AbStruc_total_interactions | Arpeggio | This parameter denotes the total of all above interactions. (atom-atom + atom-plane + plane-group + plane-plane) |
| | Average interaction distance (Angstroms) | AbStruc_mean_interaction_distance | Arpeggio | This parameter denotes the **average** pairwise distance between all interacting atoms |
| Secondary structure | φ-angle (degrees) | AbStruc_phi_angle | Biopython | This parameter denotes the **average value of φ-angle** |
| | ψ-angle (degrees) | AbStruc_psi_angle | Biopython | This parameter denotes **the average value of ψ-angle** |
| | Isolated beta bridges residue count | AbStruc_beta_bridges | Biopython | This parameter denotes the number of **residues** which were defined as isolated β-bridge |
| | Beta strands residue count | AbStruc_beta_strands | Biopython | This parameter denotes the number of **residues** which were defined as β-strands. |
| | $3_{10}$ helices residue count | AbStruc_threeten_helices | Biopython | This parameter denotes the number of **residues** which were defined as $3_{10}$ helices. |
| | α-helices residue count | AbStruc_alpha_helices | Biopython | This parameter denotes the number of **residues** which were defined as α-helices. |
| | π-helices residue count | AbStruc_pi_helices | Biopython | This parameter denotes the number of **residues** which were defined as π-helices. |
| | β-bends residue count | AbStruc_beta_bends | Biopython | This parameter denotes the number of **residues** which were defined as β-bends. |
| | Loops residue count | AbStruc_loops | Biopython | This parameter denotes the number of **residues** which were defined as loops |

| | | | | |
|---|---|---|---|---|
| Thermodynamic | Free folding energy (kcal/mol) | AbStruc_folding_energy | PROPKA | This parameter denotes free energy of folding |
| Electrochemical | Unfolded charge | unfolded_charge AbStruc_unfolded_charge | PROPKA | This parameter denotes the protein charge of the unfolded state at pH = 7 |
| | Folded charge | folded_charge AbStruc_folded_charge | PROPKA | This parameter denotes the protein charge of the folded state at pH = 7 |
| | Negative charge heterogeneity | AbStruc_ncharge_hetrgen | Custom Python script | The average pairwise distances between the negatively charged residues and the protein center of mass |
| | Positive charge heterogeneity | AbStruc_pcharge_hetrgen | Custom Python script | The average pairwise distances between the positively charged residues and the protein center of mass |
| | Structural isoelectric point of folded protein | AbStruc_pI_folded | PROPKA | - |
| | Structural isoelectric point of unfolded protein | AbStruc_pI_unfolded | PROPKA | - |
| Druggability | Developability Index (DI) | DI AbStruc_lauer_DI | Custom Python script | This parameter denotes the developability index from Lauer et al., 2012 |
| | Spatial aggregation propensity (SAP) | SAP AbStruc_lauer_SAP | Custom Python script | This parameter denotes the structural aggregation propensity from Lauer et al., 2012 |
| AA composition | Free cysteine count | AbStruc_free_cys | Prody | This parameter denotes the number of cysteines that **do not** form a disulfide bridge |
| | Bridged cysteine count | AbStruc_cys_bridges | Prody | This parameter denotes the number of cysteines that form a disulfide bridge |
| Solvent accessibility | Solvent-accessible surface area (SASA) | AbStruc_sasa | FreeSASA | The surface area of a molecule that is accessible to a solvent. |
| | SASA of side chains | AbStruc_schains_sasa | FreeSASA | This parameter denotes the average SASA over all **side-chain** atoms. |

Supp. Table 2 | **Amino acid categorization based on physicochemical properties as defined by (*137*).** This classification was used as a reference in the computation of several sequence-based DPs.

| Category | Amino acids |
|---|---|
| Aromatic | F+H+W+Y |
| Tiny | A+C+G+S+T |
| Small | A+C+D+G+N+P+S+T+V |

| | |
|---|---|
| Aliphatic | A+I+L+V |
| Nonpolar | A+C+F+G+I+L+M+P+V+W+Y |
| Polar | D+E+H+K+N+Q+R+S+T |
| Basic | H+K+R |
| Acidic | D+E |

Supp. Table 3 | **Minimum weight dominating set (MWDS) DPs as identified by the ABC-EDA for the full set (170,473) of human IgG antibodies at a Pearson correlation coefficient threshold of 0.6.** Doublet parameters (a pair of DPs, see Methods) are mentioned within the same cell and only one (randomly selected) is carried forward to perform the PC analysis (Figure 5C, Figure 7B, Supp. Supp. Figure 10A, Supp. Figure 11A) and the predictability analysis in Figure 6. DPs with incomplete values (**bolded**) were excluded from these analyses as well.

| **Sequence DPs** | | | | |
|---|---|---|---|---|
| AbChain_small_content | AbChain_10_charge" | AbChain_hydrophobicity | AbChain_acidic_content | AbChain_aromatic_content |
| AbChain_length<br>+<br>AbChain_mw | AbChain_percentextcoef<br>+<br>AbChain_molextcoef | AbChain_cysbridges_molextcoef<br>+<br>AbChain_cysbridges_percenextcoef | AbChain_aliphatic_content<br>+<br>AbChain_aliphindex | AbChain_polar_content<br>+<br>AbChain_nonpolar_content |
| AbChain_hmom | AbChain_solubility | AbChain_instaindex | **AbChain_min_rank** | **AbChain_immunopeptide_regions_span** |
| **AbChain_num_strong_binders** | | | | |
| **Structure DPs** | | | | |
| AbStruc_plane_group_interactions | AbStruc_cation_pi_interactions | AbStruc_metsulphur_pi_interactions | AbStruc_sasa | AbStruc_vdw_clashes |
| AbStruc_pbonds | AbStruc_carbon_pi_interactions | AbStruc_unfolded_pI | AbStruc_loops | AbStruc_beta_bridges |
| AbStruc_beta_strands | AbStruc_alpha_helices | AbStruc_beta_bends | AbStruc_carbonyl_interactions | AbStruc_steric_clashes |
| AbStruc_covbonds | AbStruc_hbonds | AbStruc_ibonds | AbStruc_phi_angle | AbStruc_psi_angle |
| AbStruc_vdw_interactions | AbStruc_folding_energy | AbStruc_free_cys | AbStruc_cys_bridges | AbStruc_pcharge_hetrgen |
| AbStruc_ncharge_hetrgen | AbStruc_threeten_helices<br>+<br>AbStruc_beta_turns | AbStruc_aromatic_interactions<br>+ | | |

| | | |
|---|---|---|
| | | AbStruc_plane_plane_interactions |

Supp. Table 4 | **Primers used for the generation of human IgD, IgK and IgL (experimental sequences).**

| Primer name | Sequence |
|---|---|
| Hu_IgD | GGAGTTCAGACGTGTGCTCTTCCGATCTHHHHHACAHHHHHACAHHHHGGGTGTCTGCACCCTGATA |
| Hu_IgK | GGAGTTCAGACGTGTGCTCTTCCGATCTHHHHHACAHHHHHACAHHHHNGGGATAGAAGTTATTCAGCAGGCACACAACAGAG |
| Hu_IgL | GGAGTTCAGACGTGTGCTCTTCCGATCTHHHHHACAHHHHHACAHHHHTGGCTTGRAGCTCCTCAGAGGAGG |
| Read2U | GGAGTTCAGACGTGTGCTCTTCCGATCT |
| Hu_VH_MTPX_1 | CCTACACGACGCTCTTCCGATCTGGTGGCAGCAGTCACAGATGCCTACTC |
| Hu_VH_MTPX_2 | CCTACACGACGCTCTTCCGATCTGGTGGCAGCAGCCACAGGTGCCCACTC |
| Hu_VH_MTPX_3 | CCTACACGACGCTCTTCCGATCTGGTGGCAGCAGCTACAGGTGTCCAGTC |
| Hu_VH_MTPX_4 | CCTACACGACGCTCTTCCGATCTGGTGGSAGCAGCAACARGWGCCCACTC |
| Hu_VH_MTPX_5 | CCTACACGACGCTCTTCCGATCTGCTGGCTGTAGCTCCAGGTGCTCACTC |
| Hu_VH_MTPX_6 | CCTACACGACGCTCTTCCGATCTCCTGCTGCTGACCAYCCCTTCMTGGGTCTTGTC |
| Hu_VH_MTPX_7 | CCTACACGACGCTCTTCCGATCTCCTGCTACTGACTGTCCCGTCCTGGGTCTTATC |
| Hu_VH_MTPX_8 | CCTACACGACGCTCTTCCGATCTGGGTTTTCCTCGTTGCTCTTTTAAGAGGTGTCCAGTG |
| Hu_VH_MTPX_9 | CCTACACGACGCTCTTCCGATCTGGGTTTTCCTTGTTGCTATTTTAAAAGGTGTCCARTG |
| Hu_VH_MTPX_10 | CCTACACGACGCTCTTCCGATCTGGATTTTCCTTGCTGCTATTTTAAAAGGTGTCCAGTG |
| Hu_VH_MTPX_11 | CCTACACGACGCTCTTCCGATCTGGGTTTTCCTTKTKGCTATWTTAGAAGGTGTCCAGTG |
| Hu_VH_MTPX_12 | CCTACACGACGCTCTTCCGATCTGGTGGCRGCTCCCAGATGGGTCCTGTC |
| Hu_VH_MTPX_13 | CCTACACGACGCTCTTCCGATCTCTGGCTGTTCTCCAAGGAGTCTGTG |
| Hu_VH_MTPX_14 | CCTACACGACGCTCTTCCGATCTGGCCTCCCATGGGGTGTCCTGTC |
| Hu_VH_MTPX_15 | CCTACACGACGCTCTTCCGATCTGGTGGCAGCAGCAACAGGTGCCCACT |
| Hu_VH_MTPX_16 | CACTCTTTCCCTACACGACGCTCTTCCGATCTATGGAACTGGGGCTCCGCTGGGTTTTCC |
| Hu_VH_MTPX_17 | CACTCTTTCCCTACACGACGCTCTTCCGATCTATGGACTGCACCTGGAGGATCCTCCTC |
| Hu_VH_MTPX_18 | CACTCTTTCCCTACACGACGCTCTTCCGATCTTGGCTGAGCTGGGTTTYCCTTGTTGC |
| Hu_VH_MTPX_19 | CACTCTTTCCCTACACGACGCTCTTCCGATCTGGAGTTKGGGCTGMGCTGGGTTTTCC |
| Hu_VH_MTPX_20 | CACTCTTTCCCTACACGACGCTCTTCCGATCTGCACCTGTGGTTTTTCCTCCTGCTGGTG |
| Hu_VH_MTPX_21 | CACTCTTTCCCTACACGACGCTCTTCCGATCTCACCTGTGGTTCTTCCTCCTSCTGG |
| Hu_VH_MTPX_22 | CACTCTTTCCCTACACGACGCTCTTCCGATCTCCAGGATGGGGTCAACCGCCATCCTC |
| Hu_VH_MTPX_23 | CTCTTTCCCTACACGACGCTCTTCCGATCTCAGAGGACTCACCATGGAGTTTGGGCTGAG |
| Hu_VH_MTPX_24 | CCTACACGACGCTCTTCCGATCTGGACTCACCATGGAGTTGGGACTGAGC |
| Hu_VH_MTPX_25 | CCTACACGACGCTCTTCCGATCTGGGCTGAGCTGGCTTTTTCTTGTGGC |
| Hu_VK_MTPX_1 | CTACACTCTTTCCCTACACGACGCTCTTCCGATCTATGTTGCCATCACAACTCATTGGGTTTCTG |
| Hu_VK_MTPX_2 | CTACACTCTTTCCCTACACGACGCTCTTCCGATCTATGGAARCCCCAGCGCAGCTTCTCTTCC |
| Hu_VK_MTPX_3 | CTACACTCTTTCCCTACACGACGCTCTTCCGATCTATGAGGCTCCCTGCTCAGCTCTTGGGGCT |
| Hu_VK_MTPX_4 | CTACACTCTTTCCCTACACGACGCTCTTCCGATCTATGAGGCTCCCTGCTCAGCTCCTGGGGCT |
| Hu_VK_MTPX_5 | CTACACTCTTTCCCTACACGACGCTCTTCCGATCTATGGACATGAGGGTCCCTGCTCAGC |
| Hu_VK_MTPX_6 | CTACACTCTTTCCCTACACGACGCTCTTCCGATCTATGGACATGAGRGTCCTCGCTCAGC |
| Hu_VK_MTPX_7 | CTACACTCTTTCCCTACACGACGCTCTTCCGATCTATGGAAGCCCCAGCACAGCTTCTTCTTCC |
| Hu_VK_MTPX_8 | CTACACTCTTTCCCTACACGACGCTCTTCCGATCTATGAGGCTCCTTGCTCAGCTTCTGGGGCT |

| | |
|---|---|
| Hu_VK_MTPX_9 | CTACACTCTTTCCCTACACGACGCTCTTCCGATCTATGGAAGCCCCAGCTCAGCTTCTCTTCC |
| Hu_VK_MTPX_10 | CTACACTCTTTCCCTACACGACGCTCTTCCGATCTATGGACATGAGGGTCCCCGCTCAGC |
| Hu_VK_MTPX_11 | CTACACTCTTTCCCTACACGACGCTCTTCCGATCTATGGGGTCCCAGGTTCACCTCCTCAG |
| Hu_VK_MTPX_12 | CTACACTCTTTCCCTACACGACGCTCTTCCGATCTATGGTGTTGCAGACCCAGGTCTTCATTTC |
| Hu_VK_MTPX_13 | CTACACTCTTTCCCTACACGACGCTCTTCCGATCTATGGACATGAGGGTGCCCGCTCAGC |
| Hu_VK_MTPX_14 | CTCTTTCCCTACACGACGCTCTTCCGATCTCAGGAAGATGTYGCCATCACAACTCATTGG |
| Hu_VK_MTPX_15 | CACTCTTTCCCTACACGACGCTCTTCCGATCTCTCRCAATGAGGCTCCCTGCTCAGCTC |
| Hu_VK_MTPX_16 | CACTCTTTCCCTACACGACGCTCTTCCGATCTCCTGCTCAGCTCYTGGGGCTGCTAATGC |
| Hu_VK_MTPX_17 | CACTCTTTCCCTACACGACGCTCTTCCGATCTATGGACATGAGGGTGCCCGCTCAGCGCC |
| Hu_VK_MTPX_18 | CACTCTTTCCCTACACGACGCTCTTCCGATCTATGGACATGAGGGTSCCYGCTCAGCKCC |
| Hu_VK_MTPX_19 | CACTCTTTCCCTACACGACGCTCTTCCGATCTGCTCCTGGGGCTGCTAATGCTCTGG |
| Hu_VK_MTPX_20 | CACTCTTTCCCTACACGACGCTCTTCCGATCTGGGGCTCCTGCTGCTCTGGCTCC |
| Hu_VK_MTPX_21 | CACTCTTTCCCTACACGACGCTCTTCCGATCTGGACATGAGGGTCCCCGCTCAGCTCC |
| Hu_VL_MTPX_1 | CTACACTCTTTCCCTACACGACGCTCTTCCGATCTATGGCCTGGGCTCCACTACTTCTCACCCTC C |
| Hu_VL_MTPX_2 | CTACACTCTTTCCCTACACGACGCTCTTCCGATCTATGGCCTGGTCCCCTCTCTTCCTCACCCT |
| Hu_VL_MTPX_3 | CTACACTCTTTCCCTACACGACGCTCTTCCGATCTATGGCCTGGGCTCTGCTCCTCCTCACCCT |
| Hu_VL_MTPX_4 | CTACACTCTTTCCCTACACGACGCTCTTCCGATCTATGGCCTGGAYCCCTCTCCTGCTCCCCCTC |
| Hu_VL_MTPX_5 | CTACACTCTTTCCCTACACGACGCTCTTCCGATCTATGGCCTGGGCTCTGCTGCTCCTCACTCT |
| Hu_VL_MTPX_6 | CTACACTCTTTCCCTACACGACGCTCTTCCGATCTATGGCATGGATCCCTCTCTTCCTCGGCGTC |
| Hu_VL_MTPX_7 | CTACACTCTTTCCCTACACGACGCTCTTCCGATCTATGGCATGGGCCACACTCCTGCTCCCACTC |
| Hu_VL_MTPX_8 | CTACACTCTTTCCCTACACGACGCTCTTCCGATCTATGGCCTGGGTCTCCTTCTACCTACTGCCCT |
| Hu_VL_MTPX_9 | CTACACTCTTTCCCTACACGACGCTCTTCCGATCTATGGCCTGGACTCCTCTTCTTCTCTTGCTCC T |
| Hu_VL_MTPX_10 | CTACACTCTTTCCCTACACGACGCTCTTCCGATCTATGGCCTGGACTCCTCTCCTCCTCCTGYTC C |
| Hu_VL_MTPX_11 | CTACACTCTTTCCCTACACGACGCTCTTCCGATCTATGAGTGTCCCCACCATGGCCTGGATGATG C |
| Hu_VL_MTPX_12 | CTACACTCTTTCCCTACACGACGCTCTTCCGATCTATGGCCTGGGCTCCTCTGCTCCTCACCCTC C |
| Hu_VL_MTPX_13 | CTACACTCTTTCCCTACACGACGCTCTTCCGATCTATGRCCDGCTTCCCTCTCCTCCTCACCCT |
| Hu_VL_MTPX_14 | CTACACTCTTTCCCTACACGACGCTCTTCCGATCTATGGCCTGGACCCCACTCCTCCTCCTCTTC C |
| Hu_VL_MTPX_15 | CTACACTCTTTCCCTACACGACGCTCTTCCGATCTATGGCCTGGGCTCTGCTSCTCCTCASCCT |
| Hu_VL_MTPX_16 | CTACACTCTTTCCCTACACGACGCTCTTCCGATCTATGGCCTGGATCCCTCTACTTCTCCCCCTC |
| Hu_VL_MTPX_17 | CTACACTCTTTCCCTACACGACGCTCTTCCGATCTATGGCCTGGACCSCTCTCCTCCTCRGCCTC |
| Hu_VL_MTPX_18 | CTACACTCTTTCCCTACACGACGCTCTTCCGATCTATGGCCTGGACTCTTCTCCTTCTCGTGCTCC |
| Hu_VL_MTPX_19 | CTACACTCTTTCCCTACACGACGCTCTTCCGATCTATGGCCTGGTCTCCTCTCCTCCTCACTCT |
| Hu_VL_MTPX_20 | CTACACTCTTTCCCTACACGACGCTCTTCCGATCTATGCCCTGGGCTCTGCTCCTCCTGACCCT |
| Hu_VL_MTPX_21 | CTACACTCTTTCCCTACACGACGCTCTTCCGATCTATGGCCTGGACCCCTCTCTGGCTCACTCTC |
| Hu_VL_MTPX_22 | CTACACTCTTTCCCTACACGACGCTCTTCCGATCTATGGCCTGGACCGCTCTCCTTCTGAGCCTC |
| Hu_VL_MTPX_23 | CTACACTCTTTCCCTACACGACGCTCTTCCGATCTATGGCTTGGACCCCACTCCTCTTCCTCACC |
| Hu_VL_MTPX_24 | CTACACTCTTTCCCTACACGACGCTCTTCCGATCTATGGCCTGGACTCCTCTCTTTCTGTTCCTCC |

| | |
|---|---|
| Hu_VL_MTPX_25 | CACTCTTTCCCTACACGACGCTCTTCCGATCTATGGCCTGGACTCTTCTCCTTCTCGTG |
| Hu_VL_MTPX_26 | CACTCTTTCCCTACACGACGCTCTTCCGATCTATGGCCTGGACTCCTCTYCTYCTCYTG |
| Hu_VL_MTPX_27 | CACTCTTTCCCTACACGACGCTCTTCCGATCTATGGCCTGGACCCCACTCCTCCTC |
| Hu_VL_MTPX_28 | CACTCTTTCCCTACACGACGCTCTTCCGATCTATGGCCTGGGTCTCCTTCTACCTACTGC |
| Hu_VL_MTPX_29 | CACTCTTTCCCTACACGACGCTCTTCCGATCTGCAGCATCGGAGGTGCCTCAGCCATG |
| Hu_VL_MTPX_30 | CACTCTTTCCCTACACGACGCTCTTCCGATCTGGCAGAACTCTGGGTGTCTCACCATG |
| Hu_VL_MTPX_31 | CACTCTTTCCCTACACGACGCTCTTCCGATCTGCAGCACTGGTGGTGCCTCAGCCATG |
| Hu_VL_MTPX_32 | CACTCTTTCCCTACACGACGCTCTTCCGATCTGGGCTCTGCTSCTCCTCACYCTCCT |
| Hu_VL_MTPX_33 | CACTCTTTCCCTACACGACGCTCTTCCGATCTGGGCTCTGCTCCTCCTGACCCTC |
| P5_R1 | AATGATACGGCGACCACCGAGATCTACACTCTTTCCCTACACGACGCTCTTCCGATCT |
| P7_R2_I1 | CAAGCAGAAGACGGCATACGAGATCGTGATGTGACTGGAGTTCAGACGTGTGCTCTTCCGATCT |
| P7_R2_I2 | CAAGCAGAAGACGGCATACGAGATACATCGGTGACTGGAGTTCAGACGTGTGCTCTTCCGATCT |
| P7_R2_I3 | CAAGCAGAAGACGGCATACGAGATGCCTAAGTGACTGGAGTTCAGACGTGTGCTCTTCCGATC<br>T |
| P7_R2_I4 | CAAGCAGAAGACGGCATACGAGATTGGTCTGTGACTGGAGTTCAGACGTGTGCTCTTCCGATCT |
| P7_R2_I5 | CAAGCAGAAGACGGCATACGAGATCACTGTGTGACTGGAGTTCAGACGTGTGCTCTTCCGATCT |
| P7_R2_I6 | CAAGCAGAAGACGGCATACGAGATATTGGCGTGACTGGAGTTCAGACGTGTGCTCTTCCGATCT |
| P7_R2_I7 | CAAGCAGAAGACGGCATACGAGATGATCTGGTGACTGGAGTTCAGACGTGTGCTCTTCCGATCT |
| P7_R2_I8 | CAAGCAGAAGACGGCATACGAGATTCAAGTGTGACTGGAGTTCAGACGTGTGCTCTTCCGATC<br>T |
| P7_R2_I9 | CAAGCAGAAGACGGCATACGAGATCTGATCGTGACTGGAGTTCAGACGTGTGCTCTTCCGATCT |
| P7_R2_I10 | CAAGCAGAAGACGGCATACGAGATAAGCTAGTGACTGGAGTTCAGACGTGTGCTCTTCCGATCT |
| P7_R2_I11 | CAAGCAGAAGACGGCATACGAGATGTAGCCGTGACTGGAGTTCAGACGTGTGCTCTTCCGATC<br>T |
| P7_R2_I12 | CAAGCAGAAGACGGCATACGAGATTACAAGGTGACTGGAGTTCAGACGTGTGCTCTTCCGATC<br>T |

Supp. Table 5 | **Antibody pairs from AbDb chosen for the structural variance study described in** Supp. Figure 19**B,C.**

| Pair PDB IDs (ID1–ID2) | Sequence length | Position of single aa difference | Chain type |
|---|---|---|---|
| 2V7N–5GS1 | 118 | 85 | $V_H$ |
| 2V7N–5GRV | 118 | 85 | $V_H$ |
| 5GRV–5GS1 | 118 | 85 | $V_H$ |
| 1FVE–6MH2 | 110 | 54 | $V_L$ |
| 3NAA–3NCJ | 110 | 94 | $V_L$ |
| 1DEE–1HEZ | 110 | 93 | $V_L$ |
| 5I19–6CNR | 110 | 99 | $V_L$ |
| 3X3G–5I1A | 110 | 99 | $V_L$ |
| 3HC0–3HC4 | 110 | 45 | $V_L$ |
| 4HJG–5U5F | 110 | 82 | $V_L$ |

Supp. Table 6 | **Crystal structures of mouse and human paired chain antibody Fv regions from AbDb chosen for MD simulations.**

| **PDB ID** | **Isotype** | **Organism** | **Resolution (Å)** | **Number of residues** | **Number of atoms** |
|---|---|---|---|---|---|
| 1DLF | IgG2κ | Mouse | 1.45 | 233 | 1763 |
| 1MQK | IgG1κ | Mouse | 1.28 | 226 | 1764 |
| 4GXV | IgG1λ | Human | 1.45 | 238 | 1803 |
| 5WCA | IgG1λ | Human | 1.37 | 236 | 1777 |
| 6MEG | IgG1λ | Human | 1.41 | 232 | 1747 |

#### Supplementary Figures

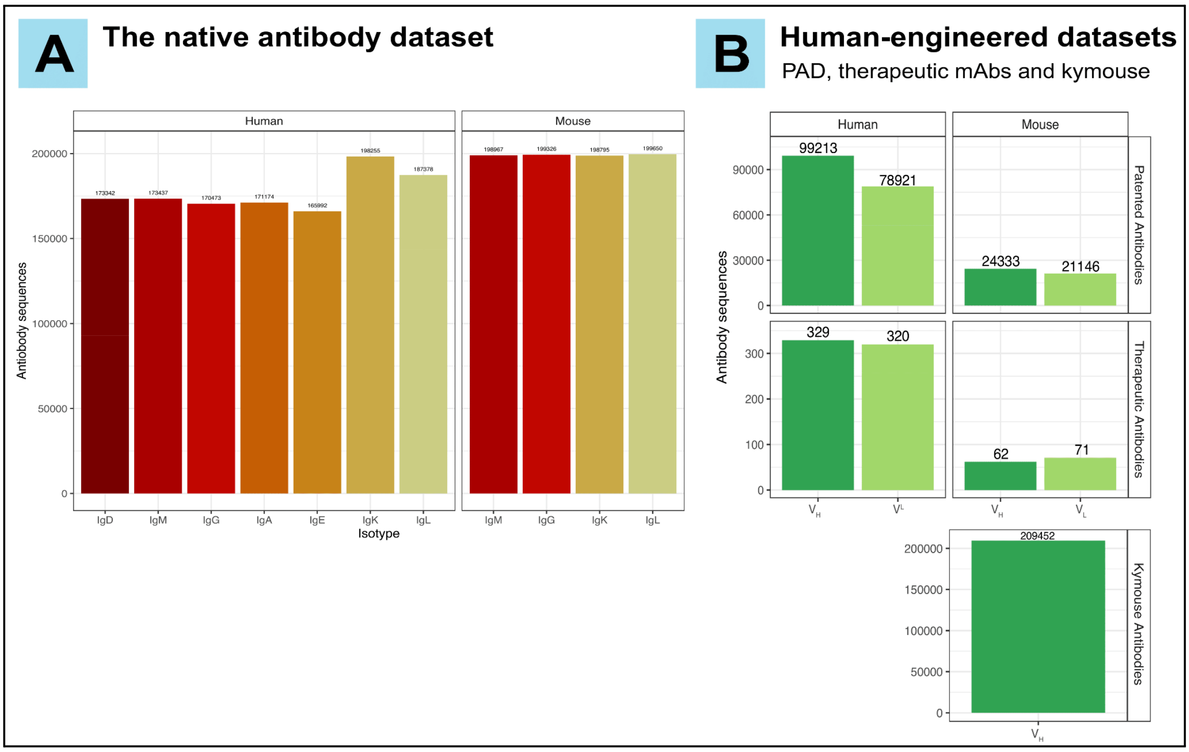

Supp. Figure 1 | **Overview of the native and human-engineered antibody datasets. (A)** The native antibody datasets. We collected a dataset of 2,036,789 unpaired native antibody sequences (variable regions: Fv) from human and mouse repertoires. The majority of them were sourced from Observed Antibody Space (OAS) database (*131*, *172–176*). We also included our own experimentally-generated sequences (298,698 antibodies) within the IgD, IgK and IgL human datasets to provide balanced antibody counts among isotypes (see Methods). **(B)** The human-engineered antibody datasets consist of (i) 223,613 patented antibody sequences obtained from the NaturalAntibody company under a non-commercial agreement (credit: Dr. Konrad Krawczyk). Isotype information was only available for the $V_L$ sequences (IgK or IgL), (ii) 782 therapeutic antibody sequences from TheraSAbDAb (*135*). $V_H$ sequences were all for the IgG isotype and $V_L$ sequences were from both IgK and IgL isotypes, (iii) 209,452 sequences from humanized mice (Kymouse) (*82*). All antibodies in this dataset are $V_H$ sequences of IgM isotype. Relates to Figure 1.

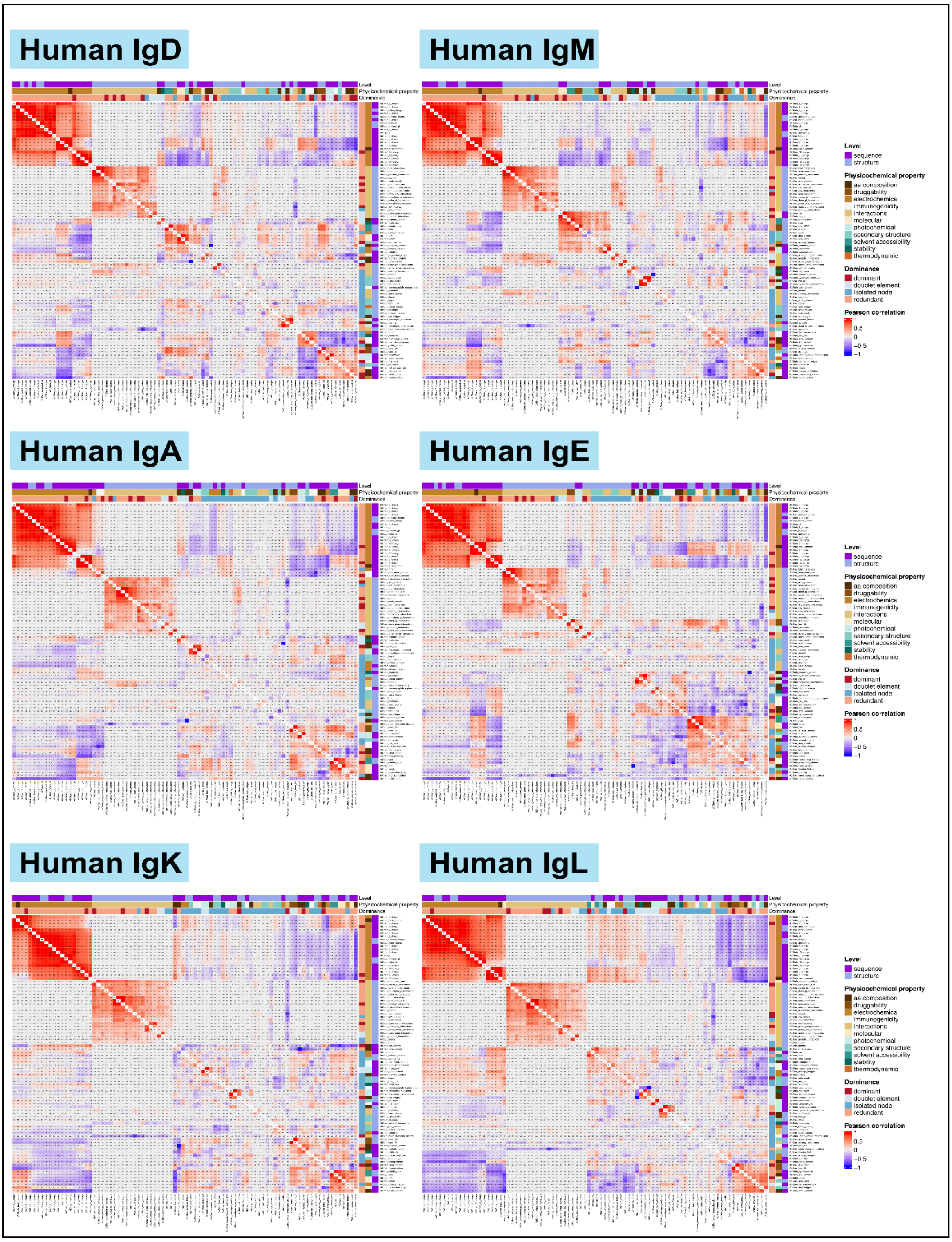

Supp. Figure 2 | **Pairwise developability parameter Pearson correlation for the native human datasets.** Isotype-specific pairwise Pearson correlation coefficient for non-IgG human antibody datasets. Relates to Figure 2.

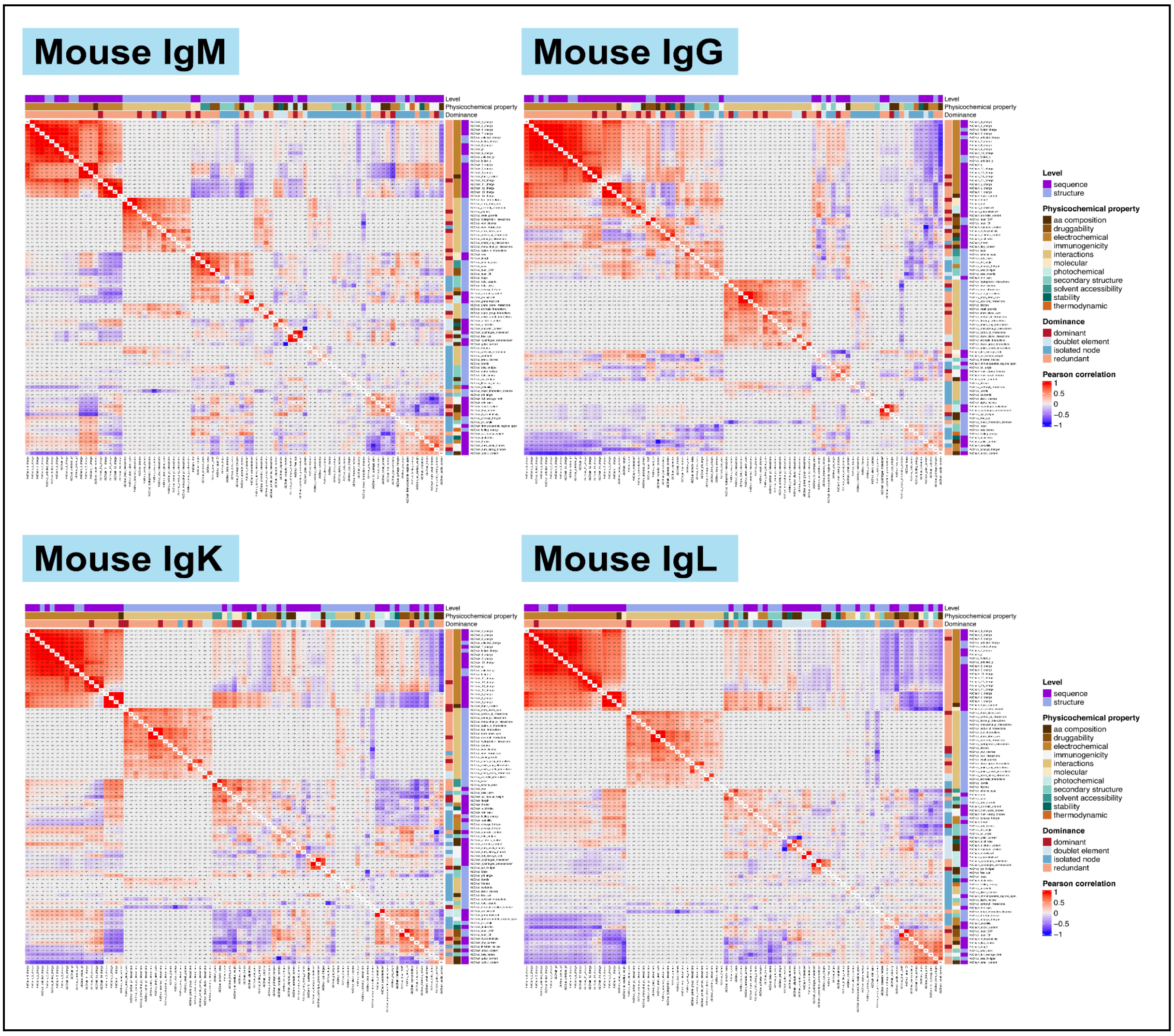

Supp. Figure 3 | **Pairwise developability parameter Pearson correlation for the native murine datasets.** Isotype-specific pairwise Pearson correlation coefficient for murine antibody datasets. Relates to Figure 2.

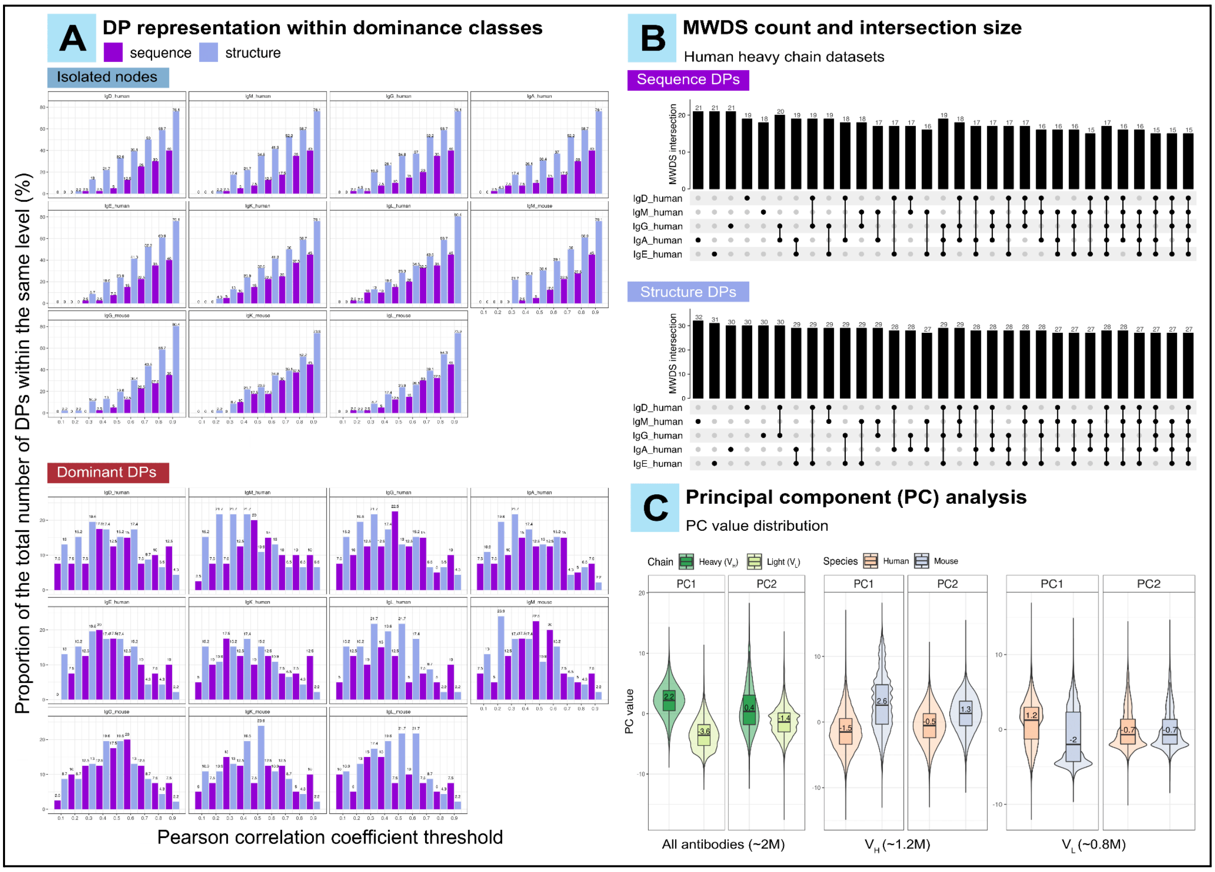

Supp. Figure 4 | **Structure-based developability parameters exhibit lower pairwise association compared to sequence-based parameters.** **(A)** The proportion of DPs within the isolated nodes (top panel) and dominant (bottom panel) classes in relation to an ascending Pearson correlation cut-off (0.1–0.9, step size: 0.1) on sequence and structure levels. **(B)** MWDS intersection size for the human heavy chain datasets (IgD, IgM, IgG, IgA and IgE). The MWDS parameters were identified using the ABC-EDA algorithm (see Methods) at a threshold of absolute Pearson correlation of 0.6. The MWDS intersection study with the murine antibody datasets is shown in Figure 3A. **(C)** The value distribution of the first two principal components (PCs) when analyzing the entire native dataset (left panel ~2M antibodies), the heavy chains (middle panel, ~1.2M antibodies) and the light chains (right panel, ~0.8M antibodies). The numerical values annotated in the figure represent the median of the corresponding metric. Relates to Figure 2 and Figure 3.

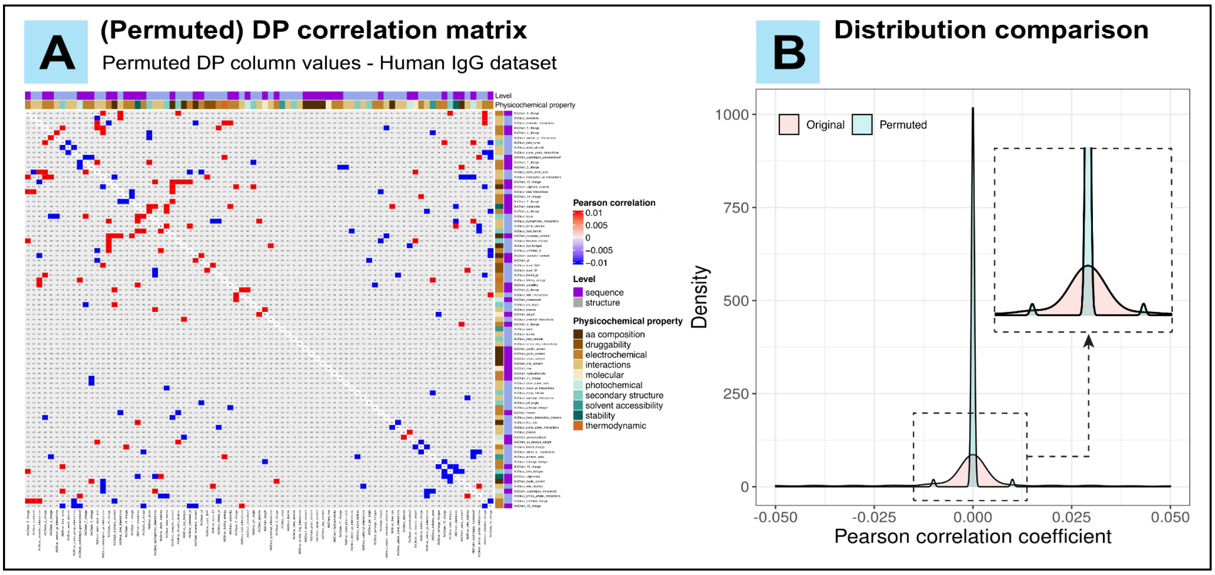

Supp. Figure 5 | **Comparison of pairwise Pearson correlation coefficients calculated on observed and permuted DP column values. (A)** Pairwise DP correlation matrix (equivalent to Figure 2A) where Pearson correlation coefficients were computed on permuted DP column values for the human IgG native dataset, i.e. every antibody from this datasets (170,473) was assigned to a permuted DP value within each DP column. **(B)** A comparison between the distribution of pairwise Pearson correlation coefficients computed on the original and the permuted DP column values for the same dataset (IgG human). A two-sample Kolmogorov-Smirnov test reported a significant difference between the two distributions (p-value < $2.2e^{-16}$). The permutation procedure was repeated 100 times, generating 100 permuted DP correlation matrices. The p-values of the Kolmogorov–Smirnov test were adjusted using the Benjamin-Hochberg method. All p-values reported statistical significance, thus the null hypothesis that the Pearson correlation coefficients computed on original and permuted DP column values are similar in their distributions was rejected. Relates to Figure 2.

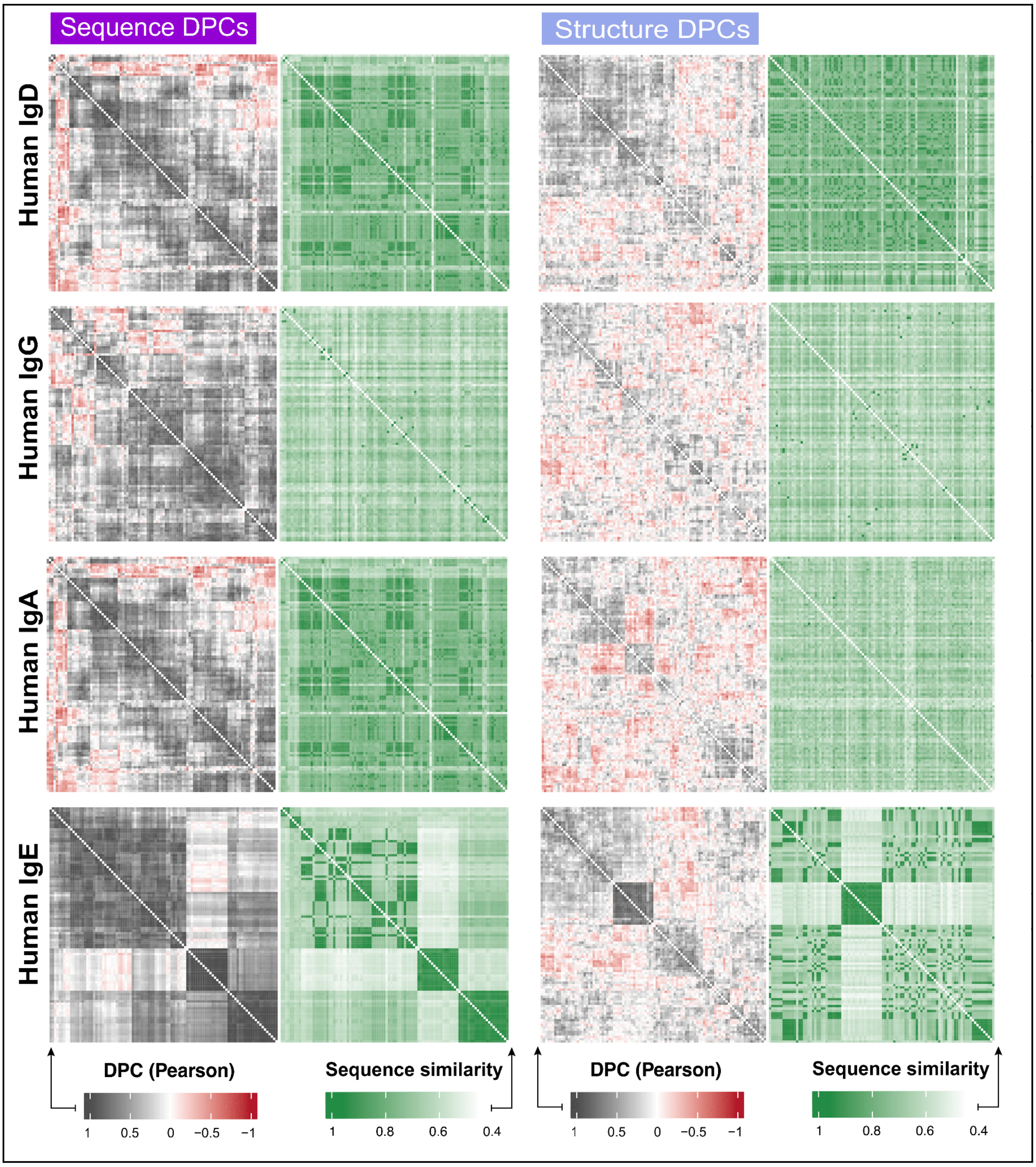

Supp. Figure 6 | **The pairwise developability profile correlation (DPC) alongside the pairwise sequence similarity score for a random sample of 100 natural antibodies from the human heavy-chain datasets (IgD, IgG, IgA and IgE).** Each row and each column represent a single antibody sequence. Rows and columns in the DPC (left) panels were hierarchically clustered. In the sequence similarity (right) panels, rows and columns were ordered in the same order as the corresponding left panel for ease of comparison. Relates to Figure 5.

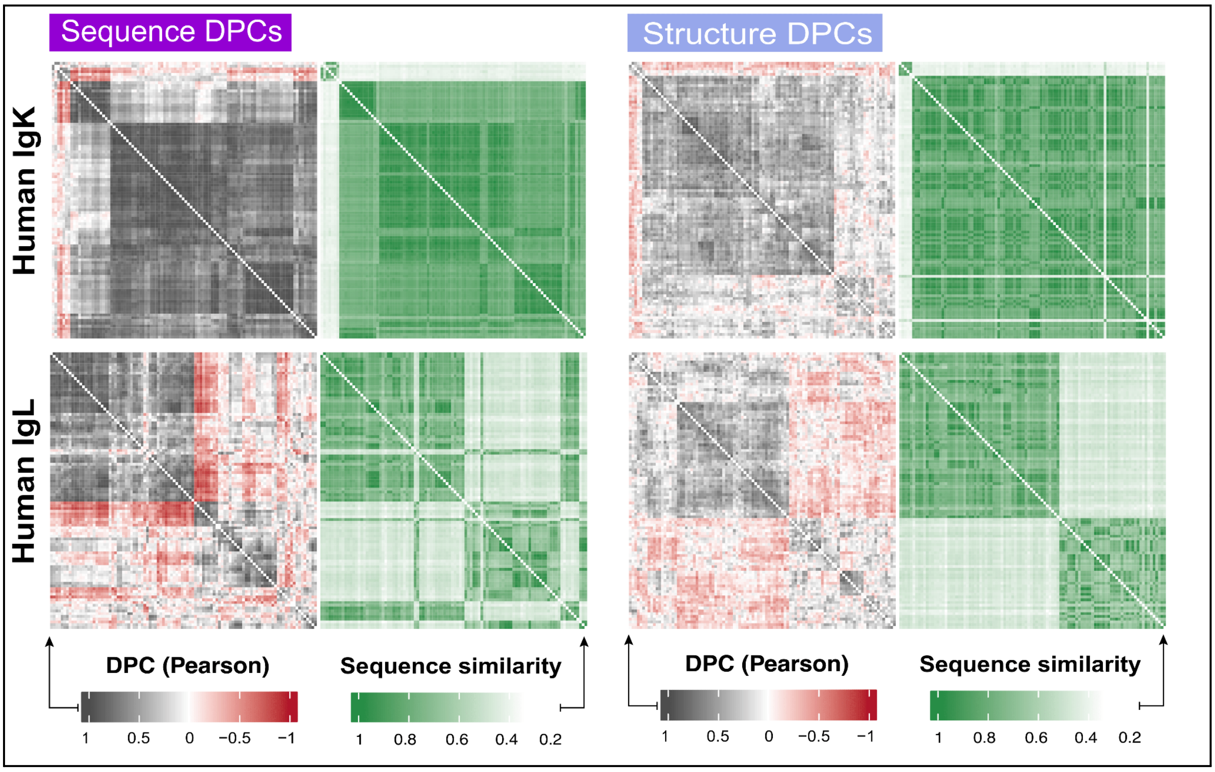

Supp. Figure 7 | **The pairwise developability profile correlation (DPC) alongside the pairwise sequence similarity score for a random sample of 100 natural antibodies from the human light-chain datasets (IgK and IgL).** Each row and each column represent a single antibody sequence. Rows and columns in the DPC (left) panels were hierarchically clustered. In the sequence similarity (right) panels, rows and columns were ordered in the same order of the corresponding left panel for the ease of comparison. Relates to Figure 5.

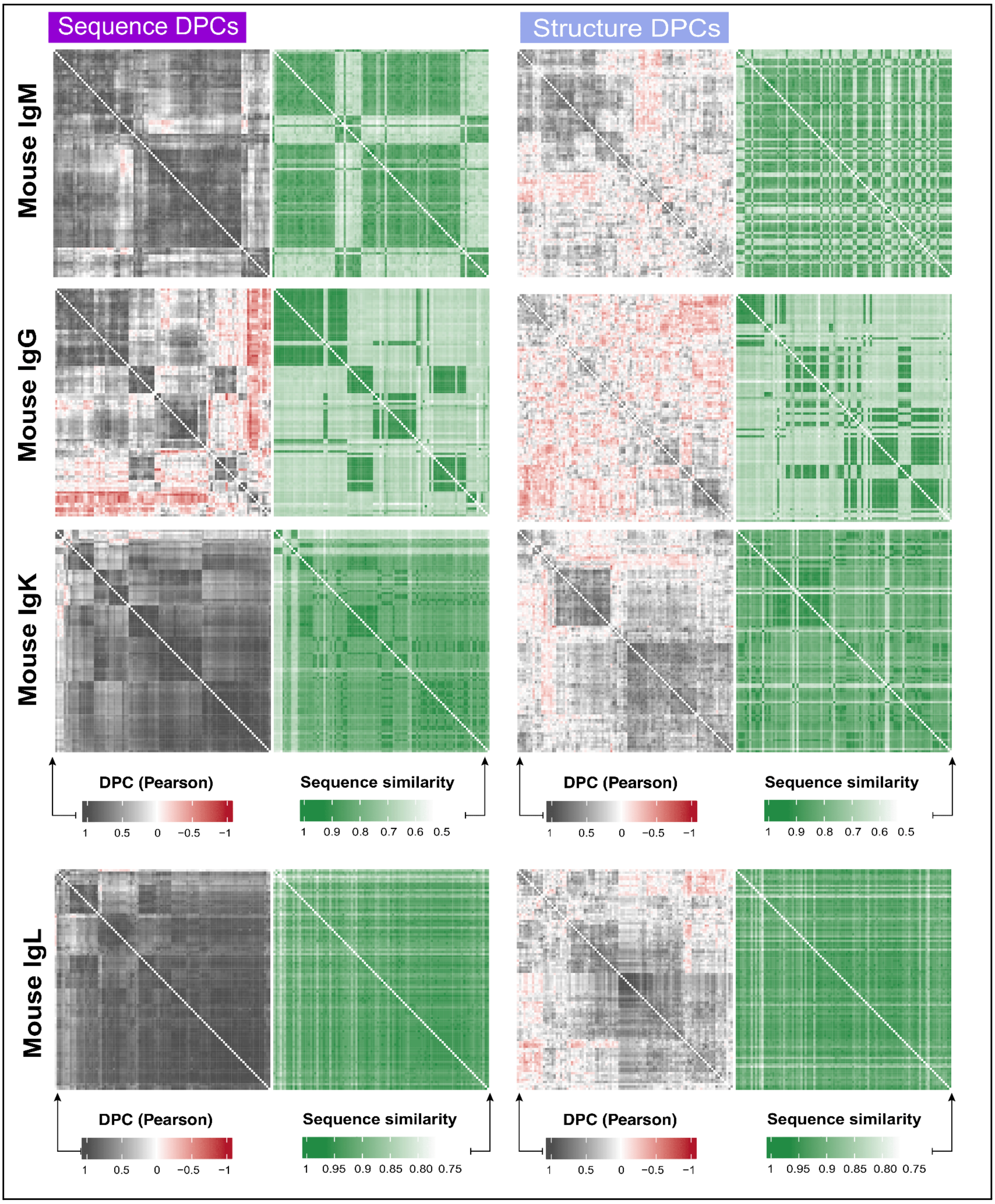

Supp. Figure 8 | **The pairwise developability profile correlation (DPC) alongside the pairwise sequence similarity score for a random sample of 100 natural antibodies from the mouse datasets (IgM, IgG, IgK and IgL).** Each row and each column represent a single antibody sequence. Rows and columns in the DPC (left) panels were hierarchically clustered. In the sequence similarity (right) panels, rows and columns were ordered in the same order of the corresponding left panel for the ease of comparison. Relates to Figure 5.

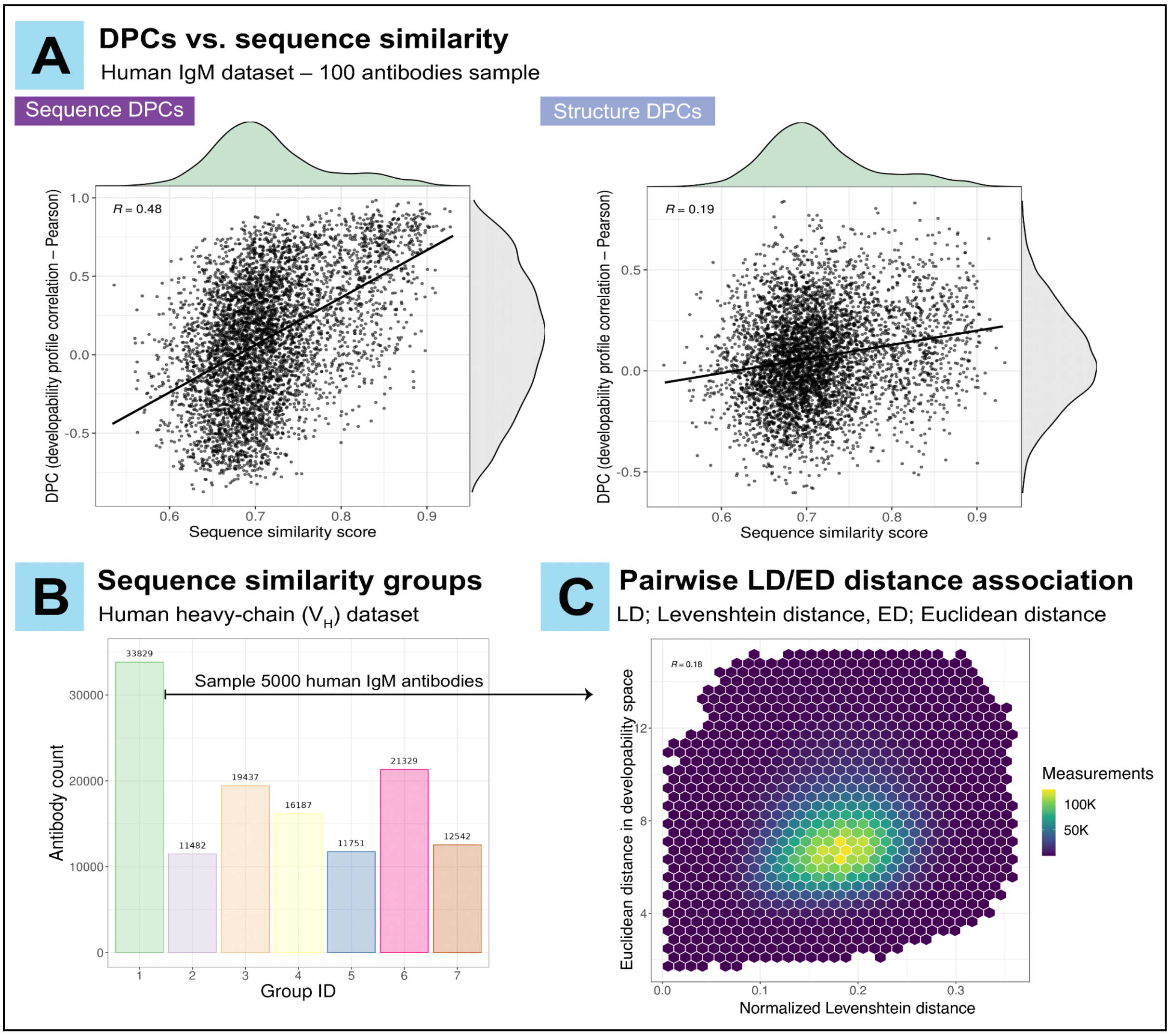

Supp. Figure 9 | **Pairwise antibody sequence similarity does not correlate with pairwise antibody developability similarity. (A)** The distribution of the pairwise developability profile correlation (DPC) and pairwise sequence similarity values for a random sample of 100 natural human IgM antibodies that share the same IGHV gene family annotation. (shown in Figure 5A) This is shown on the sequence level (left panel) and the structure level (right panel). **(B)** The count of antibodies within each sequence similarity group (as shown in Figure 5C). Antibodies that belong to the same group share at least 75% sequence similarity (see Methods). **(C)** The relationship between pairwise (normalized) Levenshtein distance and pairwise Euclidean distance in the developability ($R$) space for 5000 human IgM antibodies that belong to sequence similarity group 1 and share the same IGHV gene family annotation. Out of the total (12,497,500) pairwise measurements, we removed 0.8% where the Euclidean distance was greater than 15 to avoid the effect of outliers on data visualization. Numerical values on the figures in **(A)** and **(C)** represent the Pearson correlation coefficient. Relates to Figure 5.

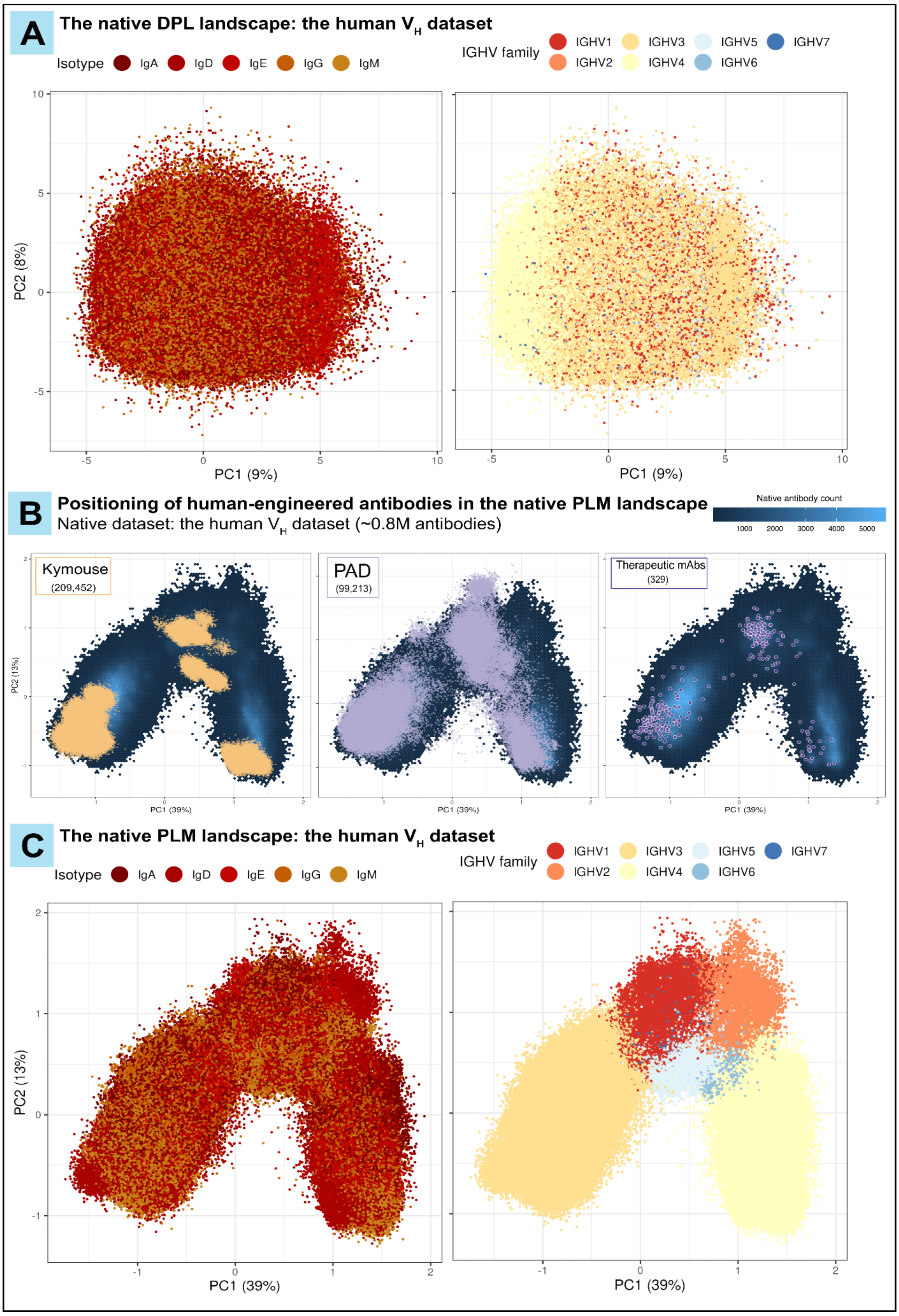

Supp. Figure 10 | **The role of germline gene family annotation and isotype in the localization of $V_H$ antibodies in the native developability and PLM-based sequence landscapes. (A)** The distribution of the native $V_H$ human antibodies (854,418 antibodies) in the developability landscape (DPL-based PCA analysis) shown by antibody isotype (left) and IGHV gene family (right). **(B)** The positioning of the human-aligned human-engineered $V_H$ antibodies (Kymouse; 209,452 , PAD; 99,213 and therapeutic mAbs; 329) in the PLM space of the native human $V_H$ datasets (854,418 antibodies). The hexagonal bins (shown in the back layer) represent the count of native antibodies (scale shown on the top right of the panel), meanwhile, the human-engineered antibodies are represented as data points. **(C)** The distribution of the native $V_H$ human antibodies (854,418 antibodies) in the protein language model landscape (PLM-based PCA analysis) shown by antibody isotype (left) and IGHV gene family (right). Antibody clustering based on germline gene family annotation is observed due to the high sequence similarity (≈80%) among antibodies that belong to the same V-gene family, resulting in PLM-based clustering biases (*177*). Relates to Figure 7.

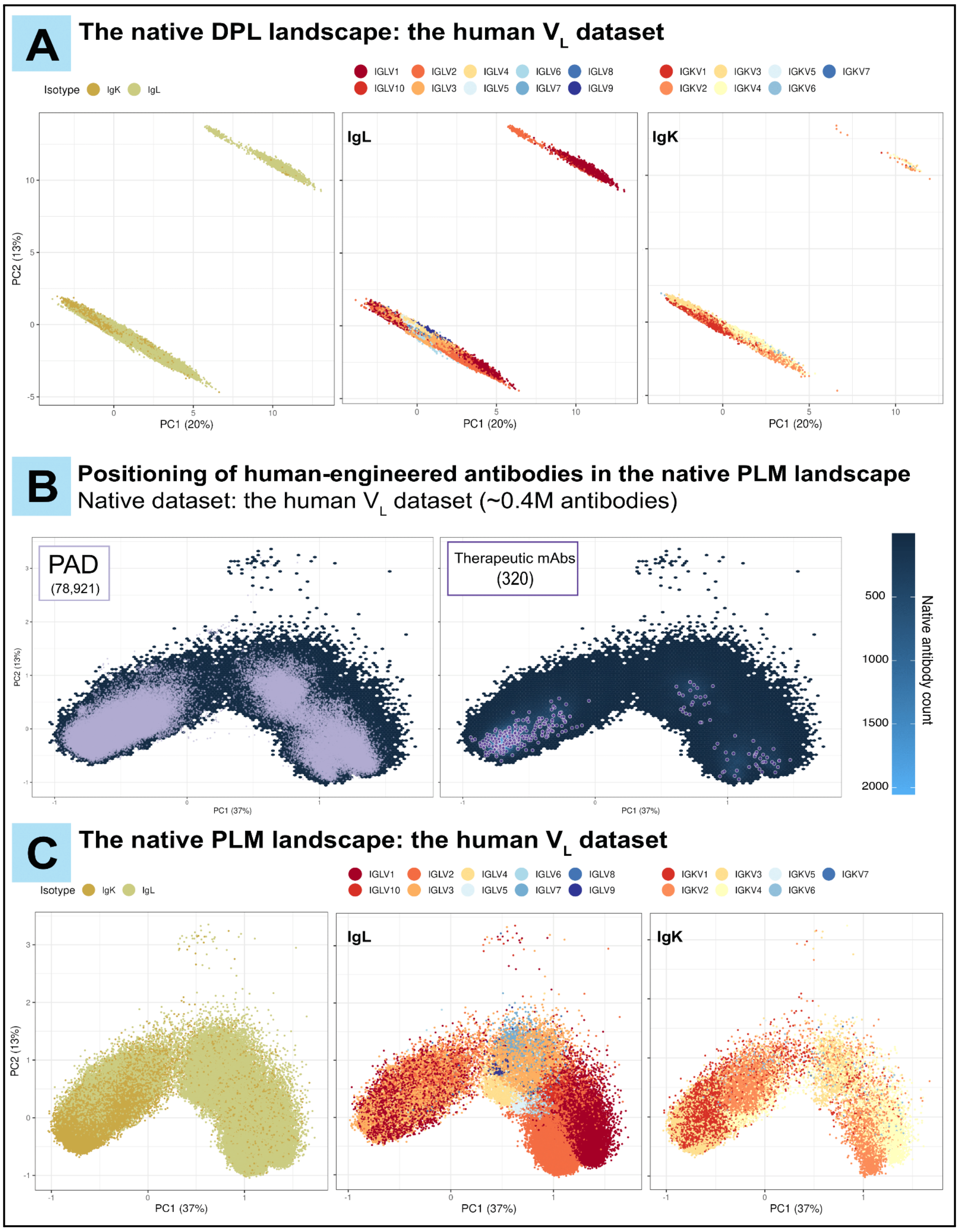

Supp. Figure 11 | **The role of germline gene family annotation and isotype in the localization of $V_L$ antibodies in the native developability and PLM-based sequence landscapes. (A)** The distribution of the native $V_L$ human antibodies (386,173 antibodies) in the developability landscape (DPL-based PCA analysis) shown by antibody isotype (left), IGLV gene family (middle) and IGKV gene family (right). **(B)** The positioning of the human-aligned human-engineered $V_L$ antibodies (PAD; 78,921 and therapeutic mAbs; 320) in the PLM space of the native human $V_L$ dataset (386,173 antibodies). The hexagonal bins (shown in the back layer) represent the count of native antibodies (scale shown on the right of the panel), meanwhile, the human-engineered antibodies are represented as data points. **(C)** The distribution of the native $V_L$ human antibodies (386,173 antibodies) in the protein language model landscape (PLM-based PCA analysis) shown by antibody isotype, IGLV gene family (middle – for IgL antibodies) and IGKV gene family (right – for IgK antibodies). Antibody clustering based on germline gene family annotation is observed due to the high sequence similarity (≈80%) among antibodies that belong to the same V-gene family, resulting in PLM-based clustering biases (*177*). Relates to Figure 7.

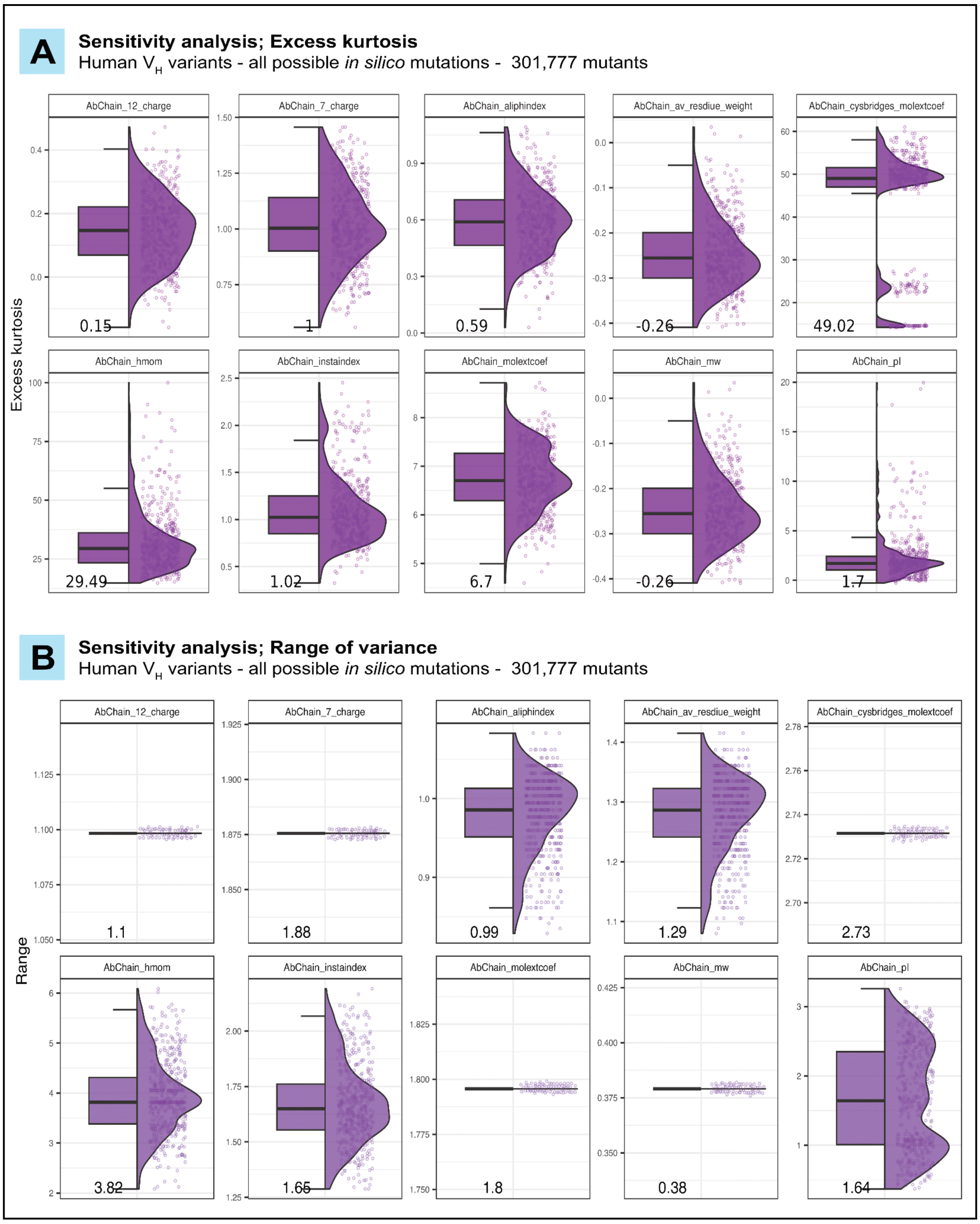

Supp. Figure 12 | **Average and potential sensitivity of sequence-based developability parameters not shown in the main Figure 4.** DP values were computed for all possible single amino acid substituted variants of 500 sampled wildtype human heavy chain sequences (100 sequences sampled per isotype; 301,777 mutants in total). For each DP, the sensitivity was quantified by analyzing the variants' dispersion of DP values compared to their wildtype antibody. Average sensitivity **(A)** was measured by excess kurtosis (low kurtosis = high average sensitivity), while potential sensitivity **(B)** was measured by the range. Relates to Figure 4.

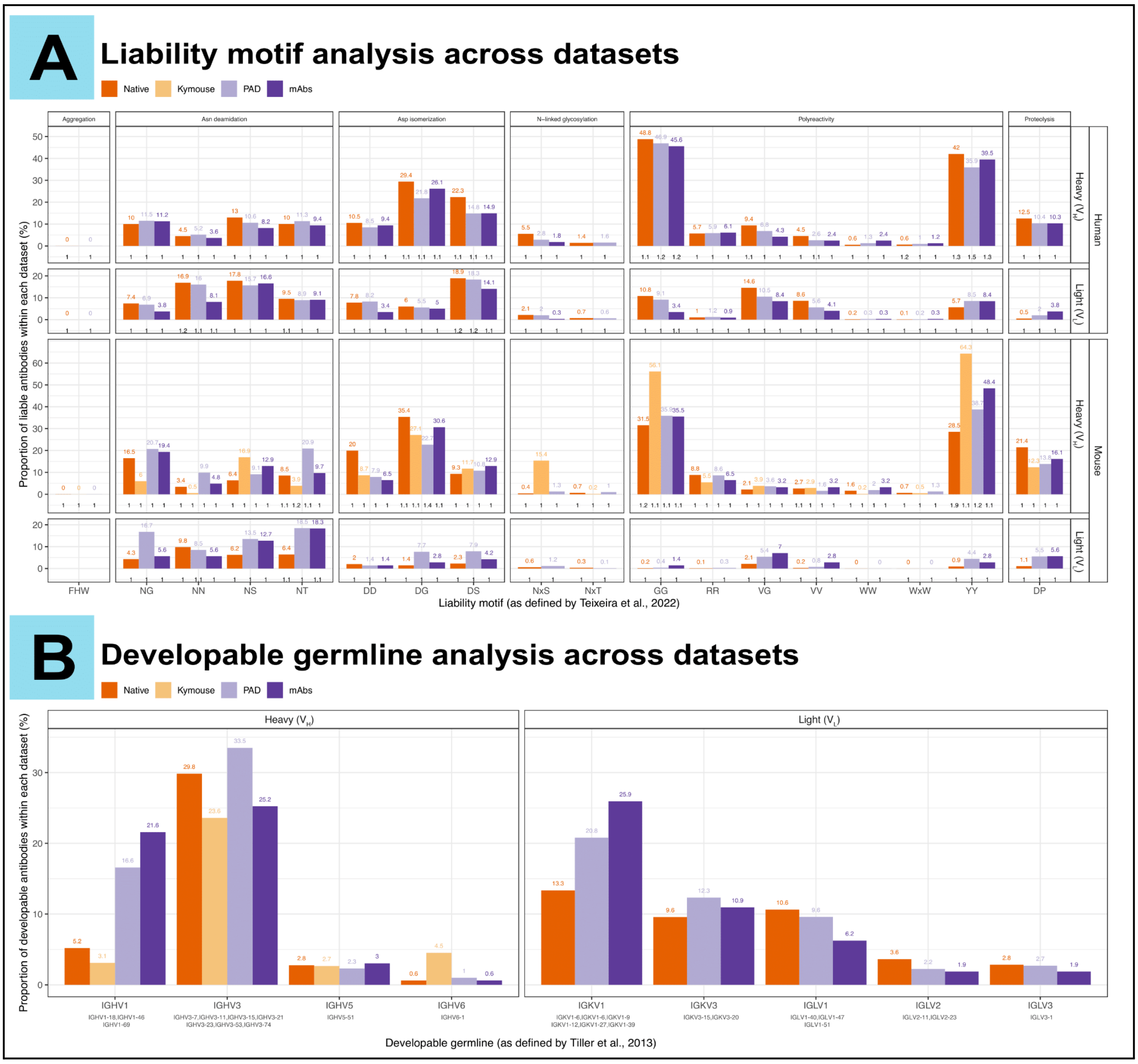

Supp. Figure 13 | **Liability motifs at the native and human-engineered antibody datasets.** Proportional abundance of (**A**) liability motifs and **(B)** developable germlines across the native (human and murine) antibody datasets (Native – Supp. Figure 1A), and the human-engineered datasets (humanized mouse antibodies: Kymouse, patented antibody dataset: PAD, and therapeutic monoclonal antibodies: mAbs). In (A), the x-axis refers to amino acid sequence liability motifs as identified by Teixeira and colleagues (*79*). The figure is segmented into columns (sequence liability type) and rows (antibody species and chain type). The height of the bars and the color-coded text (above them) represent the proportion of antibodies (%) that contain the respective liability motif at least once. The black text on the x-axis refers to the average number of occurrences of liability motifs within liable antibodies (minimum of 1). In (**B**), the x-axis represents the IGHV gene family followed by the developable germline genes (below) within each family as identified by Tiller and colleagues (*80*). The figure is segmented into columns (antibody chain type), and the height of the bars and color-coded text (above them) represent the proportion of antibodies (%) that were annotated with developable germline genes within the respective chain type. Relates to Figure 7.

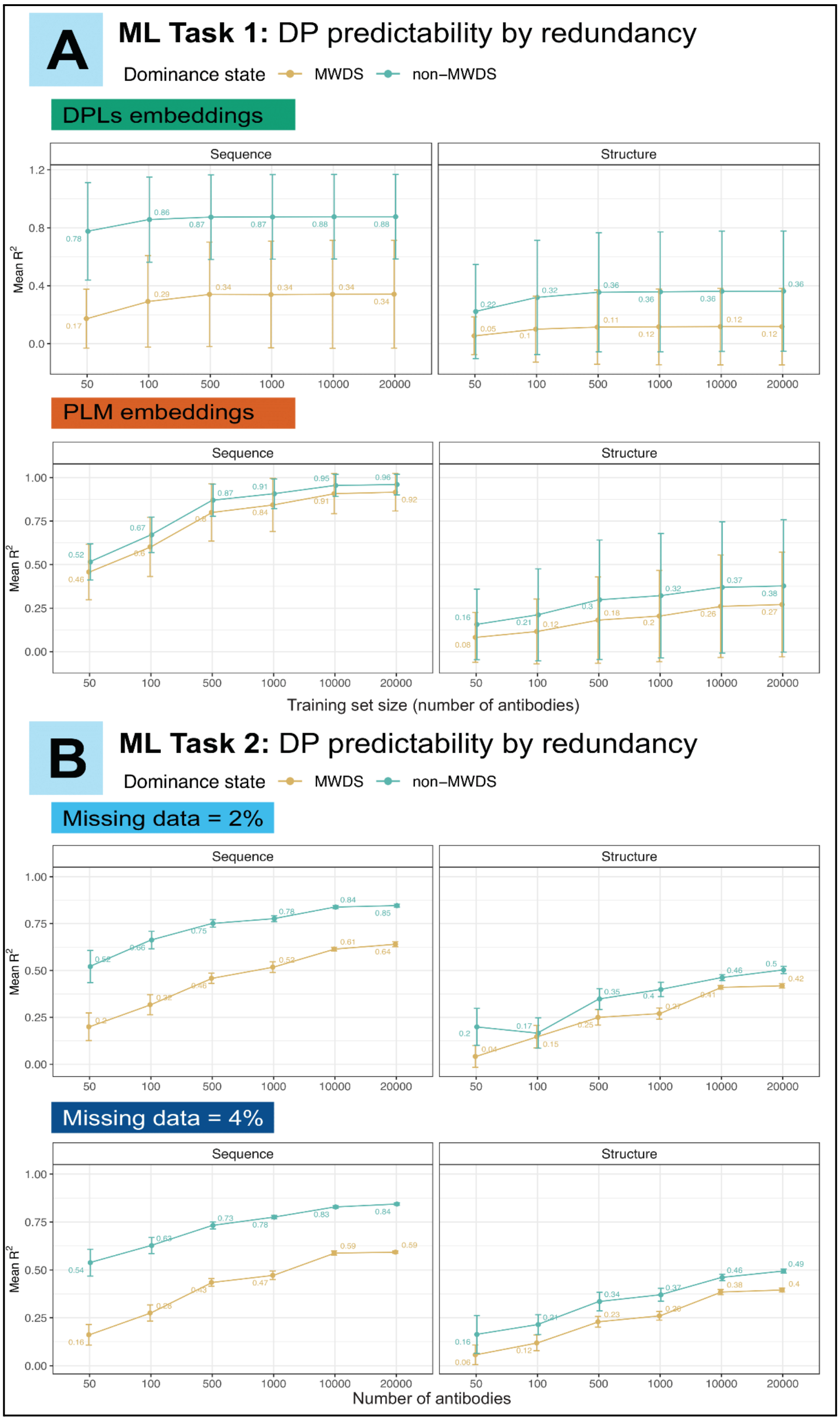

Supp. Figure 14 | **Developability parameters belonging to the MWDS are more challenging to predict than (redundant) non-MWDS ones. (A)** Evaluating the predictive power of single-DP-wise incomplete developability profiles embeddings (top) and PLM-based embedding (bottom) to predict the missing values of (non-redundant) MWDS DPs (yellow lines) and (redundant) non-MWDS DPs using multiple linear regression (MLR – ML Task 1). MWDS developability parameters were identified as previously explained in Figure 6B and listed in Supp. Table 3 (13 sequence-based and 28 structure-based). To evaluate the predictability of non-MWDS parameters, we included an equal number of (randomly-sampled) sequence-based DPs (13) and all the remaining structure DPs that do not belong to the MWDS. X-axis reflects the number of antibodies used for the model embedding (sample size). For each sample size, we repeated the prediction of missing DPs 20 times (20 independent subsamples). Y-axis represents the mean coefficient of determination ($R^2$) for sequence DPs (left facets) and structure DPs (right facets) and error bars represent the standard deviation of $R^2$. **(B)** Evaluating the predictive power of cross-DP-wise incomplete developability profiles embeddings to predict the missing values of (non-redundant) MWDS DPs (yellow lines) and (redundant) non-MWDS DPs multivariate imputation by chained random forests algorithm (MICRF – ML Task 2). MWDS and non-MWDS DPs were identified similarly to (A). Y-axis represents the mean $R^2$ for sequence DPs (left facets) and structure DPs (right facets) when the proportion of the missing data is either 2% (top) or 4% (bottom). Error bars represent the standard deviation of $R^2$. Relates to Figure 6.

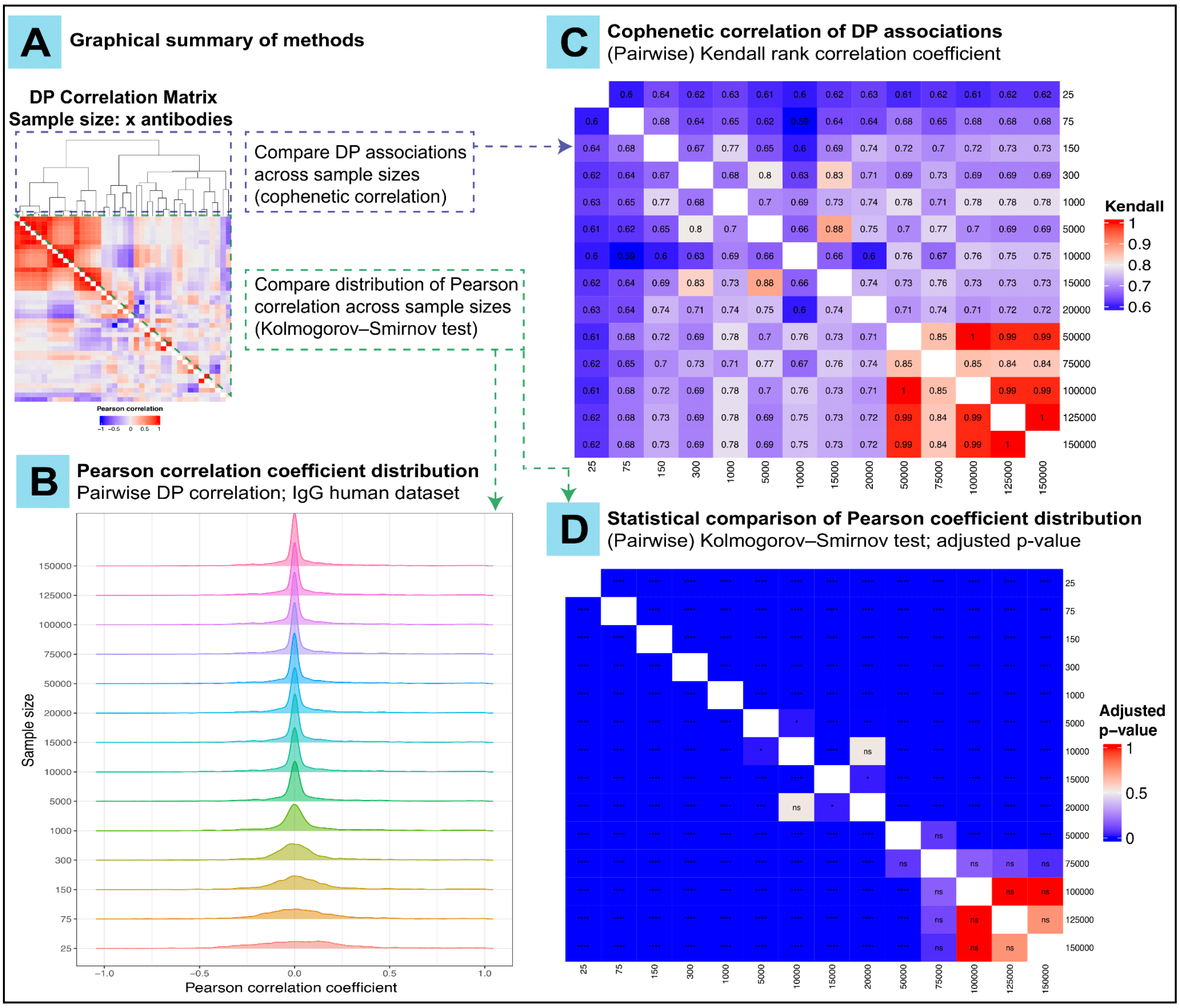

Supp. Figure 15 | **A minimum of 50K antibodies is required to stabilize the pairwise associations among DPs.**
**(A)** Graphical summary of methods followed in this analysis in which we compare DP correlation matrices as a function of sample size. Two pieces of information can be obtained from correlation matrices being (i) DP associations (clustering dendrograms – blue dashed lines) and (ii) Pearson correlation coefficients (green dashed lines). We compare the DP associations (in C) and the distribution of Pearson correlation coefficient values (in B and D) as extracted from pairwise DP correlation matrices for randomly sampled antibodies from the human IgG dataset with increasing antibody sequence sample size (25, 75, 150, 300, 1000, 5000, 10000, 15000, 20000, 50000, 75000, 100000, 125000, 150000 sequences). **(B)** Visual comparison of the distribution of Pearson correlation coefficient values as a factor of increasing antibody sequence sample size. **(C)** Pairwise Kendall correlation coefficient was computed among the hierarchical clustering dendrograms extracted from pairwise DP correlation matrices computed on. At least 50K antibodies are required to sustain the associations (dendrograms) among DPs (Kendall correlation coefficient ≥ 0.84). **(D)** Statistical comparison of the distribution of Pearson correlation coefficient values as a factor of increasing antibody sample size using Kolmogorov-Smirnov test. At least 75K antibodies are required to preserve the distribution of Pearson correlation values among DPs (the distribution difference becomes statistically non-significant from 75K antibodies onwards). ns; not-significant, * $p < 0.05$, ** $p < 0.01$, *** $p < 0.001$ and **** $p < 0.0001$. Relates to Figure 7.

#### Supplementary Methods

##### Experimentally generated native antibody sequences (human IgD, IgK and IgL)

*(1) Human subjects, B-cell isolation and RNA extraction*

Human peripheral blood was obtained from one healthy volunteer. Sample acquisition was approved by the Regional Ethics Committee of South-Eastern Norway (project 6544). Samples were collected in BD Vacutainer® K2 EDTA tubes, and pan B cells were isolated by negative selection using MACSxpress® Whole Blood B Cell Isolation Kit (Miltenyi Biotec). The cells were washed with PBS and remaining erythrocytes were lysed using Red Blood Cell Lysis Solution (Miltenyi Biotec). RNA was extracted using RNeasy Kit (Qiagen). RNA quality and concentration were measured using a Nanodrop spectrophotometer (Thermo Fisher Scientific).

*(2) cDNA synthesis*

200 ng of RNA was used for cDNA synthesis with 1 µl of 100 µM isotype-specific reverse primers, 1 µl of 10mM dNTP Mix (Thermo Fisher Scientific) and nuclease-free water up to 14.5 µl. Mixture was incubated for 5 min at 65°C, briefly placed on ice and centrifuged. Subsequently, 4 µl of 5X RT buffer (Thermo Fisher Scientific), 0.5 µl of RiboLock RNase Inhibitor (Thermo Fisher Scientific), and 1 µl Maxima RT (Thermo Fisher Scientific) were added and cDNA synthesis was performed at 50°C for 30 min with reaction termination at 85°C for 5 min. The obtained cDNA was purified using MinElute PCR Purification Kit (Qiagen) and eluted in 20 µl of EB buffer.

*(3) 5' Multiplexing (MPTX) PCR*

4 µl of cDNA were amplified with 0.5 µl of 100 µM Read2U primer, 1 µl of chain-specific 5′ forward leader primer mix (Supp. Table 4), 10 µl of KAPA HiFi HotStart ReadyMix (Roche Molecular Systems), and 4.5 µl of nuclease-free water at the following conditions: 96°C – 5 min; 25 cycles of 95°C – 20 sec, 68°C – 20 sec, 72°C – 20 sec; 72°C – 5 min; 4°C – hold. Amplified product was run in a 1.2% agarose gel in TBE buffer. Bands corresponding to the amplified regions of interest (approx. 480 bp) were cut and purified using QIAquick Gel Extraction Kit (Qiagen) with elution in 20 µl of EB buffer.

*(4) NGS library generation*

For indexing PCR, 10 ng of the 5' MTPX product were mixed with 0.5 µl of 100 µM P5_R1 forward indexing primer, 0.5 µl of 100 µM P7_R2 reverse indexing primer containing Illumina index sequence, 12 µl of KAPA HiFi HotStart ReadyMix, and nuclease-free water up to 24 µl and reaction was performed at the following conditions: 96°C – 5 min; 10 cycles of 95°C – 30 sec, 68°C – 30 sec, 72°C – 30 sec; 72°C – 10 min; 4°C – hold. The resulting libraries were bead-purified with AMPure XP (Beckman Coulter) using a 1:1 beads ratio. The molarity of the libraries was determined using Qubit™ 4 Fluorometer. The final libraries were inspected with BioAnalyzer (average product length 550 bp) and sequenced on the Illumina MiSeq platform (V3 chemistry 300x2 bp). The raw sequencing data is available on the Sequence Read Archive (BioProject number PRJNA1043047).

##### Validation datasets

*(1) The crystal structures dataset*

We obtained 859 crystal structures of paired-chain antibodies (Fv regions) from the antibody structure database (AbDb) (*178*). We extracted the amino acid sequences for both (heavy and light) chains and utilized them to further predict the structure of paired and unpaired antibody chains *in silico*. We used several tools to benchmark our analysis including ABodyBuilder (*36*), ABodyBuilder2 (*37*), IgFold (*38*), AlphaFold2 (*179*), and AlphaFold-multimer (*180*). Of note, the alignment process and regional sequence definition (CDRs and FRs) was performed similarly to the processing steps mentioned for the therapeutic antibodies dataset (see Methods). We finally computed the values of structure-based DPs on these structures to perform the rigid model analysis described in Supp. Figure 17.

*(2) The in silico mutated antibody dataset at CDRs*

We subsampled 10 WT antibodies per isotype, except 9 for IgD, and all their corresponding CDR mutants (a total of 30,015 antibody sequences) from the *in silico* mutated dataset (see Methods), and predicted their structures (including WT antibodies) using IgFold version 0.1.0 (*38*). We used IgFold for this task as template search-based tools (including ABB) tend to utilize an identical template for antibodies that are within the distance of a single amino acid substitution from their wildtype antibody (*181*). We used this dataset to conduct the structural variance study described in Supp. Figure 19A.

*(3) The AbDb antibody pairs*

To study the structural variance of antibody mutants with a single aa difference, we sourced 10 pairs of antibodies (Supp. Table 5) from the public AbDb database (*178*). We ensured that each pair of antibodies (1) has the same sequence length and (2) the different (single) amino acid is located in the loops. This dataset was used in the structural variance study described in Supp. Figure 19B,C.

#### Molecular dynamics simulations

We selected five paired-chain ($V_H$ and $V_L$) antibody structures from the crystal structures dataset with the best resolutions (Supp. Table 6) to perform classical MD simulations. Two antibody structures (4TRP and 5WCA) contained missing atoms which were corrected with the MODELLER's function "complete_pdb" (*182*). Hydrogens were added to initial antibody structures using Reduce ((*147*)). All MD steps were performed in Gromacs v.2022.4 (*183*). We used AMBER99SB-ILDN (*184*) force field and TIP3P (*185*) water model for MD system preparation (*186*). The simulation box was defined as a cube centered around the antibody placed at the 1 nm distance between the solute and the box. All systems contained both Na and Cl ions at 0.1 mol/liter salt concentration. We minimized the energy of each initial system with the steepest descent algorithm (*187*) for 20000 steps. For the equilibration (*188*), we first considered constant volume simulations (NVT) at 300 K for 100 ps and followed up by constant pressure simulations (NPT) at 1 bar for another 10 ns. During these equilibration simulations (NVT and NPT), where applicable, we used the Parrinello-Rahman barostat (*189*) and V-rescale thermostat (*190*) with velocity rescaling using 0.1 ps and 0.1 ps time constants, respectively. During all the simulations, we constrained the length of all bonds using the linear constraint solver (LINCS) algorithm (*191*) and kept the water molecules rigid via the SETTLE algorithm (*192*). We used particle mesh Ewald (PME) (*193*) for treating the electrostatic interactions with a real-space cutoff of 1.0 nm (*193*). We simulated a 100 ns independent production run with 2 fs time step for each system, all continued from the last step of the NPT simulations at 300 K and 1 bar using a 2 fs time step. Further details of the parameter specifications can be found in .mdp files. MD simulations were performed starting from crystal structures and ABB-predicted models for the five selected antibodies (Supp. Table 5), followed by the computation of structure-based DPs on the convergent frames (4001 frames per antibody per structure origin). This data was used to perform the molecular dynamics analysis described in Supp. Figure 18.

#### Computation of the overlap index (η)

For the analysis in Supp. Figure 18C, we used the 'overlapping' R package (*194*) to quantify the overlap between the distributions of DP values when measured on the MD-simulated crystal structures and the MD-simulated ABB-predicted structures.

### Supplemental File

#### Verification of strong correlation between DP values measured on single and paired-chain antibody structures

We found that the values of structure-based DPs measured on a subset of 859 paired-chain antibodies (see Methods) strongly correlated with the corresponding DP values measured on each of the unpaired single chains ($V_H$ and $V_L$) separately (median Pearson correlation 0.84–0.90, Supp. Figure 16) thus supporting the notion that some DP work performed on single-chains data translates to paired-chain data.

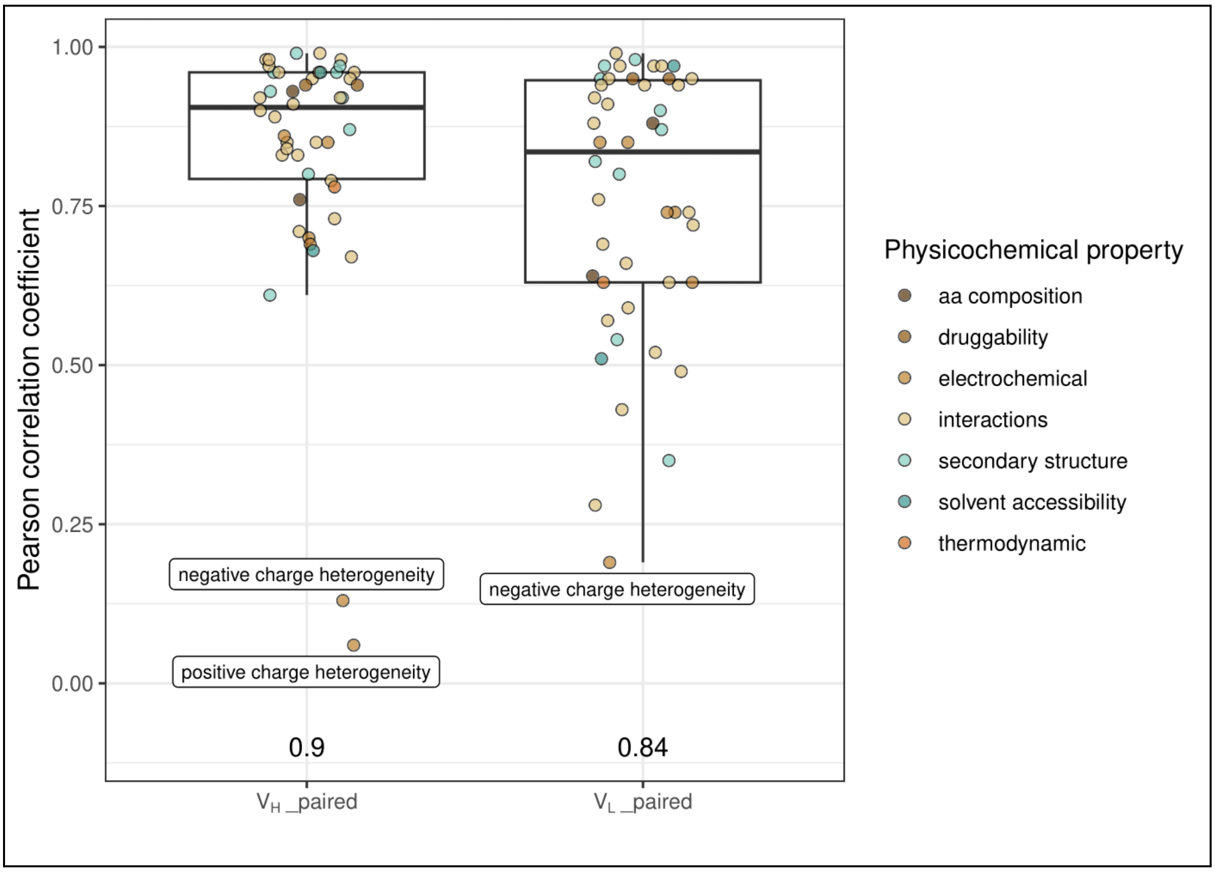

Supp. Figure 16 | **Pearson correlation of 46 structure DPs measured on single-chain ($V_H$: "heavy", $V_L$: "light") compared to corresponding paired-chain structure DP values.** DP values were computed on the crystal structure dataset (859 antibodies – see Methods). The figure legend refers to the classification of DPs by physicochemical property (as detailed in Supp. Table 1). Numbers on the x-axis represent the median Pearson correlation coefficient. The positive and negative charge heterogeneity DPs show the lowest correlation across all structures (Pearson correlation <0.2, labeled data points). Relates to Figure 2.

#### Analysis of dependence of structure-based DPs on computational antibody structure prediction method

We set out to verify to what extent structure DPs are stable across computational antibody structure prediction methods. We performed these analyses both for rigid (Supp. Figure 17) and dynamic antibody models (i.e., molecular dynamics, Supp. Figure 18) starting from 859 reference (crystal) antibody structures obtained from AbDb (crystal structures dataset, see Methods) (*178*).

*(1) Rigid model analysis:*

The rigid structure prediction tools included in this analysis were ABodyBuilder (*36*), ABodyBuilder2 (*37*), IgFold (*55*), AlphaFold2 (*179*), and AlphaFold-multimer (*180*). For this analysis, we measured the structural variance (RMSD – Supp. Figure 17B) and the correlation of DP values (Pearson correlation – Supp. Figure 17A) between crystal structures and their predicted models. We conducted this analysis on $V_H$-$V_L$ paired structures as well as $V_H$ and $V_L$ (unpaired) structures separately (Supp. Figure 17A,B).

ABB-predicted rigid structures harbored minimal atomic distances from their reference counterparts (median RMSD = 0.34, 0.2 6, 0.27 ordered for paired, $V_H$ and $V_L$ structures – Supp. Figure 17B), stemming from high structural alignment with reference antibodies (example shown for the antibody 1DLF in Supp. Figure 17C). Nevertheless, despite this seemingly high structural similarity, the correlation of the DP values measured on ABB structures with those measured on crystal structures was low (median Pearson correlation coefficient = 0.31, 0.4, 0.31 for paired, $V_H$ and $V_L$ respectively – Supp. Figure 17A). Moreover, we reported similar low correlation coefficients between DP values measured on crystal structures and all predicted structures included in this study (Supp. Figure 17A). These observations were reported despite ABB being a former software release in comparison to the remaining tools included in this study. These results demonstrate the dependence of structure-based antibody developability predictions on the choice of computational structure prediction method. A similar observation was made recently by Jain and colleagues (*31*).

Given the differences in DP values among the structure prediction tool, we asked in the next section whether these differences may be explained by comparing rigid structures from frames belonging to a conformational ensemble (dynamic continuum).

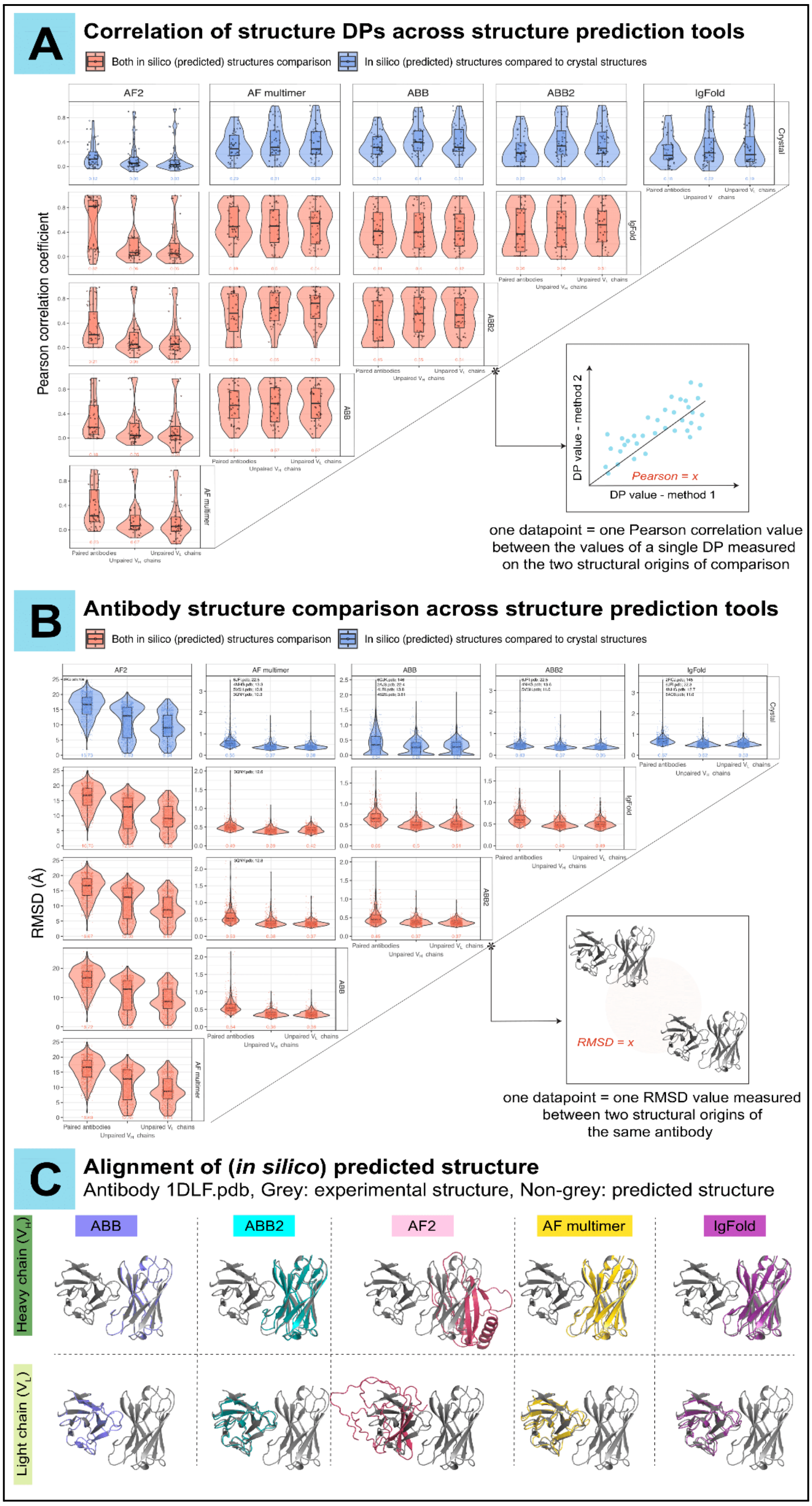

Supp. Figure 17 | **ABodyBuilder-predicted antibody structures exhibit comparable conformational agreement and closer developability resemblance with reference antibody structures compared to several other state-of-the-art structure prediction tools. (A)** Pairwise Pearson correlation coefficients of structural DPs between antibody structure of experimental or in silico origin. The structures of antibodies (paired and unpaired) in the crystal structures dataset (859 structures – see Methods) were predicted using the in silico tools (AF2; AlphaFold2 (*179*), AF multimer; AlphaFold-multimer (*180*), ABB; ABodyBuilder (*36*), ABB2; ABodyBuilder2 (*37*), and IgFold (*38*)). Structure-based DP values were computed on all resulting structures (**46** structure-based DPs, Supp. Table 1). Each data point in a boxplot refers to one Pearson correlation coefficient between the values of a single structural DP measured on the two structural origins of comparison (see inset graphic on the right) for all 859 antibody structures in the crystal datasets. Numbers on the x-axis represent the median of Pearson correlation values within each chain group (paired, unpaired $V_H$, unpaired $V_L$). **(B)** Pairwise root mean square deviation (RMSD) between two antibody structural origins. RMSD values were computed between experimental (crystal) and in silico predicted structures of antibodies from the same dataset (as in A). Each data point in a boxplot represents one RMSD value measured between two structural origins of the same antibody (see inset graphic on the right). Numbers on the x-axis represent the median RMSD within each chain group (paired, unpaired $V_H$, unpaired $V_L$). Values higher or equal to ten times the median RMSD within each chain group were considered outliers, and are listed above the corresponding chain group and antibody origin comparison pair (PDB ID; RMSD). **(C)** Example visualization of pairwise alignment between in silico predicted structures and the reference structure of a randomly selected antibody from the crystal structure dataset (PDB ID: 1DLF). Relates to Figure 2.

##### *(2) Molecular dynamics analysis:*

We investigated whether the rigid ABB-predicted structures represented a mere snapshot of the structural conformations of the corresponding crystal (reference) antibodies when examined with molecular dynamics simulations (MD). To this end, we included five paired-chain antibodies in this analysis, prioritizing those with the highest structural resolution within the crystal structure dataset (PDB IDs: 1DLF, 1MQK, 4GXV, 5WCA, 6MEG – Supp. Table 6). The selection of sample size (five) and antibody structure type (paired-chain) was made considering (i) the computational cost associated with molecular dynamics simulations, and (ii) the strong correlation between single and paired-chain structure-based antibody developability as previously shown (Supp. Figure 16). We simulated the crystal and ABB-predicted structures of these antibodies over 100 ns resulting in 5001 frames/antibody, and confirmed structural convergence for all antibodies (crystal and ABB-predicted) from 20 ns onwards (see Methods – Supp. Figure 18A). Examining the pairwise structural variance (measured with RMSD) among convergent ABB frames (4001 frames/antibody) and convergent reference frames (4001 frames/antibody) revealed limited structural disparities (Supp. Figure 18B). Indeed, ABB frames were minimally distant from their reference counterparts (median pairwise RMSD 1DLF;1.34Å, 1MQK;1.12Å, 4GXV;1.1Å, 5WCA;1.08Å, 6MEG;0.97Å – Supp. Figure 18B). Importantly, the distribution of their structural DP values highly overlapped with those of the crystal (reference) equivalents with a mean overlap index ($\eta$) of 0.73, 0.65, 0.77, 0.85, 0.77 for 1DLF, 1MQK, 4GXV, 5WCA, 6MEG, respectively (Supp. Figure 18C), suggesting that the difference in structure-based DPs as a function of 3D-structure origin (be it experimental or computational) may be a consequence of undersampling the dynamics of the antibody structure space. Our findings are in line with recent MD-driven findings in regards to structure-based DPs (*29*, *33*, *34*). Future research in ML-based MD may enable repertoire-level MD as, currently, performing MD calculations beyond a handful of structures is prohibitively computationally expensive (*195*).

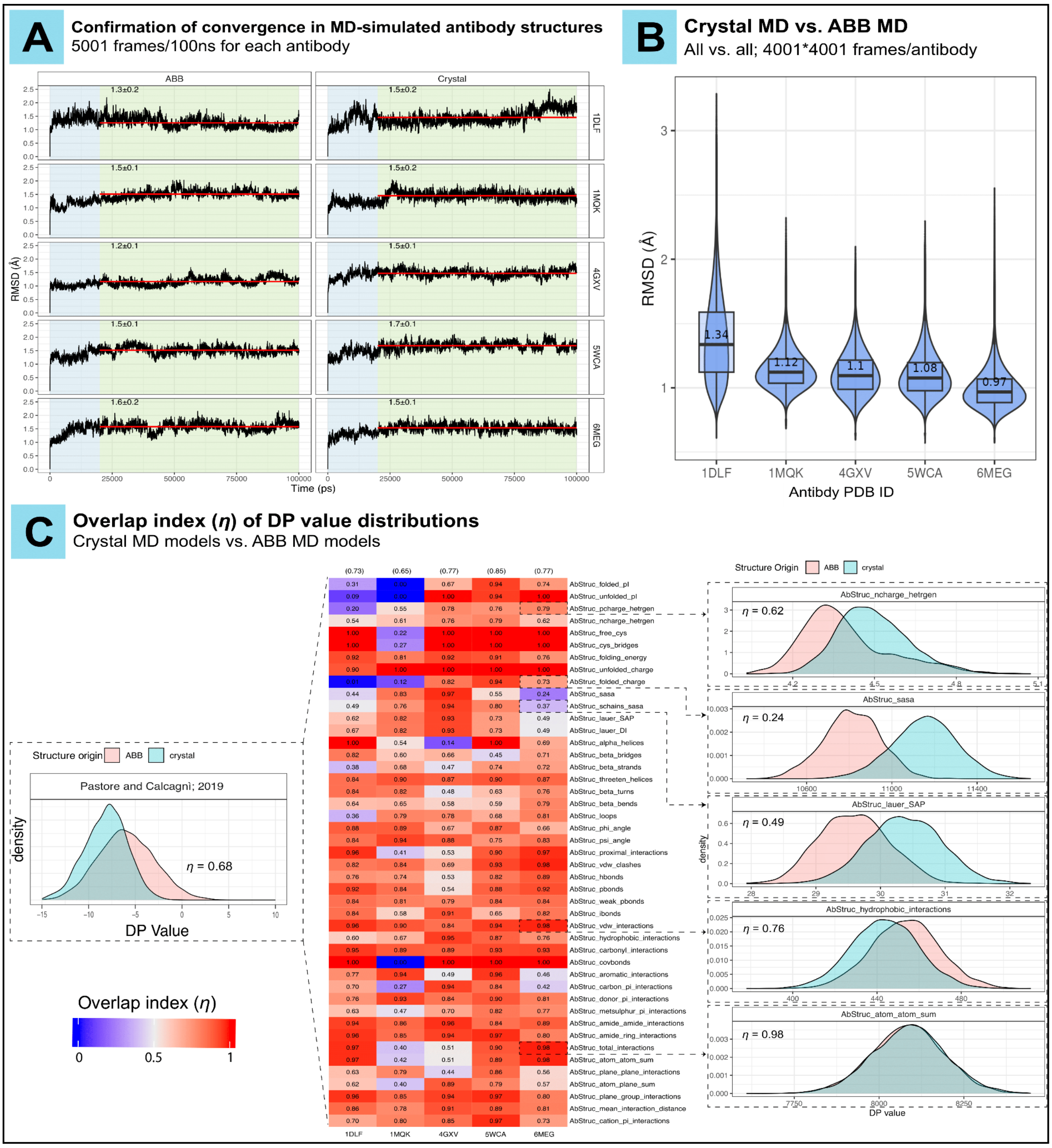

Supp. Figure 18 | **ABodyBuilder(ABB)-predicted structures show minimal conformational variance and overlapping structural developability in molecular dynamic (MD) simulations in comparison to reference antibody crystal structures.**

Due to the computational cost associated with MD analysis, we selected five antibodies (PDB IDs: 1DLF, 1MQK, 4GXV, 5WCA, 6MEG) from the crystal structure dataset with the highest resolution, and performed the following analyses on their paired-chain structures (details in Methods section and Supp. Table 6). **(A)** Confirmation of structural convergence. We simulated the ABB-predicted and the crystal structures of the five antibodies in MD for 100,000 ps (100 ns), resulting in a total of 5001 frames per antibody for each structural origin (one frame per 20 ps). The RMSD values for each antibody from the origin structure is shown on the y-axis and the simulation time is shown on the x-axis. The blue-shaded blocks represent pre-convergent frames (0–20,000 ps), and the green-shaded blocks highlight the convergent antibody frames (from 20 ns onwards). The red horizontal lines represent the mean RMSD for convergent frames. The text on the plots represents the mean RMSD ± SD values. Only convergent frames were included in downstream analyses (panels B, C of this figure). **(B)** Pairwise RMSD measurements of MD-simulated antibody structures comparing the conformations of ABB against the conformations of crystal (reference) structures (all vs. all; 4001*4001 datapoints/antibody). Numbers shown on the figure represent the median RMSD (Å). **(C)** Overlap index ($\eta$) between the distribution of DP values when measured on the MD-simulated crystal structures and the MD-simulated ABB-predicted structures. Left panel: a graphical representation of the overlap index as defined by Pastore and Calcagnì (*196*). $\eta$ is a distribution similarity measure that can take values from 0–1 and it quantifies the overlapping area between two distribution curves. Middle panel: The overlap index measured for each antibody by DP. Each column represents an antibody and each row represents a structural DP. The color intensity in the heatmap and the numerical values within each cell reflect the overlap index. Numbers shown in brackets above each column represent the mean overlap index for each antibody. Right panel: examples of the overlap index and the value distribution of five DPs for the crystal (reference) and ABB-predicted MD-simulated convergent frames (4001 datapoints/origin) of the antibody with PDB ID 6MEG. Relates to Figure 2.

#### Feasibility analysis of inferring structures of close-distance antibody mutants

To potentially complement the sequence-based sensitivity analysis (Figure 4, Supp. Figure 12) with a structure-based one, we set out to investigate whether computational antibody structure prediction methods may be suitable for sensitivity analysis where structure-based DPs of *single* amino acid substituted antibody mutants would be compared to that of their wildtype. To this end, we studied the predicted structures of all single-amino acid substituted CDR variants of 49 sampled heavy chain sequences (10 per isotype except 9 for IgD; 30,015 mutants in total – see Methods). Of note, we performed structure prediction for this analysis using the deep learning-based tool IgFold (*55*) because template search-based tools (e.g., ABB) tend to utilize an identical structural template for antibodies that are single-amino-acid-distant from their wildtype antibody due to the high sequence similarity (*181*, *197*). We found that structural deviations induced by single amino acid substitutions in CDRs (as determined by RMSD) between wildtype and mutant were minimal (0.09–1.2 Å) with a median value of 0.31 Å (Supp. Figure 19A). This range of RMSD values is within or below the current technical resolution of experimental protein crystallography (RMSD ≈ 2Å) (*178*, *198*).

Thus, we asked whether these findings obtained by computational structure prediction are comparable with the variation observed among antibody structures determined experimentally by crystallography. To this end, we compiled a dataset of ten antibody pairs from the AbDb database (*178*) where antibodies of the same pair are one-amino-acid different and are similar in sequence length (see Methods, Supp. Table 5). Of note, the small size of this dataset is due to the limited availability of antibody crystal structure pairs that satisfy these criteria (same length and one amino acid difference in the loop).

We found that the structural variations within pairs were greater on average for experimental and IgFold predicted structures (median RMSD 0.9 and 0.7 Å, respectively – Supp. Figure 19B) in comparison to what we reported for the *in silico*-generated native antibody mutants (Supp. Figure 19A). To further investigate the difference between experimental and IgFold-predicted models, we visually inspected the structural alignment of five randomly-sampled antibody pairs from the same dataset (Supp. Figure 19C). In the case of IgFold models (shown in gold, Supp. Figure 19C), antibody pairs exhibited complete superposition, while crystal structures (shown in gray, Supp. Figure 19C) displayed subtle global variations. Nevertheless, it is challenging to solely attribute such variations to the single amino acid difference within each pair as several factors could be the origin of variance including (i) the inherent limitations and uncertainties associated with experimental variability to resolve each individual structure within the pair (*199*, *200*) and (ii) the flexibility of antibody structures, especially their CDR loops (*201*, *202*). To limit the effect of external factors (as much as possible), we focused on the exact locus of amino acid difference by computing the Euclidean distance (ED) between carbon alpha atoms ($ED_{\text{carbon alpha}}$) within each pair (Supp. Figure 19B). In both types of structures (crystal and IgFold-predicted), we reported negligible $ED_{\text{carbon alpha}}$ values (median of 0.4 (crystal) and 0.2 (IgFold) Å).

Based on our analysis, and considering the (current) shortfalls of antibody structure prediction tools and experimental variance, we hypothesize that the actual distance values among antibody structural variants, particularly at the mutation site, to be lower than experimentally-measured values and higher than computationally-predicted values. This indicates that the actual impact of single-amino-acid antibody structural variance might fall somewhere between that of the predicted computationally and that of measured experimentally. Nevertheless, as *in silico* structure-based developability prediction relies on 3D antibody structures, this analysis highlighted how current technical insufficiency hinders conducting a single-amino-acid sensitivity analysis on structure-based antibody developability parameters. For the above reasons, we limited our sensitivity analysis to sequence-based DPs (Figure 4, Supp. Figure 12).

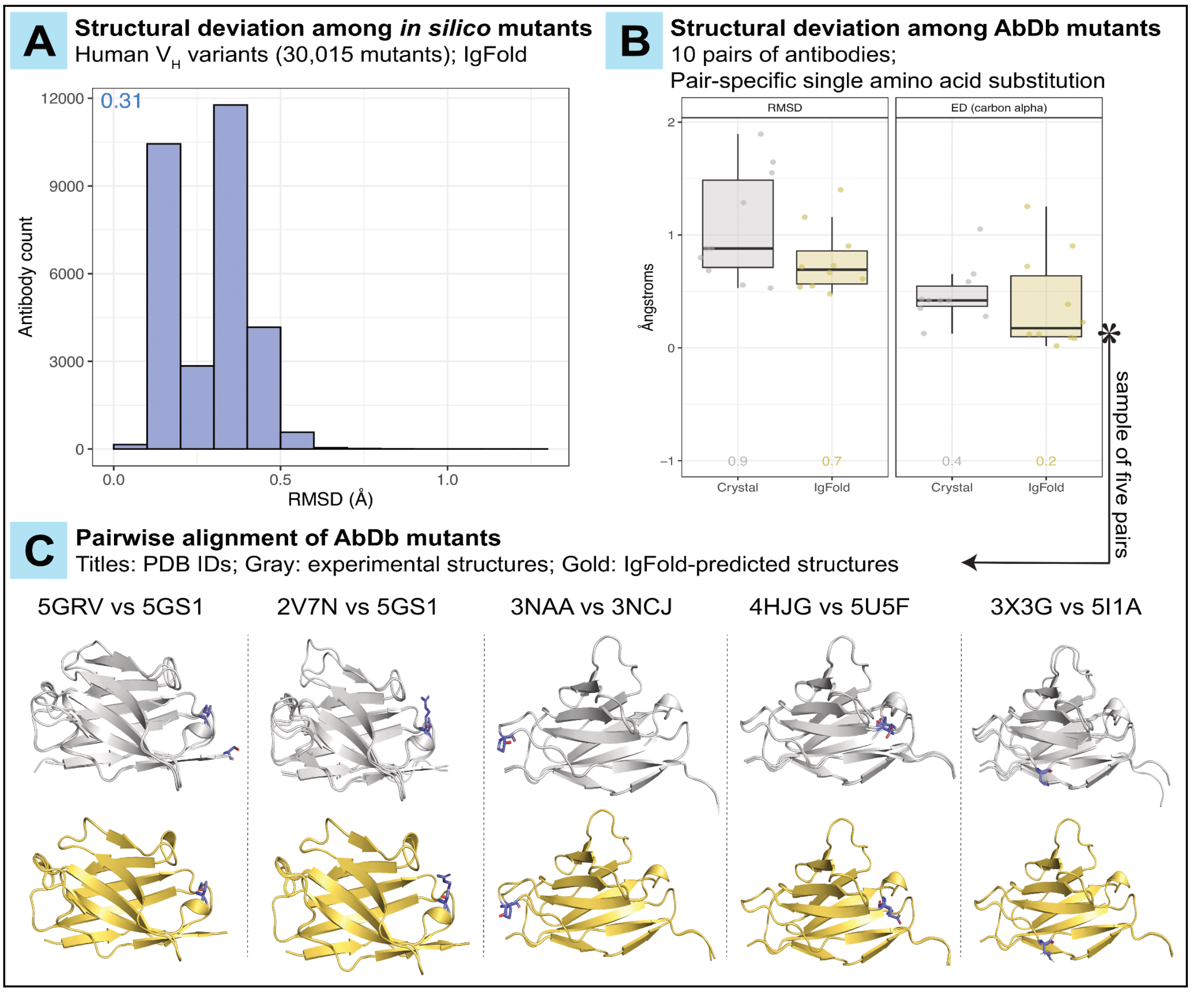

Supp. Figure 19 | **Structural-DP variance induced by single amino acids is challenging to recapitulate.** **(A)** The distribution of RMSD values among the human $V_H$ *in silico* mutants (30,015 antibodies). RMSD was measured between all CDR mutants and their corresponding WT (native) antibody after predicting their structures with IgFold (see Methods). The numerical value reported on the plot (0.31) represents the median RMSD. **(B)** The structural deviation among AbDb (*178*) antibody pairs measured with RMSD (global measure – left facet) and the euclidean distance (ED) between carbon alpha atoms (focused measure – right panel). Of note, the carbon alpha of interest refers to the first carbon atom that bears the amino acid side chain in the exact location of sequence disparity between the antibodies of each pair (Supp. Table 5). Both metrics (RMSD and $ED_{carbon\ alpha}$) were measured on crystal (experimental) structures and IgFold-predicted structures. The numerical values shown in the figure reflect the median of the corresponding metric. **(C)** Visualisation of the pairwise structural alignment of five antibody pairs (from B) shown for the crystal structures (top – gray) and IgFold-predicted structures (bottom – yellow). The side chains where the pair differ in a single mutation are drawn in all structures (represented in blue sticks). The PDB IDs of the pairs are shown on the top. Relates to Figure 4**.**
